# Grape ripening speed slowed down using natural variation

**DOI:** 10.1101/2024.08.12.607560

**Authors:** Luigi Falginella, Gabriele Magris, Simone Diego Castellarin, Gregory A. Gambetta, Mark A. Matthews, Michele Morgante, Gabriele Di Gaspero

## Abstract

Understanding ripening patterns and governing ripening speed are central aspects of grapevine (*Vitis vinifera*) berry biology owing to the importance of grape ripeness in winemaking. Despite this, the genetic control of ripening is largely unknown. Here, we report a major quantitative trait locus that controls ripening speed, expressed as speed of sugar accumulation. A haplotype originating from the species *Vitis riparia* halves maximum speed regardless of crop levels and berry sizes. The sequence of events that are normally completed at the onset of ripening in a two-week period known in viticulture as veraison are taking place at a slower speed, thereby attaining ripeness under milder weather conditions in late summer. *V. vinifera* cultivars show limited phenotypic variation for ripening speed and no selective sweep in the causal genomic region that could derive from domestication or improvement. Closely related species make up for the lack of standing variation, supplying major effect alleles for adapting grape cultivars to climate change.

**HIGHLIGHT / SIGNIFICANCE STATEMENT:** Reducing the speed of fruit ripening genetically is a means for adapting the grape berry developmental program to the changing needs of the wine industry and in response to global warming. We identified a haplotype in a wild grape species that slows down the speed of ripening in progenies of *Vitis vinifera* by limiting the speed of sugar accumulation throughout the duration of ripening, a condition of great importance for winemakers to harvest their grapes at the desired level of technological ripeness.

## INTRODUCTION

The typicity of the world’s most iconic wines is threatened by climate change^1^. Heat waves and water deficits, conditions that are increasingly impacting vineyards, accelerate grape ripening. Extreme conditions associate with negative changes in berry composition and wine sensory attributes: high alcohol concentration, low acidity and lack of balance in aroma compounds^2,3^. Vineyard management techniques that retard the premature start of the annual growth cycle in spring and postpone berry ripening in summer can mitigate accelerated ripening^4,5^. Mitigation actions ameliorate the problem, escaping seasonal peaks of air temperature and solar radiation, but do not provide an ultimate solution. Grapevine cultivars vary in phenology and in ripening speed to some extent^6^. Changing to cultivars that are better suited to new weather patterns provides one means of adaptation for viticulture. However, the standing variation in *Vitis vinifera* is limited relative to the pace of warming, casting doubts on the possibility that the benefits of changing between existing cultivars will outweigh the challenges of changing to wines with diverse varietal sensory attributes.

The grape berry is a non-climacteric fruit and follows a double sigmoidal curve of growth. Ripening starts at the end of an intermediate phase of reduced growth (lag phase)^7^ that is marked by the completion of organic acid accumulation. The inception of ripening—in viticulture referred to as veraison—takes place between 45 and 70 days after flowering (DAF) in *V. vinifera* domesticated grapevines^8,9^. This variation in timing is explained to a greater extent by genotypic differences among cultivars and somatic mutations within cultivars than by environmental factors^10–12^. A genetic determinant on Chr16 controls the duration of the interval between flowering and veraison^13,14^. A somatic mutation in a bud sport of ‘Pinot Noir’, one of the earliest ripening cultivars, anticipates veraison by a further 10-20 days^15^. Cellular, molecular and metabolic switches accompany the inception of ripening. Studies on cell turgor, berry elasticity, gene transcripts, growth regulators, sugars and organic acids have shown how these changes are ordered chronologically^16,17^, not yet how they are causally connected. Among those relevant to the experiments of this article, drops in cell turgor precede berry softening, which is reflected by lower elasticity^16^. Berry softening is operated on an individual berry basis. Berries are developmentally asynchronous on the same bunch and on the same vine. Assessment of softening-to-the-touch is a reliable, instantaneous and nondestructive indicator of the start of sugar accumulation that becomes macroscopic within hours or days using refractometer readings of juice concentration^16,18,19^.

Once activated, the ripening process is less regular and governable. The time required to complete sugar accumulation, and therefore the speed of ripening, is a puzzling metric of the ripening process. Ripening speed variance is unaccounted for by classic factors, e.g. genotype, seasonal climate variation, their interaction and experimental error^20^. Physiologically, grapevine berries operate sugar import and accumulation with outstanding efficiency^21^. They store hexoses at high concentration relative to other fruits^22^, without outperforming other species for phloem sucrose concentration^23^. Agronomically, it is difficult to impose severe restrains on sugar accumulation using vineyard management techniques that modify crop level, modulate plant water status and/or change light interception capacity. High yields nearing vine capacity, less negative water potentials and postveraison leaf removal decrease sugar accumulation, but only mildly^5^. As there is a trade-off between carbon assimilation and vine physiology balance, these interventions must be moderate in intensity and not compromise carbon storage. Otherwise, mitigating one problem in one season may result in generating negative carry-over effects in the following seasons^24^. Experimentally, growth regulators and transcription factors promote or repress diverse aspects of grape ripening^25–29^. Nonetheless, a causal relationship between one genetic factor and the speed of sugar accumulation, as a quantitative indicator of the speed of grape ripening, is not known yet.

Here, we report on an allelic variant donated by a crop wild relative that is sufficient to halve the maximum daily pace of sugar accumulation in progenies of *V. vinifera*, without regard to concomitant variation in counteracting factors and without deflecting the curve of berry growth. Variation in sugar accumulation patterns associates with variation in malate breakdown patterns suggesting that, beyond sugar metabolism, the ripening process slowed as a whole. We also show that there is little coding sequence variation between causative and alternative haplotypes, with a few non-synonymous substitutions in predicted proteins but no obvious differences in gene model structures and in insertion/deletion of transposable elements within or nearby transcriptional units. In *V. vinifera* berries, there are two opposite coexpression patterns involving multiple genes in the QTL confidence interval. Subtle disarrangements of coexpression domains is one of the possible causative factors that are compatible, genetically, with the dominant inheritance of the slow-ripening phenotype and, evolutionarily, with the presence of the slow-ripening haplotype in wild species and rootstocks.

## RESULTS

### Discovery of a ripening**-**related segregation: field observations and phenotyping

A full-sib family consisting of the two parental lines *V. vinifera* ‘Chardonnay’ (VvCHA, cultivated variety, hermaphrodite) and *V. riparia* ‘Gloire de Montpellier’ (VrGdM, undomesticated, bearing staminate flowers, commonly used in viticulture as a rootstock) and 118 F^1^ interspecific hybrids was generated in 2002. Seedlings were grown for two annual cycles. Cuttings were grafted on clonal rootstocks. After one year of adventitious root formation in nursery beds, the progeny was planted in a germplasm repository in 2006, initiated floral organogenesis in 2007 and generated inflorescences in 2008. The progeny showed a 1:1 segregation (χ^2^ = 0.14) for flower morphology (perfect flowers vs. a mixture of staminate and partially perfect flowers) and sexual dimorphism for inflorescence-related traits (**Figure 1a-c**). From 10.6 to 90 % of the phenotypic variance in secondary sex characteristics (**Figure 2**) was explained by the flower sex locus, with LOD scores ranging from 4.6 to 40.4 (**Supplementary Table S1**). Genetically male seedlings generated larger inflorescences (>433 flower buttons per panicle) than genetically hermaphrodite seedlings (<357 flower buttons per panicle) and bloomed on average two days ahead (**Figure 2**). While genetically hermaphrodite seedlings produced only perfect flowers, genetically male seedlings with morphologically male inflorescences produced a few perfect flowers with a rudimentary yet functional gynoecium (**Figure 1d-f**), thereby reproducing sexually as andromonoecious plants. They set fruit, although at a much lower rate of one berry every on average 111.1 flowers compared to hermaphrodites (one berry every 1.9 flowers), averting post-flowering abscission of one or more inflorescences per vine that developed into loose bunches (**Figure 1g-I, Supplementary Figure S1**). As a result, 94.1 % of the progeny cropped in at least one season with ample variation in crop level among individuals. Fertilized ovules promoted seed development with minor differences in seed content and viability between hermaphrodite and andromonoecious individuals (**Figure 2**).

**Figure 1.**
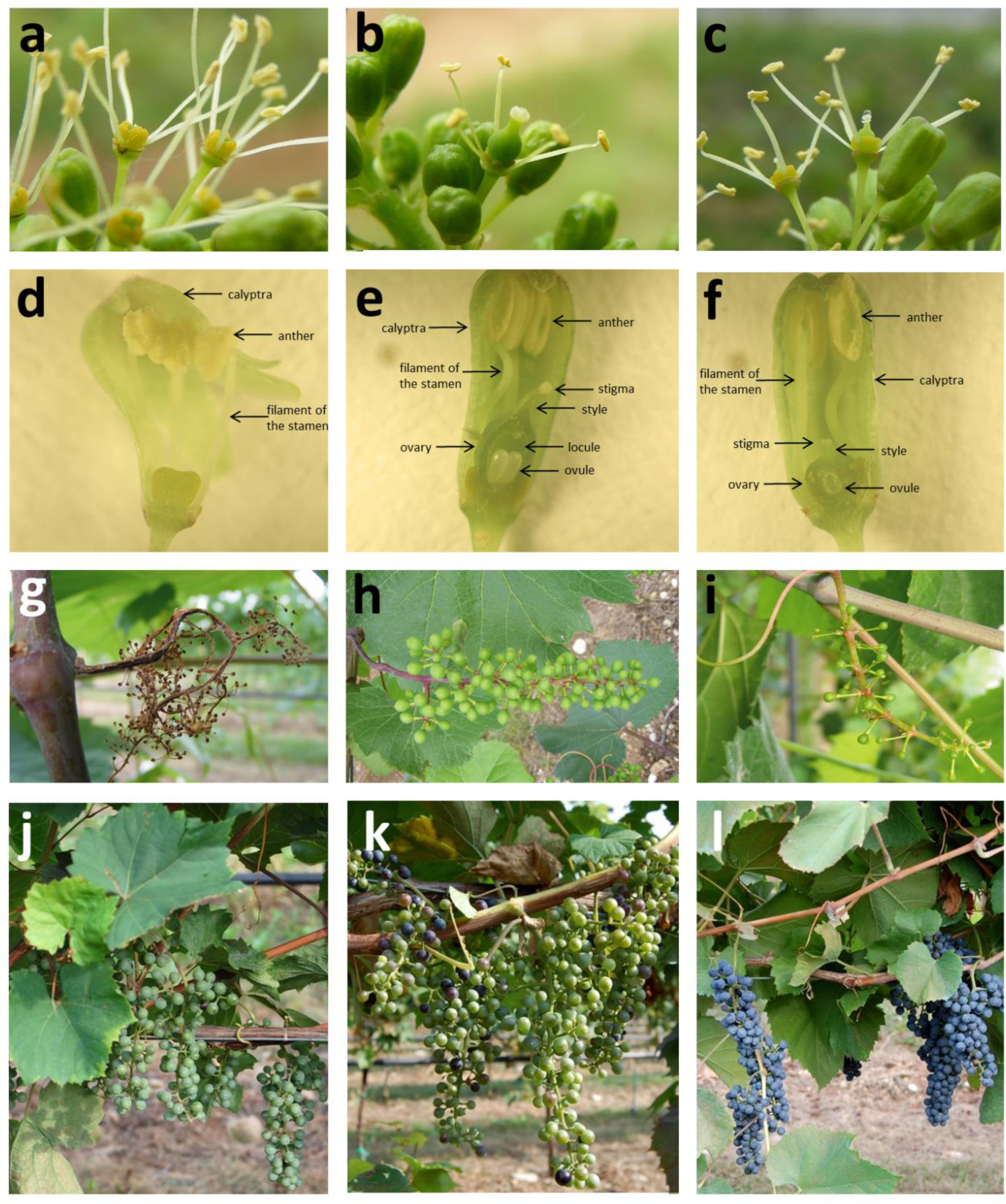
Segregation of flower types, fruit set and ripening-related traits in the VvCHA×VrGdM progeny. Flowers (**a-c**) and sections of flower buttons (**d-f**) were pictured at flowering. Withering inflorescences in the absence of any ovule fertilisation and bunches (**g-i**) were pictured at the stage of groat-sized berries. **a,d** Staminate flowers of genetically male siblings. **b** Perfect flowers of genetically hermaphrodite siblings carrying (**e**) ovaries with fully developed locules and ovules, elongated style and enlarged stigma. **c,f** Development of an ovule in rudimentary perfect flowers of genetically male siblings with functional sigma producing exudate (**c**). **h-i** Fruit set in hermaphrodites (**h**) and andromonoecious siblings (**i**). Hermaphrodites with entirely hard- and green-berried bunches (**j**), mixed (**k**), or entirely soft-and red-berried bunches (**l**) were pictured on September 8^th^, 2008, as representatives of the distribution shown in **Supplementary Figure S2**.

**Figure 2.**
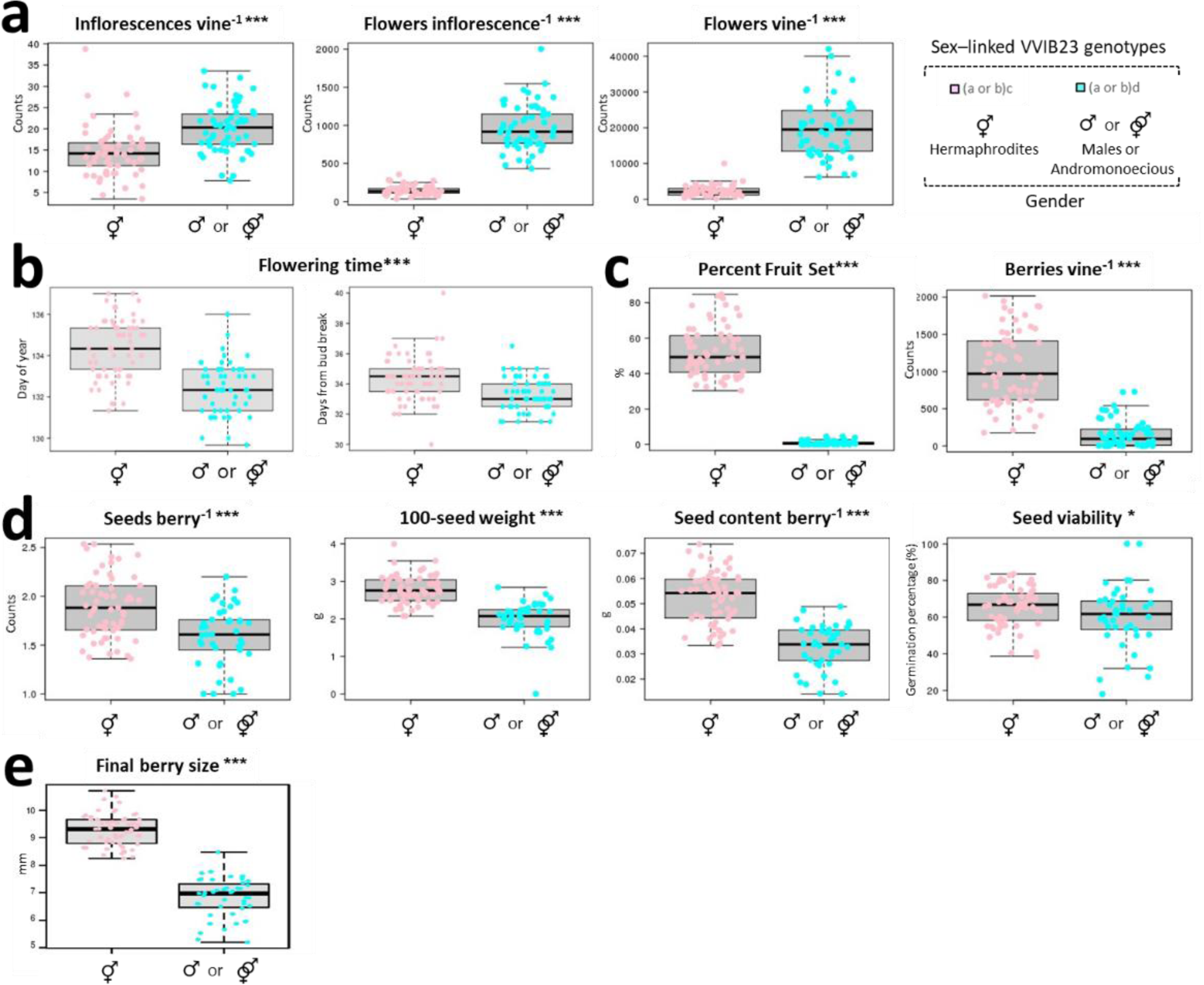
Sexual dimorphism in the VvCHA×VrGdM progeny. **a** Secondary sex characteristics at flowering. **b** Flowering time, expressed as day of year and period of time from bud break to flowering. **c** Fruit set per flower button and per vine. **d** Seed-related traits. **e** Berry size at the end of ripening. In all panels, siblings are sorted by their genotype at the sex-linked marker VVIB23. Alleles sizes are: a^VvCHA^ 284 bp, b^VvCHA^ 288 bp, c^VrGdM^ 300 bp, d^VrGdM^ 308 bp. The allele d^VrGdM^ is associated with male-type inflorescences. Values in **a-e** are averages per individual in two growing seasons. Asterisks indicate significant differences in the distributions between hermaphrodite and male genotypes using a two-sided Wilcoxon test (* <0.05, ** <0.01, ***<0.001).

At normal harvest time, periodic monitoring of soluble solids concentration showed that *V. vinifera* cultivars and breeding material in the germplasm repository ranged between 16 and 23°Brix, with the exception of the VvCHA×VrGdM progeny. Approximately a half of the VvCHA×VrGdM full-sibs showed ripening delays (**Figure 1j-k**, **Supplementary Figure S2**). The progeny did not segregate for berry skin colour and all siblings accumulated anthocyanins in the epicarp when soluble solids concentration surpassed 12°Brix (**Supplementary Figure S3**). Following these observations, we phenotyped the progeny for berry growth and ripening curves across two consecutive annual growth cycles (2009 and 2010 years with contrasting seasonal weather patterns, **Supplementary Figure S4**) for QTL analysis of ripening-related indicators and yield-related parameters. We used a subset of 30 hermaphrodites for quantifying differences in seed content and seed-to-berry ratio between ripening classes. We confirmed the stability of the QTL in the 2011 season and excluded carryover effects of genotype-dependent crop level variation on ripening speed for three seasons in a row (**Supplementary Figure S5**). In the 2012 season, we monitored single-berry growth and quantified sugars and organic acids in representatives of each ripening class. In the 2012 and 2013 seasons, ripening curves were determined in additional siblings to support QTL location.

### Phenotypic and genetic variation in soluble solids accumulation and berry growth

The curves of soluble solids accumulation showed ripening patterns in half of the VvCHA×VrGdM progeny that are similar to those observed in grapes of the seed parent VvCHA, typical of *V. vinifera* cultivars, increasing from pre-veraison basal levels to 16°Brix in approximately 3 weeks (**Figure 3**). These segregants showed fast-ripening phenotypes, hereafter also referred to as normotypes. The other half of the progeny progressed into ripening following flattened curves, uncommon in *V. vinifera* cultivars, with a gradual and continuous increase in soluble solids concentration over an extended period of time, representing slow-ripening phenotypes. According to a time-course QTL analysis for soluble solids concentration at various ripening stages (**Figure 3f** and **Supplementary Figure S6)**, a single locus on VrGdM Chr6, hereafter referred to as *RIPENING SPEED*, explained up to 83.0 % and 86.7 % of the phenotypic variance in 2009 and 2010, respectively. The explained variance was confirmed at 85.7 % in 2011 despite interannual variability in weather conditions. Consistently among years, soluble solids concentration in the first decade of August (DOY222-223, **Figure 3f inset** and **Supplementary Figure S6a**), approximately 80-90 DAF and a few days around the exact date when heat accumulation surpassed 1200 GDD^C^ (**Supplementary Figure S6b-c**), showed the highest percentage of the phenotypic variance explained by the *RIPENING SPEED* QTL. Several ripening-related indicators correlated with soluble solids accumulation and with the *RIPENING SPEED* locus (**Supplementary Figure S5** and **Supplementary Table S2**). The most notable ones are the day of year (DOY) on which each seedling surpassed 16°Brix and the interval of time since flowering that was required to surpass 16°Brix. When we sorted VvCHA×VrGdM siblings according to their *RIPENING SPEED* genotype, the average curve of soluble solids accumulation differed remarkably between slow-ripening and fast-ripening classes (**Figure 3d-e inset** and **Supplementary Figure S7c-d**). The daily pace of sugar accumulation reached the maximum 3 to 4 weeks after the resumption of berry growth in fast-ripening genotypes. A lower relative maximum was reached 7 to 8 weeks later in slow-ripening genotypes (**Figure 4a-d**). Differences in sugar accumulation between ripening classes were insensitive to variation in crop level from nearly 0 to >2,000 berries vine^-1^, in timing of fruit set from 1 to 9 days (in **Supplementary Figure S7** date of flowering is a proxy for ovule fertilization) and in final berry size from 6 to 11 mm attained at maturity (**Supplementary Figure S7**). This variation was genetically determined within each ripening class by the independent segregation of flower morphology (see below). We therefore exclude that these physiological factors, which are known to inversely correlate with soluble solids^30^, explained *RIPENING SPEED*.

**Figure 3.**
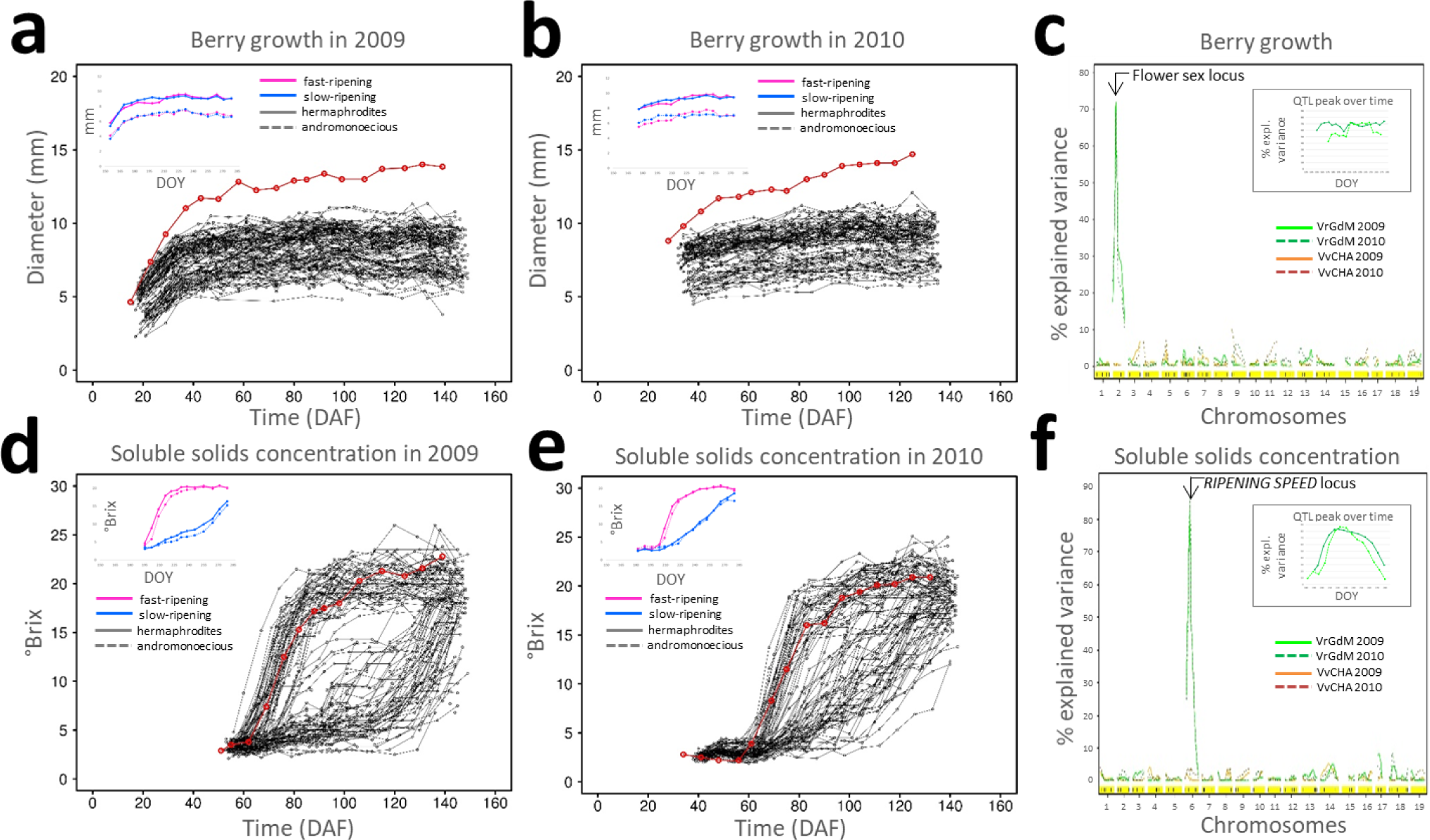
Biennial curves of berry growth and soluble solids accumulation in the VvCHA×VrGdM progeny and genetic basis of phenotypic variation. Data in **a-b** and **d-e** are referred on the x-axis to days after flowering (DAF) in each individual. Insets in **a-b** and **d-e** show average values per genotypic class (**c** and **Supplementary Figure S7**). Data in insets are referred on the x-axis to the day of year (DOY) of monitoring in order to express average values per genotypic class. QTL plots of explained phenotypic variance for berry growth (**c)** and soluble solids concentration (**f**) at the time point of the highest explained variance during berry growth and ripening (**insets** and **Supplementary Figure S6a-c**). Explained variance by VvCHA alleles is shown with orange lines and by VrGdM alleles with green lines and it is referred on the x-axis to chromosomal coordinates of the markers. Solid lines represent data from the 2009 season, dashed lines from the 2010 season.

**Figure 4.**
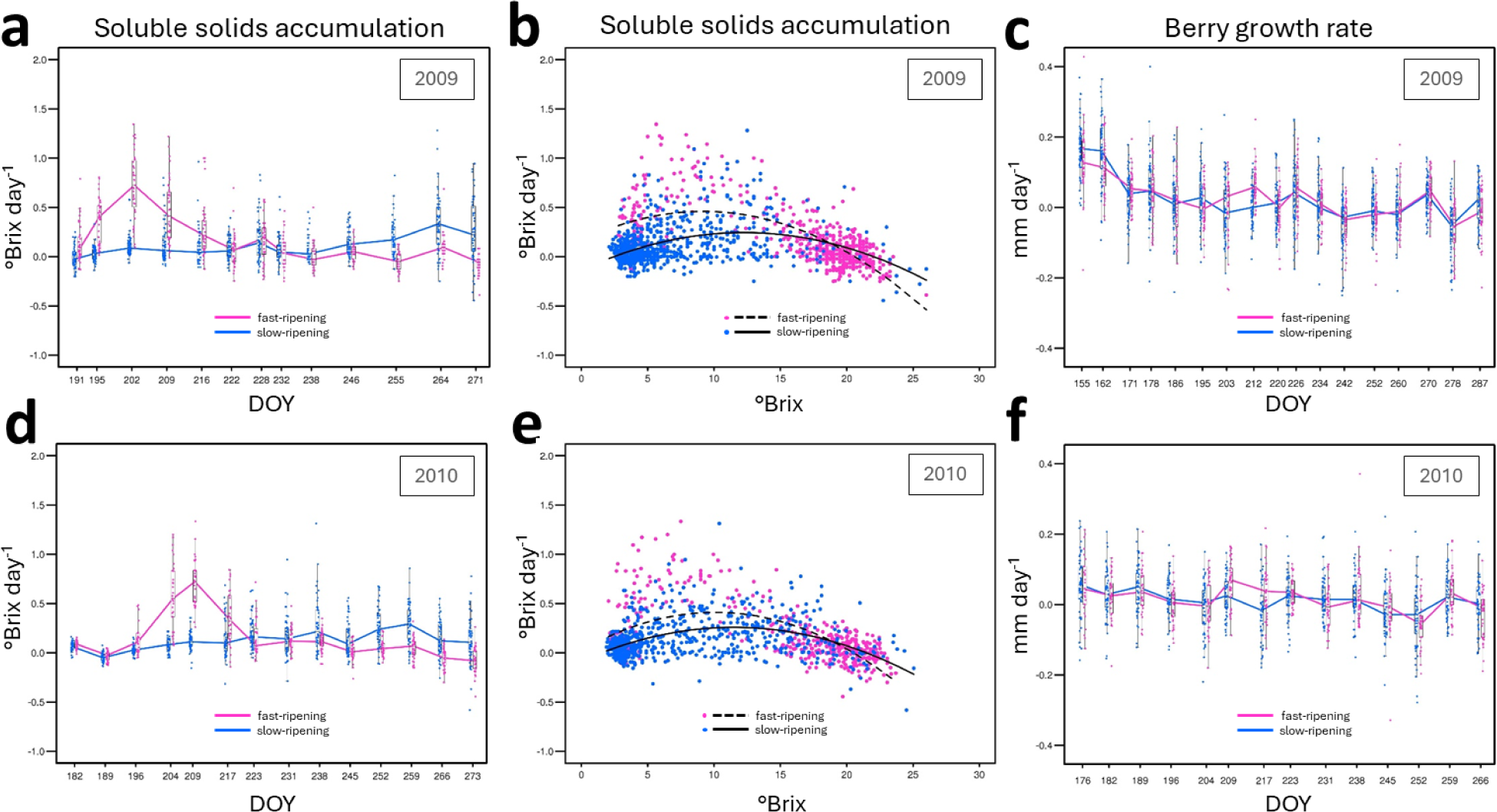
Biennial pace of soluble solids accumulation and berry growth. Dots represent values per individual in each genotypic class. Lines represent averages (**a**, **c**, **d** and **f**) and regression curves (**b,e**) per genotypic class. Daily averages of nearly weekly intervals are plotted on the initial point of each interval. °Brix day^-1 i^n **a** and **d** are referred on the x-axis to the day of year (DOY) of monitoring. mm day^-1 i^n **c** and **f** are referred on the x-axis to the ripening stage at the moment of monitoring in each individual. Graphs in **b** and **e** are provided in **Supplementary Figure S8** with fitting regression models per genotypic class.

The curves of berry growth (**Figure 3a-b**) did not reflect the differences shown by the patterns of soluble solids accumulation (**Figure 3d-e**), suggesting that post-veraison berry growth was uncoupled from soluble solids accumulation in slow-ripening genotypes. When we sorted VvCHA×VrGdM siblings according to their *RIPENING SPEED* genotype, the average curve of berry growth overlapped between genotypic classes (**Figure 3a-b inset** and **Supplementary Figure S7a-b**). Differences among VvCHA×VrGdM siblings in berry growth patterns and in final berry size were due to an independent segregation for hermaphrodite and andromonoecious phenotypes (**Supplementary Figure S7a-b)** and were largely explained by the flower sex locus (**Figure 3c** and **Supplementary Figure S6d-f**). At the beginning of the second phase of berry growth, slow-ripening genotypes showed a transiently lower pace of berry growth in comparison to fast-ripening genotypes despite a far lower pace of sugar accumulation. This difference occurred within a narrow window of time of 10 days in 2009 (DOY203-212) and 9 days in 2010 (DOY209-217). In order to express the pace of soluble solids accumulation in relation to the stage of progression of soluble solids accumulation, we arranged rates in a °Brix sequence (**Figure 4c,f** and **Supplementary Figure S8**). Second-order polynomial regression curves indicate that fast-ripening is operated with the highest accumulation rate (0.41-0.46°Brix day^-1^) occurring at the stage of 9.4-10.0°Brix. Slow-ripening is operated with a lower maximum rate of 0.24-0.26°Brix day^-1^, which is reached later on during the progress of ripening at the stage of 11.5-12.2°Brix. Regression curves also suggested that soluble solids accumulation is ceased at approximately 20°Brix in both classes regardless of their differences in ripening speed.

### Physiological aspects of slow-ripening

Slow-ripening associates with small yet distinctive changes in berry development that were measurable before and after the onset of ripening (**Supplementary Figures S9-S10**). When ontogenetic drift and size-dependent effects are discounted by plotting sugar content per berry versus berry fresh mass (**Figure 5a,c**), following the allometric analysis proposed by Sadras and McCarthy^31^, fast- and slow-ripening patterns were fitted to two different polynomial models. The fast-ripening pattern followed a sigmoidal curve that is best approximated by a four-parameter log-logistic model (**Figure 5c**, Spearman correlation = 0.9176, Pearson correlation = 0.8863). In contrast, the slow-ripening pattern followed an exponential trend that is best approximated by a power function (**Figure 5c**, Spearman correlation = 0.8177, Pearson correlation = 0.8509). The growth of individual berries showed that a rapid glucose and fructose accumulation on a per berry basis occurs suddenly in berries exiting the lag-phase in fast-ripening genotypes (**Figure 5c** and **Supplementary Figure S9**). In slow-ripening genotypes, berries continued to grow and to accumulate organic acids, in particular malate, while the fast-ripening berries were stalling in their lag-phase. As the slow-ripening genotypes started to accumulate glucose and fructose, malate breakdown also slowed, without plateauing until the end of our monitoring period (**Supplementary Figure S9f,h,j**). These data therefore suggest that the slow-ripening phenotype is either partially or entirely caused by physiological conditions that are established prior to the onset of sugar accumulation and do not change as sugar accumulation takes place or by a long-lasting cascade of consequences triggered by earlier events.

**Figure 5.**
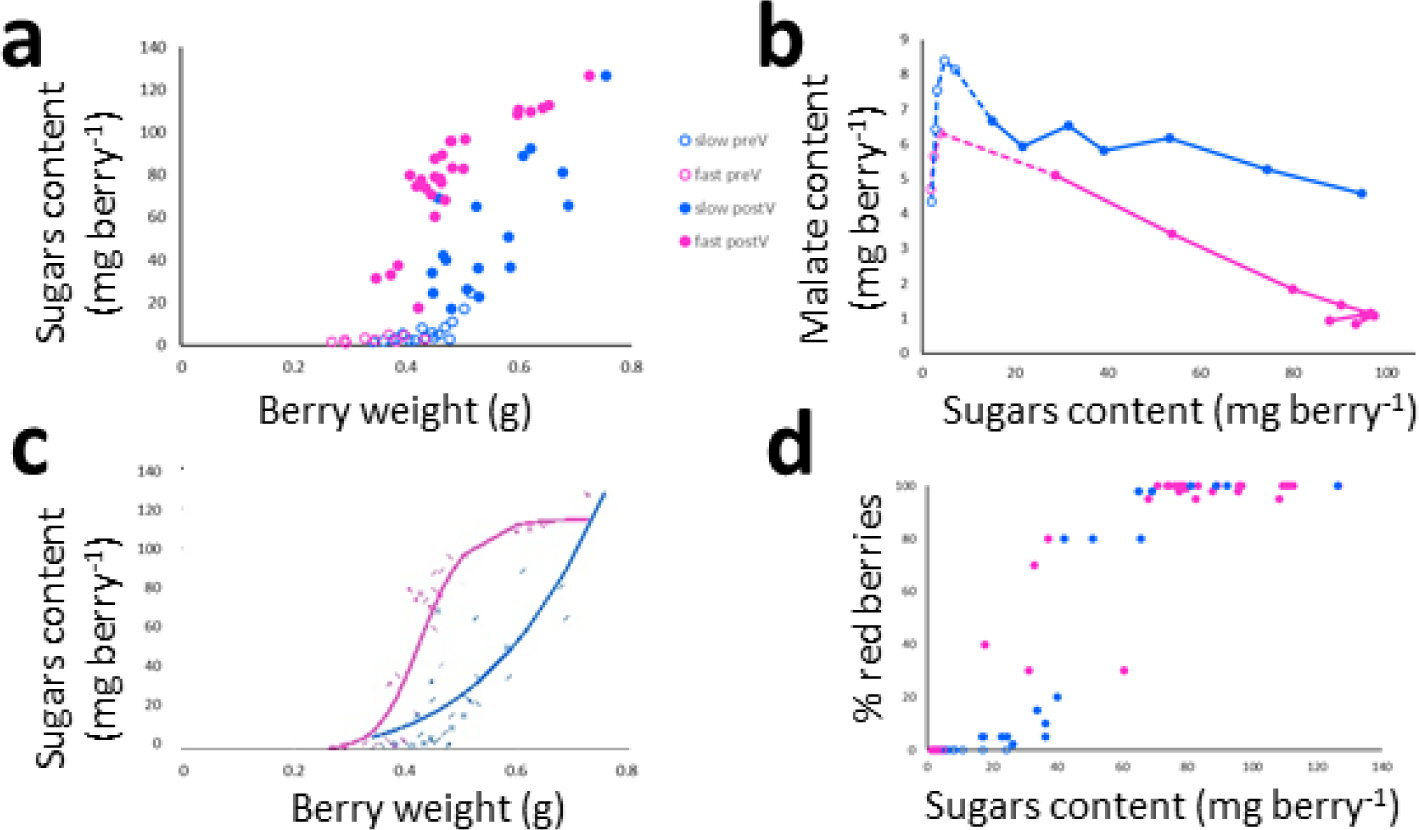
Relations between sugars and berry weight, malate and sugars, colour transition and sugars in slow-ripening and fast-ripening siblings. Dots in **a**, **c** and **d** represent values in three individuals per genotypic class. Dots in **b** represent averages among individuals per genotypic class. Lines in **b** connect consecutive stages. Pre-veraison (preV) refers to stages with green berried bunches (empty dots). Post-veraison (postV) refer to stages in which colour transition is initiated (solid dots). Lines in **c** represent log-logistic (magenta) and power (blue) fit curves for fast-ripening and slow-ripening patterns, respectively.

One of these conditions may originate in the fact the *RIPENING SPEED* QTL also accounted for higher berry weight in slow-ripening seedlings during the phase of green berry growth (**Supplementary Figure S10**) and longer berry growth that was followed, without discontinuity in time, by the onset of soluble solids accumulation (**Supplementary Figure S9a**). One possible factor for setting this condition early in berry development and for operating a conditional effect throughout ripening is the total seed weight per berry (hereafter referred to as seed content). Seed content is largely determined during the phase of green berry growth and is known to influence both growth^32,33^ and ripening^34,35^. The 100-seed weight and seed content at berry maturity (**Figure 2**) were chiefly controlled in the VvCHA×VrGdM progeny by the flower sex locus (**Supplementary Table S1**). However, a residual part of the observed variation was controlled by the *RIPENING SPEED* QTL in the absence of significant variation for seed number between slow-ripening and fast-ripening genotypes (**Supplementary Table S2**). In order to clarify whether genetically-determined variation in seed growth is antecedent or subsequent to the establishment of the differences in sugar accumulation patterns, we determined seed and berry weight in developing and ripening berries (**Figure 6**). We found that 100-seed weight and seed content per berry are higher in slow-ripening genotypes during the phase of green berry growth. Berry weight is higher as well. Average seed-to-berry ratio is slightly yet consistently higher over time. This suggests that heightened berry weight in slow-ripening genotypes at the end of green berry growth might be partially due to higher seed weight and partially mediated by higher seed weight. In normotypes, transition berries that initiate ripening are characterized by lower seed-to-berry ratios than asynchronous lagging berries that coexist on the same bunch. As a result, low seed-to-berry ratios are considered a physiological factor that promotes anticipated ripening in the absence of genetic variation among berries^36^. We also observed sharp drops in seed-to-berry ratio in fast-ripening siblings and in ‘Chardonnay’ concomitant with a rapid pericarp growth driven by sugar accumulation, thereby representing normotypes. The slow-ripening siblings showed, instead, a continued linear decrease in seed-to-berry ratio that is nearly insensitive to the onset and progression of ripening (**Figure 6**).

**Figure 6.**
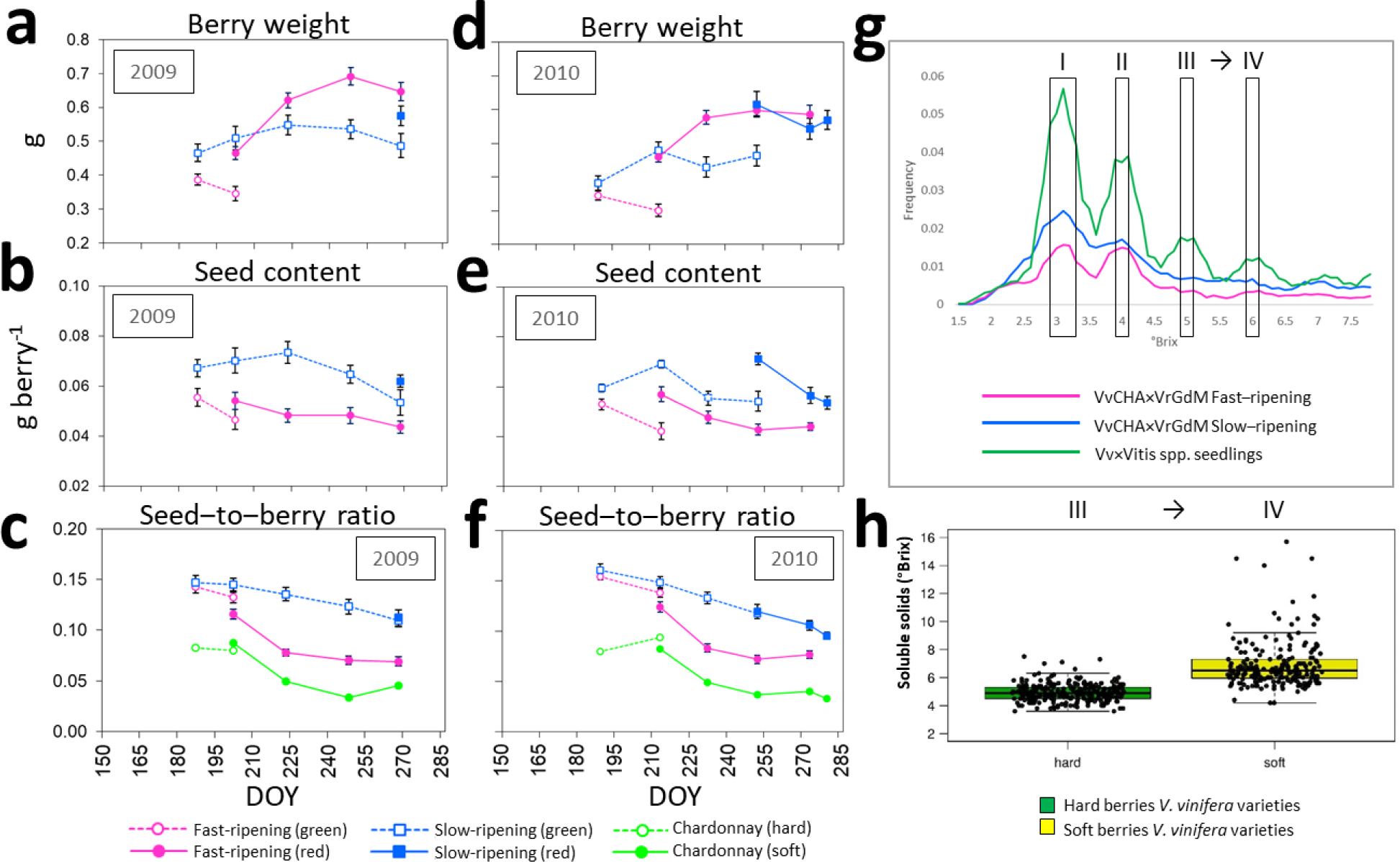
Relations between berry weight and seed content before and after the onset of ripening. **a-f** Slow-ripening and fast-ripening siblings. Green berries were collected before the onset of ripening. Red berries were collected after the onset of ripening. Green and red berry stages correspond to hard and soft berry stages in the control ‘Chardonnay’. At the stage of colour transition, green and red berries were collected on the same day from the same bunch. Bars represent standard errors. **g-h** Onset of ripening based on °Brix frequency spectrum from daily monitoring of hybrid progenies (**g**) and °Brix in hard and soft berries of *V. vinifera* cultivars (**h**). °Brix were taken with a 0.2 resolution. The graph in **g** is based on *n*=5,447 °Brix measurement in two seasons in fast-ripening siblings, *n*=3,781 measurements in slow-ripening siblings and *n*=13,860 measurements in 295 siblings of two *Vitis vinifera* × *Vitis* spp. crosses (VvCs×Vspp20/3 and VvCHA×VsppBIA) described in^63^ and grown at the same site. Box plots in **h** are based on measurements in 81 genotypes of a *V. vinifera* diversity panel (data obtained from^42^). Hard and soft berries were collected on the same day from the same bunch.

Physiologically, slow-ripening is associated with five anomalies that transcend the known range of variation in the crop species. First, a lag-phase of reduced berry growth is not manifest in slow-ripening genotypes. We monitored the patterns of berry growth and ripening in eight *V. vinifera* cultivars and two bud sports that span the widest known variation for timing and duration of the ripening period. A lag-phase was evident in *V. vinifera* regardless of among-genotypes differences in phenology (**Supplementary Figures S11-S12)**. Second, a sudden increase in soluble solids concentration concomitant with berry softening— corresponding to the third leap in chronological order in the stepwise pattern of soluble solids accumulation in *V. vinifera* cultivars (**Figure 6h**), in their progeny (**Figure 6g**) and in fast-ripening genotypes (**Figure 6g**)—is lagged or suppressed in slow-ripening genotypes (**Figure 6g**). We monitored soluble solids concentration at short regular intervals (i.e. every other day) at the onset of ripening on thousands of genotypes and berries (**Figures 3** and **6**). Hard berries exiting the lag phase in normotypes were associated with refractometric readings ranging from 4.8 to 5.2°Brix and softening berries were associated with readings higher than 5.8°Brix, while intermediate °Brix readings were rarely detected, indicating that a 5.2 to 5.8°Brix transition occurs rapidly. This rapid soluble solids increase is smoothed in slow-ripening genotypes, resulting into the third anomaly: a reduced ripening speed. Sugars-to-(total)acids (S/TA) ratio in *V. vinifera* evolves as a linear function of thermal summation for some weeks following the onset of ripening. The slope of this relation (α) is a proxy for the ripening speed and -β/α is a proxy for the timing of mid-veraison^6^. Fast- and slow-ripening genotypes differ by less than one day in their predicted mid-veraison time (Δ-β/α = ∼12 GDD). Then, α is 13-fold lower in slow-ripening phenotypes (0.349) than in fast-ripening phenotypes (4.6027) and ‘Chardonnay’ (4.4462) for 7 weeks. In slow-ripening phenotypes, this period is followed by another period wherein the (S/TA)/GDD relationship remains linear with a steeper slope (α=1.6359), yet lower than the maximum values in normotypes, for 4 weeks until the first decade of October (**Figure 5d**). The fourth anomaly is a slow colour transition in reddening bunches of slow-ripening genotypes. Colour transition is completed in 8-19 days in fast-ripening genotypes. In slow-ripening genotypes, colour transition starts in bunches that show an average juice hexoses concentration comprised between 21.6 and 43 g L^-1 a^nd it is nearly accomplished (99.3 %) in bunches with an average juice hexoses concentration of 201 g L^-1^, taking 36-61 days to complete (**Supplementary Figure S9**). The fifth anomaly is a persistently high seed-to-berry ratio in slow-ripening genotypes (**Figure 6c,f**). While there is substantial variation in preveraison seed-to-berry ratios among *V. vinifera* cultivars and some cultivars show similar values to those shown by slow-ripening genotypes, precipitous drops of seed-to-berry ratio occur invariably at veraison in *V. vinifera* and never occur in slow-ripening genotypes (**Supplementary Figure S11**).

### Genetic control of slow-ripening

The *RIPENING SPEED* QTL lies on the upper arm of Chr6 in an active chromatin compartment with high transcriptional activity (**Supplementary Figures S13-S14**). The segregating SNP alleles of VrGdM that are associated with the slow-ripening phenotype in the progeny are phased in the 349.RGM.pseudoHap2.1 scaffold of the VrGdM genome assembly^37^, hereafter referred to as slow-ripening haplotype. The alternative SNP alleles are phased in the 349.RGM.pseudoHap2.2 scaffold, hereafter referred to as fast-ripening haplotype (**Figure 7**). Another rootstock, Kober 5BB, an offspring of VrGdM and *Vitis berlandieri* that was selected at the dawn of the past century (**Supplementary Figures S15-S16**) and for which a nearly gapless, haplotype-resolved assembly has been recently generated using PacBio HiFi reads^38^, shares the slow-ripening haplotype with VrGdM (**Supplementary Figure S17**). We resequenced a slow-ripening VvCHA×VrGdM offspring (UD-53,105) to confirm the genealogical relationships and exclude the occurrence of recent mutations between the biological parent of the VvCHA×VrGdM progeny and the sequenced specimens of VrGdM^37^ and ‘Kober 5BB’^38–40^ available from public databases. IBD analysis confirmed a parent-offspring relationship between VrGdM and UD-53,105 and a half-sibling relationship between UD-53,105 and ‘Kober 5BB’ **(Supplementary Figures S18-S19**), which have inherited alternative homologs from VrGdM across 17.1 % of their *V. riparia* haploid genome. UD-53,105 and ‘Kober 5BB’ have therefore a coefficient of relationship of 32.9 %, against an expected 25 % for full-siblings based on random assortment. The sequence of the slow-ripening haplotype, shared by descent by VrGdM, UD-53,105, and Kober 5BB, is identical across the QTL confidence interval in the sequenced specimens (**Supplementary Figures S18-S21**, Integrative Genome Viewer^41^).

**Figure 7.**
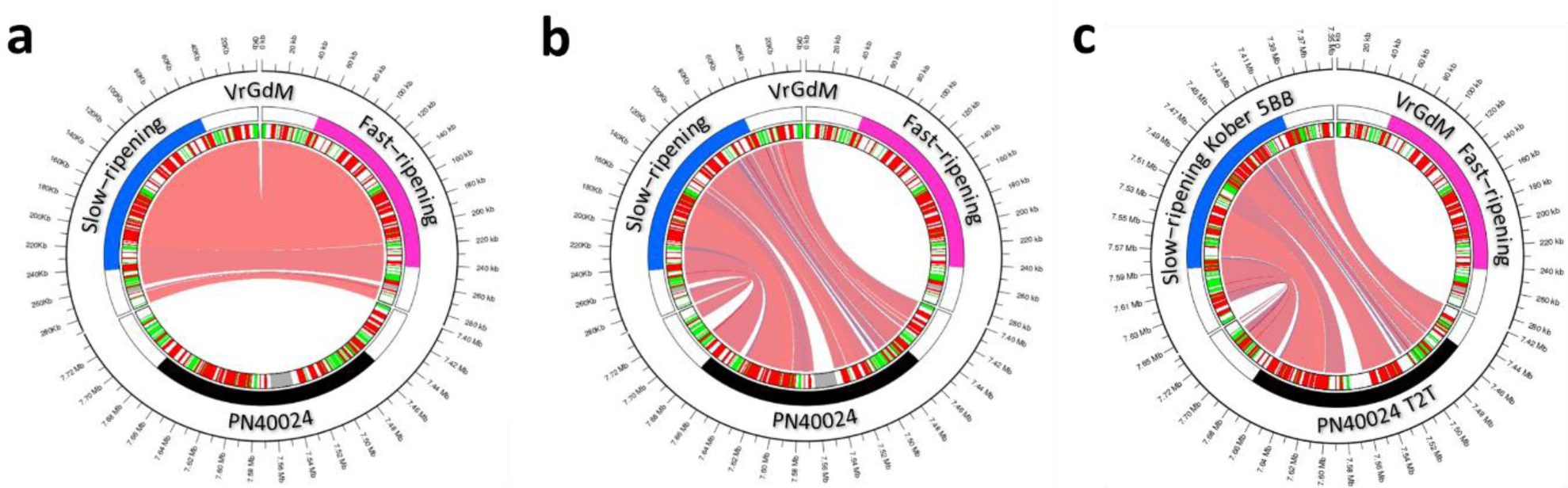
Synteny between haplotypes in the *RIPENING SPEED* QTL. Circos show comparison between the slow-ripening and fast-ripening haplotypes assembled from VrGdM (**a**), between the slow-ripening haplotype from VrGdM and the *V. vinifera* ‘PN40024’ 12Xv0 assembly (**b**), and between the gapless assemblies of the slow-ripening haplotype from ‘Kober 5BB’ and ‘PN40024’. The rulers indicate chromosomal coordinates in Mb. The outer circle shows the *RIPENING SPEED* QTL (coloured). The inner circle shows genes (red) and repetitive DNA (green). Ribbons connect intervals of shared DNA. Details of the comparisons between haplotypes are provided in **Supplementary Figures S21-S23** and **S25.**

Four recombinant chromosomes between VrGdM homologs on each side of the QTL peak support a confidence interval of ∼1.2 Mbp for the region containing the causal factor of the slow-ripening phenotype (**Supplementary Figure S13**). The QTL peak spans a physical size of 188-225 Kb in different genomes and assemblies (**Supplementary Table S3**). It corresponds to a sequence of 187.7 Kb in the gapless slow-ripening haplotype assembled from ‘Kober 5BB’, consisting of 16.7 % of CDS, 37.2 % of introns and UTRs, 11.7 % of intergenic repetitive DNA and 34.4 % of low-copy intergenic DNA. It contains 24 predicted gene models and 5 actively transcribed regions without open reading frames or similarity of their translated nucleotide sequence with plant proteins, which we considered non-coding RNA (ncRNA). All gene models and ncRNA loci are collinear with those found in the fast-ripening haplotype (**Supplementary Figures S21-S22**). Based on alignments of VrGdM short reads with the gapless assembly of the slow-ripening haplotype, we called 619 SNPs and small indels between the slow-ripening and the fast-ripening haplotype. Of these, 60 reside in coding sequences and are present in 19 genes, accounting for 18 nonsynonymous substitutions, one inframe indel and 41 synonymous substitutions (**Supplementary Tables S4-S5**). In 9 polymorphic nonsynonymous sites, which are located in 7 genes, the slow-ripening haplotype of VrGdM carries a variant allele with respect to the normotype ‘Chardonnay’. Another 132 SNPs and small indels lie in introns, 43 in UTRs and 384 in intergenic space. Counts of RNA-Seq reads spanning exonic SNPs from gene expression data in VrGdM leaves and roots excluded that sequence variants lead to allele-specific transcriptional inactivation and nonsense-mediated RNA decay in the slow-ripening haplotype compared to the fast-ripening haplotype of any gene and ncRNA wherein this analysis was informative (**Supplementary Table S6**). There is no evidence of chromosomal rearrangements between haplotypes in the QTL peak. There is no LTR retrotransposon. There are only two small DNA transposons. They are present in the fast-ripening haplotype and absent from the slow-ripening haplotype. The first element consists of a 428-bp sequence, including 78/79-bp long terminal inverted repeats (TIR) flanked by a 3-bp target site duplication (**Supplementary Figure S22**). This element has invaded the intergenic space downstream of the 5’UTR of *Vri.chr06.ver1.0.g079260* (- stranded, corresponding to *V. vinifera VIT_206s0004g06760*, ∼450 bp away from the TSS) and upstream of the 3’UTR of *Vri.chr06.ver1.0.g079270* (+ stranded, corresponding to *V. vinifera VIT_206s0004g06770*). The insertional event has putatively occurred in a common ancestor of *V. riparia* and closely related species (**Supplementary Figure S22**). As the ancestral state is shared by the VrGdM slow-ripening haplotype and *V. vinifera* haplotypes, it is unlikely that the absence of this DNA transposon alone could explain the slow-ripening phenotype. The second element is a Harbinger transposon consisting of a 280-bp sequence that has invaded the 5’ intergenic region of the gene *Vri.chr06.ver1.0.g079450*, which encodes a RING-H2 finger E3 ubiquitin ligase, ∼300 bp upstream of the TSS (**Supplementary Figure S22**). There is localized insertion/deletion variation in the 5’ region of *Vri.chr06.ver1.0.g079450*. The fast-ripening haplotype has an 18-bp and a 3-bp insertion a few bases upstream of the Harbinger insertion. *V. vinifera* does not have the Harbinger element, the insertional site lies in the middle of a 529-bp deletion and it has a downstream insertion of a 7.8-Kb LTR retrotransposon (**Supplementary Figure S23**). Both flanking protein-coding sequences at the edges of this intergenic regions show relatively high levels of antisense transcription in VrGdM^41^. There is also localized and small insertion/deletion variation between the segregating VrGdM haplotypes in the very close vicinity of the 3’UTR of *Vri.chr06.ver1.0.g079350*, which encodes a protein phosphatase 2C. Notably, the slow-ripening haplotype has a private 64-bp deletion, compared to the fast-ripening haplotype and *V. vinifera* haplotypes, within a ∼30-bp distance from the 3’ most distal evidence of transcription and another 40-bp deletion nearby.

The QTL peak region shows an irregular pattern of gene density (**Supplementary Figure S24**), despite being very poor in transposable elements. Some genes are widely spaced with ample intergenic regions. Other genes are narrowly spaced and some of pairs are arranged in a head-to-tail orientation with a small 5’ region in one of the members of the pair. Other pairs are arranged in a tail-to-tail orientation and their transcriptional units are separated by a few nucleotides. Stranded RNA-Seq libraries suggest that they potentially generate transcripts from opposite DNA strands with overlapping 3’UTRs (see Integrative Genome Viewer^41^). ncRNA is also transcribed from intergenic regions of the QTL peak (**Supplementary Table S4**) in both VrGdM and *V. vinifera*. In one case (LOC117916006), ncRNA is transcribed from downstream of the 3’UTR of a gene from the opposite strand relative to the neighboring gene *Vri.chr06.ver1.0.g079420*, which encodes a Chloroplast Outer Envelope Protein (CHUP1) or INCREASED PETAL GROWTH ANISOTROPY 1 (IPGA1) protein, thereby generating partially overlapping reverse complement transcripts (**Supplementary Figures S25-S27**). In another case (LOC117917190), a small nucleolar RNA (snoR136) shows differences in transcriptional levels between ripening berries of two ‘Pinot Noir’ isogenic lines that differ for ripening patterns (**Supplementary Figure S28a**). There is also evidence that snoR136 transcripts are processed into small RNAs in the size range comprised between 20 and 25 bp in ‘Cabernet Sauvignon’ and ‘Sangiovese’ berries (**Supplementary Figure S28b**).

### Positional and functional candidate genes

As *V. vinifera* show little variation for ripening speed, we scanned the QTL region for selection signals in a panel of cultivated genotypes^42^. Haplotype diversity, Tajima’s D and relative nucleotide diversity with respect to the wild progenitor (π_cult_/ π_wild_) did not show local deviations from chromosome-wide patterns that could derive from population bottlenecks (e.g. domestication, improvement or breeding, **Supplementary Figure S29**). However, in developing and ripening berries of domesticated *V. vinifera*, we detected local coexpression modules that involve genes in the QTL peak and in QTL the confidence interval (**Supplementary Figure S30**). Seven genes within the QTL peak, four of which are possibly associated with endogenous hormone levels, sensing, signal transduction and response and two of which are possibly involved in cell wall metabolism, showed distinctive expression patterns in relation to various berry developmental phases. Two of these positional candidate genes attracted our attention because they showed the tightest relation, though in opposite direction, between expression levels and the time course of berry ripening in *V. vinifera* (**Supplementary Figure S31**). The expression of *VIT_206s0004g06790*, which encodes GA2-oxidases that are reported to deactivate bioactive gibberellins^43,44^, is high during green berry growth and during the lag phase. It is then suppressed at softening and throughout ripening in ‘Pinot Noir’ and ‘Cabernet Sauvignon’. This pattern is consistent among varieties^45^ (**Supplementary Figures S32**) and it is mirrored by the expression pattern of *VIT_206s0004g06950*, which is located ∼150 Kb away and encodes a protein with the highest similarity with HOOKLESS1 (HLS1). HLS1 regulates responses to ABA^46^ and ethylene^47^ and plays a negative role in sugar and auxin signaling^48^. *VIT_206s0004g06910*, which encodes a putative microtubule-associated protein similar to Chloroplast Outer Envelope Protein (CHUP1) or INCREASED PETAL GROWTH ANISOTROPY 1 (IPGA1), and *VIT_206s0004g06820*, which encodes a pectin acetyltransferase PMR5, also show a decreasing expression trend. IPGA1 influences cell growth by guiding cellulose deposition in the cell wall^49^. PMR5 alters physical and mechanical properties of cell walls by modifying cell wall pectins^50^. Other genes in the QTL confidence interval that may have functional implications in berry ripening show similar trends, notably *ethylene-overproduction protein 1* (*eto1*, a negative regulator of ethylene biosynthesis), farnesol kinase (a negative regulator of abscisic acid signaling) and sugar phosphate/phosphate translocator. On the contrary, the expression of *VIT_206s0004g06770* increases sharply at softening in ‘Pinot Noir’ and ‘Cabernet Sauvignon’. We investigated gene expression differences that are associated in ‘Pinot Noir’ with differences in ripening patterns^15^ and in ‘Merlot’ with physiological variation in berry size^51^. The upregulation of *VIT_206s0004g06770* is earlier in timing and stronger in intensity in berries of the early-ripening mutant of ‘Pinot Noir’ than in berries of the wild type, while the expression is not different between berry size classes in ‘Merlot’ (**Supplementary Figure S31**). In a wider set of genetically diverse *V. vinifera* cultivars^45^, the characteristic expression pattern of *VIT_206s0004g06770* is highly conserved with the exception of ‘Glera’ in which expression remains low until harvest (**Supplementary Figure S33**). ‘Glera’ is a high-yield cultivar producing low-sugar and high-acidity juice that is fermented into the Prosecco sparkling wine. *VIT_206s0004g06770* encodes a conserved 77-amino acid plant protein, yet functionally uncharacterized. The expression of *VIT_206s0004g06850*, which encodes a mitogen-activated protein kinase kinase kinase (MAPKKK), and *VIT_206s0004g06840*, which encodes a protein phosphatase 2C, possibly acting as regulators of signal transduction pathways, also increases with the progression of ripening (**Supplementary Figure S31**).

## DISCUSSION

Understanding the genetic control of fruit ripening is necessary for adapting the fruit developmental program to the changing needs of agriculture and in response to climate change. Spontaneous mutants that show inhibition of ripening in fruit crops, such as *ripening inhibitor* (*rin*), *non-ripening* (*nor*) and *Colorless non-ripening* (*Cnr*) in tomato, carry knockout mutations at transcription factor genes that act as master regulators. These mutants enable studies of fruit ripening physiology, but they have limited applications in agriculture because fruit appearance and edibility that are associated with the common notion of ripeness would never be attained. Positional candidate genes in the grapevine *RIPENING SPEED* locus do not correspond to any of the switch genes that were proposed to control the phase transition from berry growth to ripening^17,52,53^. This evidence supports the hypothesis that variation in the QTL region affects the speed of ripening rather than the developmental activation of the ripening program^54^, acting through a non-disruptive mechanism that does not raise concern for the exploitation of the slow*-*ripening haplotype in crop production.

Much of the research in fruit ripening molecular biology focuses on the pericarp and overlooks seeds, with some exceptions. Deluc and collaborators^55^ showed that the initiation of ripening in individual berries on the same bunch—a condition that ensures no genetic and no environmental variation—depends on variation in seed-to-berry ratio among berries. Berries with higher seed-to-berry ratio enter the ripening stage later than berries with lower seed-to-berry ratio. Berries originating from early or late blooming flowers—a condition that generates ontogenetic variation among berries—enter the ripening stage simultaneously if they have attained similar seed-to-berry ratio at the end of the lag phase^36^. Wang and collaborators^35^ showed that berries with higher seed-to-berry ratio accumulate sugars at a slower pace than berries with lower seed-to-berry ratio. Here, we observed a genetically determined inverse relationship between seed-to-berry ratio at the end of lag phase and the subsequent rate of sugar accumulation. Both seed content and berry weight are higher in slow-ripening than in fast-ripening seedlings ahead of ripening, but seed content is disproportionally higher relative to berry weight.

We discovered a major effect haplotype in crop wild relatives that serve viticulture as pest resistant rootstocks against a soil-borne aphid and are interfertile with the crop species. This haplotype slows sugar accumulation and malate breakdown in their hybrid progeny, alleviating accelerated ripening without impairing berry growth. The resulting slow-ripening phenotype is expressed in heterozygotes according to a model of dominant gene action. The slow-ripening haplotype may contain a DNA variant that is critical only for technological berry ripeness and does not affect fitness in natural populations of wild species, thereby being fortuitously passed on to and maintained in their derived rootstocks. This haplotype is easily introgressed into the genetic background of *V. vinifera* cultivars by conventional breeding—an operation that could be assisted by the genetic markers and the nucleotide sequence annotation provided by the present article (**Supplementary Tables S4-S5** and **S7**).

Further research will be needed for modifying ripening speed in traditional cultivars using new genomic techniques and without resorting to cross-breeding. Some gene functions in *RIPENING SPEED* locus are compatible with present and previous findings^34^ showing that preveraison pericarp growth is promoted and postveraison pericarp ripening is repressed by seed content. We cannot, however, exclude that other genetic or epigenetic variation in the *RIPENING SPEED* locus could dysregulate expression homeostasis, generating a steady condition that is negligible in terms of instantaneous intensity but persists perniciously over the course of berry growth and/or ripening^56,57^. ncRNA is transcribed from QTL peak region and may serve diverse functions^58–60^. Antisense transcription initiation is detectable therein and may originate from haplotype-specific alterations in chromatin state^61,62^. All things considered, slow-ripening is compatible, genetically, with the action of a dominant negative allele or epiallele and, evolutionarily, with the origin and persistence of this variant in wild populations and rootstocks.

## MATERIALS AND METHODS

### SSR Genotyping

The VvCHA×VrGdM pseudo-testcross progeny was genotyped using microsatellite markers (**Supplementary Figure S34**). PCR, capillary electrophoresis and fragment analysis were performed following the procedures described in similar mapping experiments^63^. Parental genetic maps were constructed with Lep-MAP3 and a LOD threshold of 5.0. Ungrouped markers were assigned to existing linkage groups using the JoinSingles2All module. Map distances were expressed in cM using the Kosambi mapping function (**Supplementary Figures S35-S36**).

### Phenotyping

Experimental vines were grown and phenotyped at the germplasm repository of the University of Udine (46.03 N, 13.23 E). Refractometric readings were taken with an ATC-1 hand-held refractometer 0-32 % w/Box with automatic temperature compensation (ATAGO CO., Tokyo, Japan). Diameters were taken with a caliper. Berry softening was identified by firmness-to-the-touch and colour transition rate was scored by visual perception by two trained operators. Determination of glucose, fructose, malic acid and tartaric acid was performed using HPLC. The clarified juice from 15 squeezed berries per genotype was 1:10 diluted with water, filtered with a 0.22 μm Nylon 13 mm filter (Whatman Inc., Sanford, USA), separated and measured on an Agilent 1100 HPLC system with a refractive index detector (Agilent Technologies, USA) as described by Wong and colleagues^51^. Concentration was calculated using calibration curves of standards and expressed as mg L^-1 o^f juice. Hexose and organic acid content per berry was calculated with the following formula: mg berry ^-1 =^ (1×1.3^-1^) × (g L^-1^) × (1×100 ^-1^) × [100-berry weight (g)]. *V. vinifera* cultivars were phenotyped at the germplasm repository of Vivai Cooperativi Rauscedo (46.06 N, 12.84 E).

### QTL analysis and genetic mapping

QTL analysis was performed using Rqtl. A total of 278 VvCHA×VrGdM full-siblings were genotyped with the microsatellite markers flanking the QTL peak in order to identify recombinants (**Supplementary Figure S13**). Ten full-siblings carrying recombination between VrGdM-derived homologs in this interval were genotyped via Sanger sequencing. We amplified 10 fragments amounting to 11 Kb of low-copy DNA. Amplicons were generated using the primers reported in **Supplementary Table S7**, purified with Agencourt Ampure beads and sequenced with BigDye Terminator v.3.1 chemistry on an AB3730xl sequencer.

Sequences were analysed with Phred and Phrap^64^ and visualized with Consed^65^ for the identification of heterozygous sites.

### Genomic analyses

The VrGdM genome assembly, PacBio CLR and Illumina raw reads were retrieved from the BioProject PRJNA512170^37^. The haplotype-phased assembly of ‘Kober 5BB’ (GCA_946000715.1) and PacBio HiFi raw reads were retrieved from BioProject PRJEB55013^38^. Comparisons between assembled sequences were done using wfmash^66^ and dotter^67^. Original VrGdM raw reads were aligned with the VrGdM-derived assembled haplotype of ‘Kober 5BB’ using Minimap2^68^ and BWA-MEM^69^, respectively for long and short reads, to validate structural variation and sequence accuracy. Allelic SNPs and small indels were called from short read alignments using GATK^70^ UnifiedGenotyper with the option -- heterozygosity 0.01. The following filtering parameters were applied: reference-to-alternative read counts ratio between 0.25 and 0.75 for heterozygous sites, lower than 0.1 for homozygous reference sites and higher than 0.9 for homozygous alternative sites, and not included in repetitive DNA intervals. Variant sites were annotated using the Ensembl Variant Effect Predictor^71^. The predicted gene models for ‘Kober 5BB’ were downloaded from^72^. The GFF file was manually curated in the *RIPENING SPEED* QTL using evidence from VrGdM RNA-Seq read alignments. RNA-Seq reads from pooled biological replicates of VrGdM untreated leaf and root samples (BioProject PRJNA516950) were aligned with the VrGdM-derived assembled haplotype of ‘Kober 5BB’ using STAR^73^. FPKM and TPM values of gene expression were calculated using StringTie^74^. Counts of allele-specific RNA reads spanning exonic SNPs were obtained using GATK UnifiedGenotyper with no filtering parameters. Repetitive DNA in the VrGdM-derived assembled haplotype of ‘Kober 5BB’ was identified using Repeat Masker and two databases of transposable elements^75,76^. Centromeric were identified using Repeat Masker and the tandem repeats identified by^77^. Genomic DNA of UD-53,105 was fragmented and sequenced using a Tecan Celero EZ DNA-Seq kit on an Illumina NovaSeq6000 sequencer following manufacturers’ instructions. Variant sites in ‘Chardonnay’ (BioProject PRJNA388292) and UD-53,105 were called from short read alignments as described for VrGdM. Coordinates of the locus and haplotypes were referenced to the *Vitis vinifera* PN40024 12Xv0 (GCA_000003745.2)^75^ and T2T (10.5281/zenodo.7751391)^77^ assemblies and the VrGdM-derived assembled haploid Chr6 of ‘Kober 5BB’ (GCA_946000715.1, OX254016.1)^38^ in **Supplementary Tables S3-S4**.

### Gene expression analysis in *V. vinifera*

RNA-Seq raw reads were retrieved from Gene Expression Omnibus (GSE98923^17^, GSE62744 and GSE62745^45^), ENA (PRJEB30338^78^) and NCBI (PRJNA316157^51^ and PRJEB39261-4^15^) and aligned with the GCA_000003745.2 assembly using STAR^73^. FPKM and TPM values were obtained from public dataset associated with literature reports where available or were calculated from raw reads using StringTie^74^ and the V2.1 version^79^ of predicted gene models for GCA_000003745.2.

## ACKNOWLEDGEMENTS

We thank Pierre-François Bert for kindly providing VrGdM pseudohaplotypes described in^37^ and associated with the BioProject PRJNA512170.

## AUTHORS CONTRIBUTION

L.F., S.D.C. and G.D.G. conceived the study and collected field data; L.F. performed genetic mapping; G.M. performed genome analyses; S.D.C., G.A.G. and M.A.M. generated HPLC data; G.M. and G.D.G. plotted graphs and assembled figures; G.D.G. and M.M. wrote the manuscript; all authors contributed to data interpretation and critically revised the manuscript.

## DATA AVAILABILITY

Raw reads are deposited in NCBI under the BioProject number PRJNA1118063. Genotypic and phenotypic datasets are deposited in the figshare database DOI: 10.6084/m9.figshare.25991668. **Supplementary Data** consisting of DNA and RNA read alignments, curated gene predictions and masked repetitive DNA regions, and polymorphic sites for generating Integrative Genome Viewer tracks of the features described in this article are included in DOI: 10.6084/m9.figshare.25991668.

