## Supplementary Figures for "Grape ripening speed slowed down using natural variation"

**Supplementary Figures S1-S36**

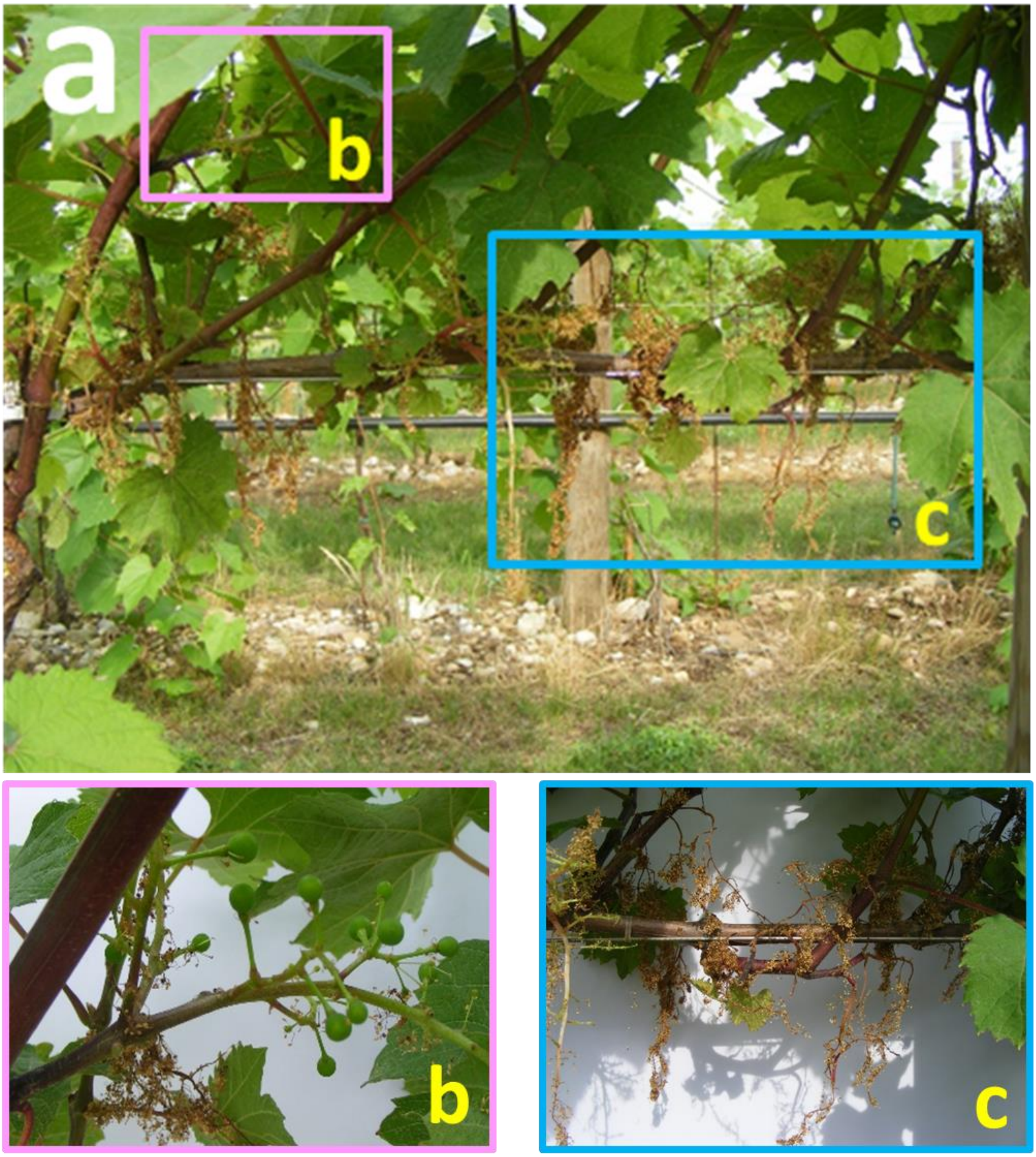


**Supplementary Figure S1 - Andromonoecy in the VvCHA×VrGdM progeny two weeks after flowering. a** A two year-old horizontal fruiting cane giving rise to vertically-positioned shoots that bear male-type inflorescences and mostly staminate flowers. **b** Magnified view of an inflorescence in **a** with sparse fruit set, turning into a loose bunch with developing berries. **c** Magnified view of a withering inflorescence in **a** after pollen shedding with no fruit set, facing abscission. Photographs of the magnified details were taken with contrast provided by a white paper background.

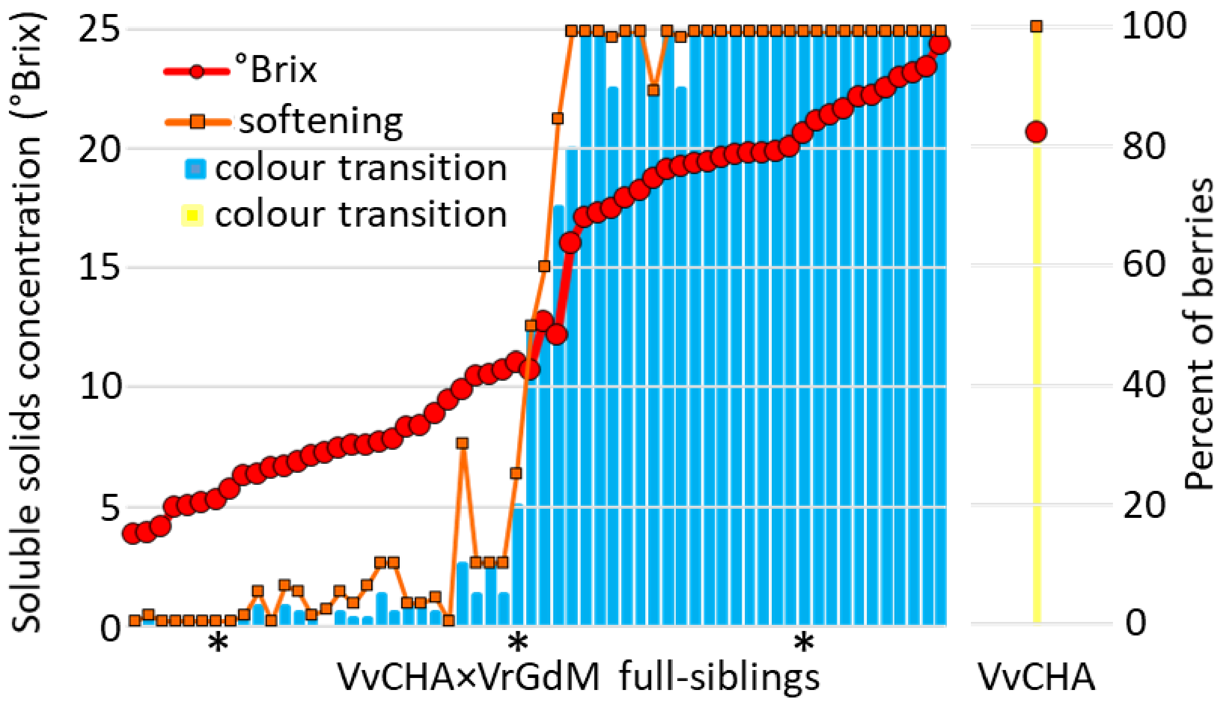


**Supplementary Figure S2 - Discovery of a ripening-related trait segregation in VvCHA×VrGdM full-siblings at harvest time in the first decade of September 2008.** Distribution of green-to-red colour transition rate (% red berries), softening rate (% soft-to-the-touch berries) and soluble solids concentration in the progeny and in the hermaphrodite seed parent VvCHA as of September 8$^{th}$, 2008. Due to impaired anthocyanin biosynthesis in VvCHA, opaque-to-translucent berry transition rate is reported for VvCHA. Soluble solids concentrations are averages of four random berries per seedlings, except for seedlings with lower than 25 % or higher than 75 % soft berries for which averages were calculated using two hard berries and two soft berries. Softening and colour transition were estimated by firmness-to-the-touch and by visual assessment. Asterisks indicate the seedlings pictured in **Figure 1j-l**.

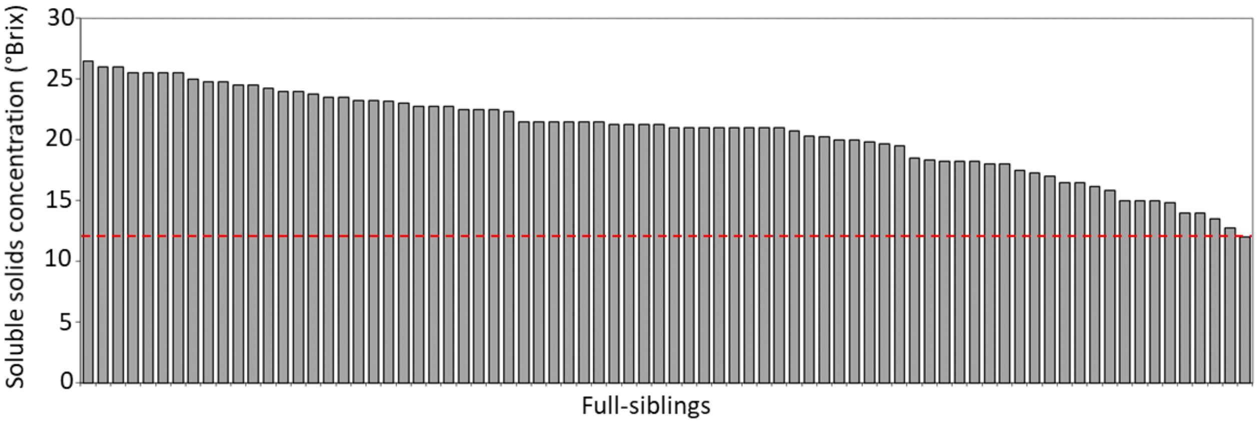


**Supplementary Figure S3 - Soluble solids concentration in the VvCHA×VrGdM progeny as of October 10th, 2008.** At that time, all seedlings surpassed a soluble solids concentration of 12°Brix (red dashed line) and completed or commenced the phase of green-to-red colour transition.

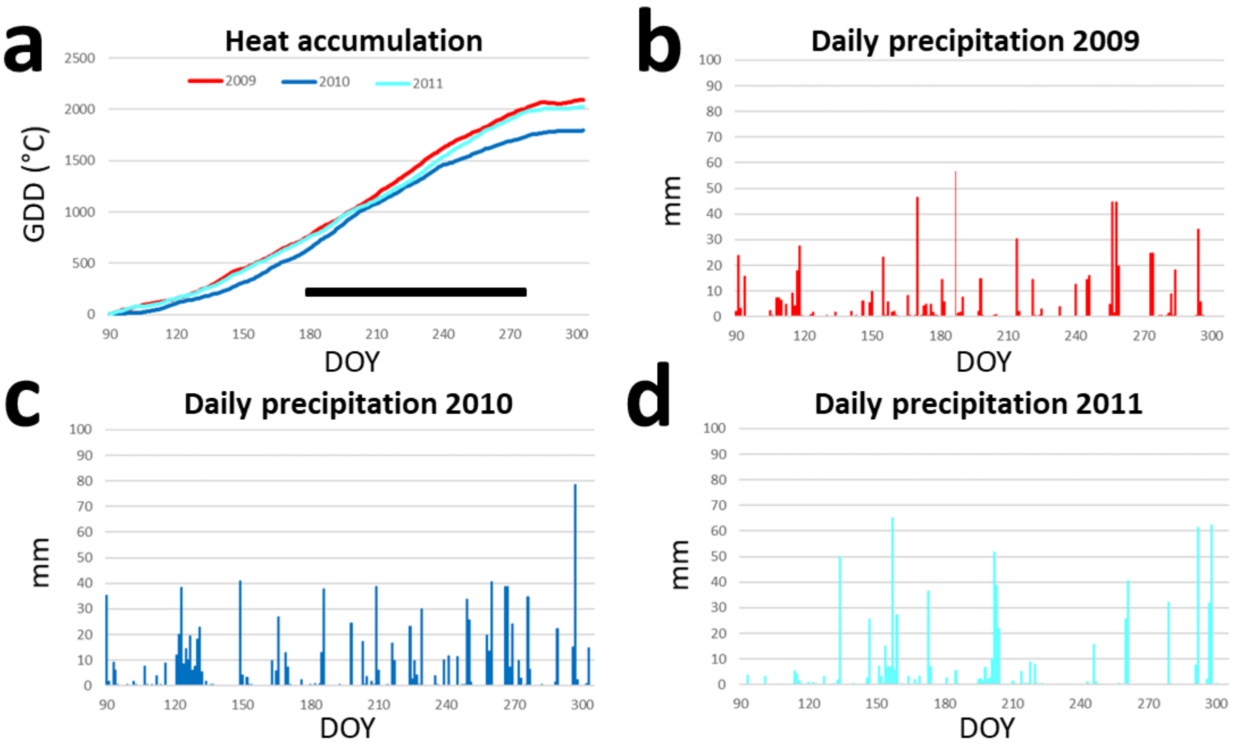


**Supplementary Figure S4 - Annual and interannual variability in heat accumulation and precipitation at the experimental site.** Growing degree days in °Celsius ($GDD^C$) and daily precipitation (mm) in the 2009, 2010 and 2011 seasons. The horizontal black line in **a** spans the interval of time shown in **Supplementary Figure S6**, corresponding to the window of time in which QTL mapping of soluble solids concentration was performed using data recorded at approximately weekly intervals.

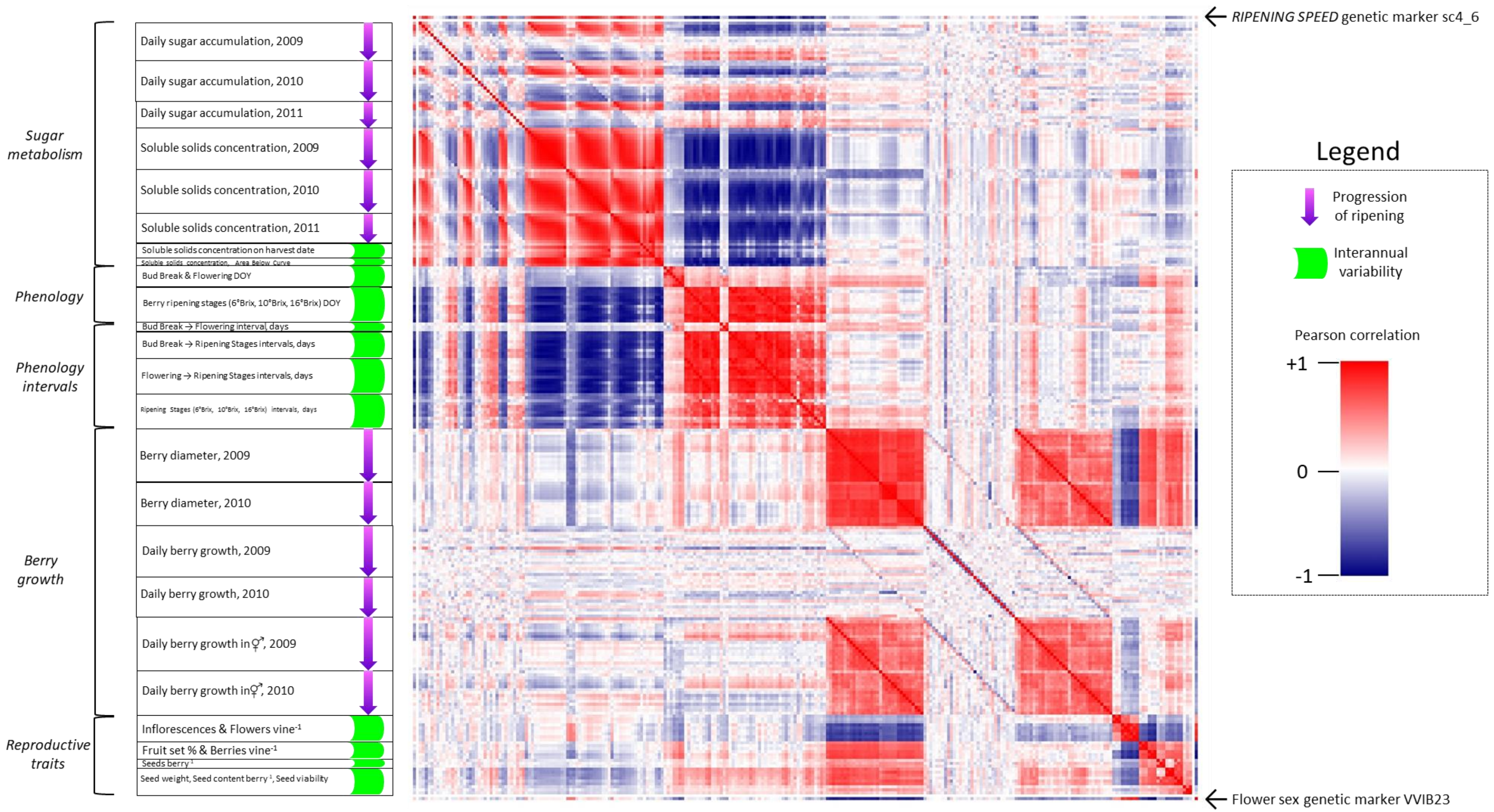


**Supplementary Figure S5 - Relations among phenotypic traits related to sugar metabolism, phenology, berry growth and reproductive (inflorescence-, bunch- and seed-related) traits.** Correlation with the most tightly linked genetic markers in the *RIPENING SPEED* QTL and in flower sex locus is shown at the edges of the matrix. The order of phenotypic traits in the present matrix follows the order reported in **Supplementary Tables S1** and **S2**, in which each trait is defined in detail. The subcategory "Daily berry growth in hermaphrodites" in the "Berry growth" category of the present matrix is a subset of "Daily berry growth" (in the entire progeny) data, it is not included in **Supplementary Tables S1** and **S2,** but phenotypic traits therein follow the same order as in the subcategory “Daily berry growth”.

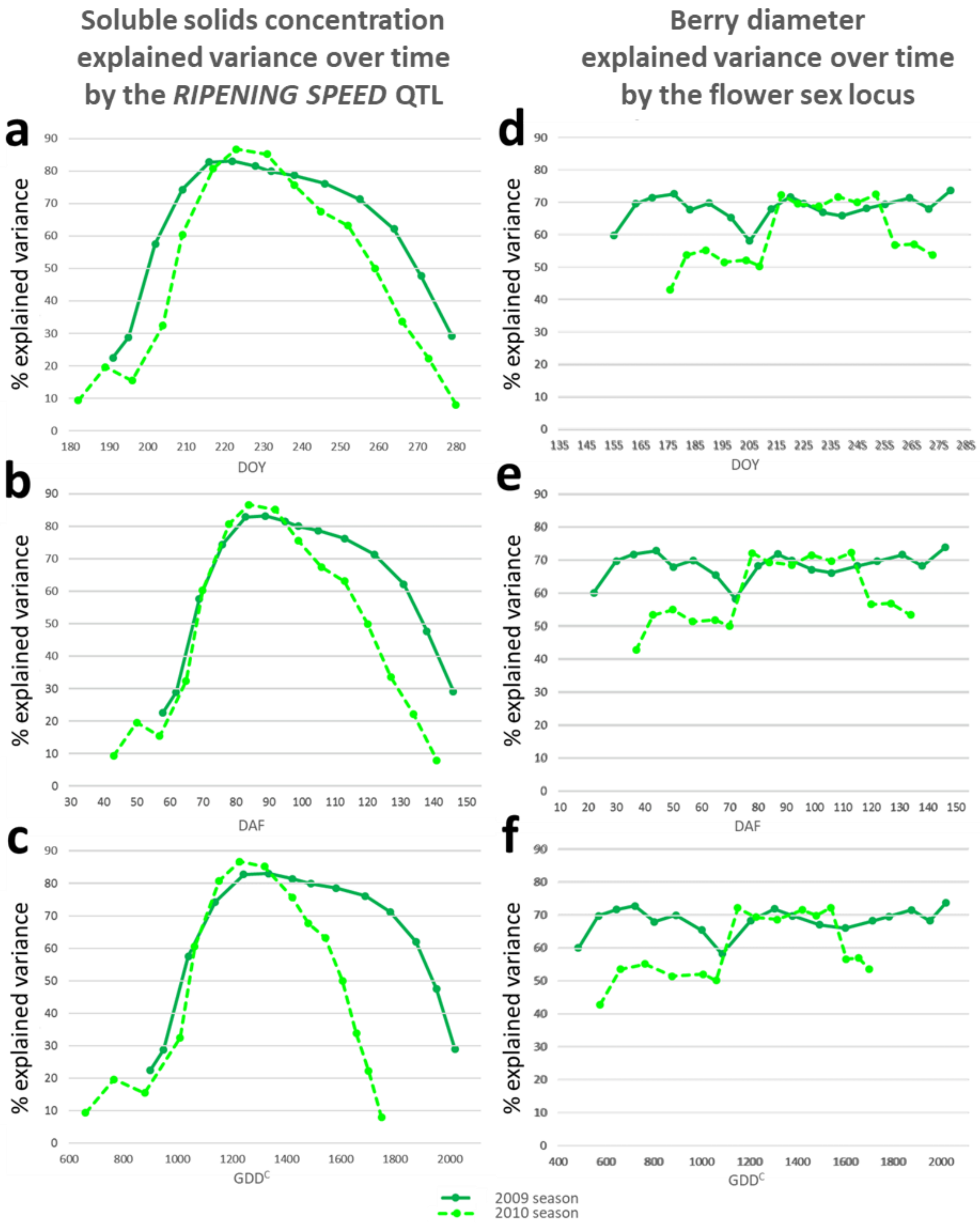


**Supplementary Figure S6 - Variation over time in the percentage of explained phenotypic variance for soluble solids concentration accounted for by the *RIPENING SPEED* QTL on Chr6 and for berry diameter accounted for by the flower sex locus on Chr2, in the 2009 (forest green) and in the 2010 season (green).** Time course on the x-axis is expressed as day of year (DOY, **a** and **d**), days after flowering (DAF, **b** and **e**, i.e. days after the average date of flowering across seedlings), and growing degree days ($GDD^C$, **c** and **f**).

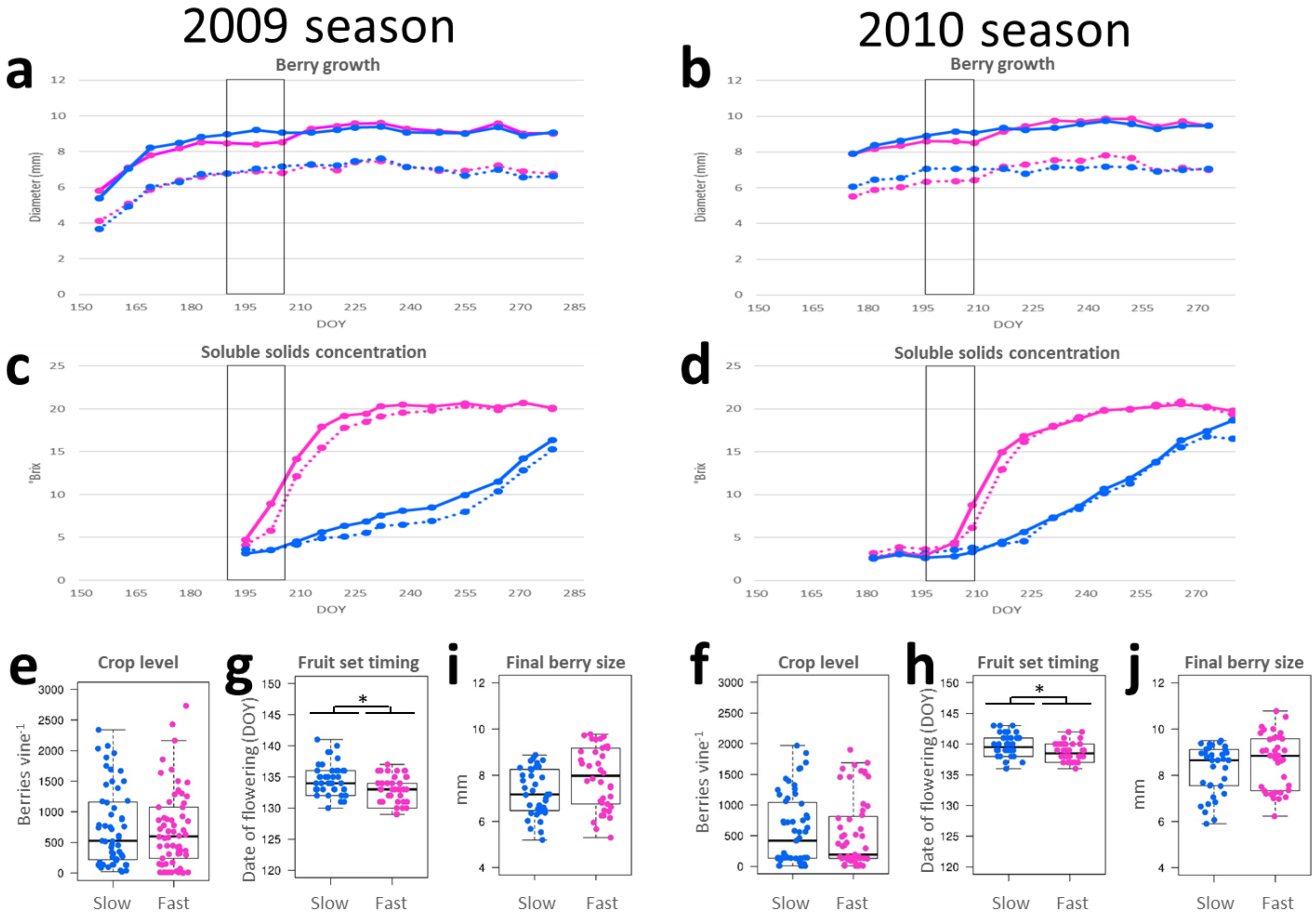


**Supplementary Figure S7 - Average berry diameter (a-b) and average soluble solids accumulation curves (c-d) across two seasons in VvCHA×VrGdM fast-ripening (magenta) and slow-ripening (blue) genotypes, further subdivided into hermaphrodite (solid lines) and andromonoecious (dashed lines) genotypes and variation in cofactors (e-j) that may affect those parameters.** Averages among individuals are referred to the DOY of monitoring. The range of variation in shifts between DOY and DAF among seedlings is shown in **Figure 2**. The period encompassing the lag-phase of berry growth and the beginning of the nearly-linear phase of soluble solids accumulation in VvCHA×VrGdM fast-ripening genotypes, corresponding in VvCHA×VrGdM slow-ripening genotypes to a prolonged phase of berry growth concentration is shown by the boxed area. Variation of crop level (**e**-**f**), inception of berry development (**g**-**h**) and final berry size (**i**-**j**) among slow-ripening and fast-ripening genotypes in the 2009 season (**e-i)** and in the 2010 season (**f-j)**. The latest stage of berry growth shown in **i-j** corresponds to DOY279 and DOY273, 146 and 134 days after the average date of flowering, in 2009 and 2010, respectively. Significant differences between slow-ripening and fast-ripening genotypes in the distributions shown in **e**-**j** were determined using a two-sided Wilcoxon test (* <0.05, ** <0.01, ***<0.001).

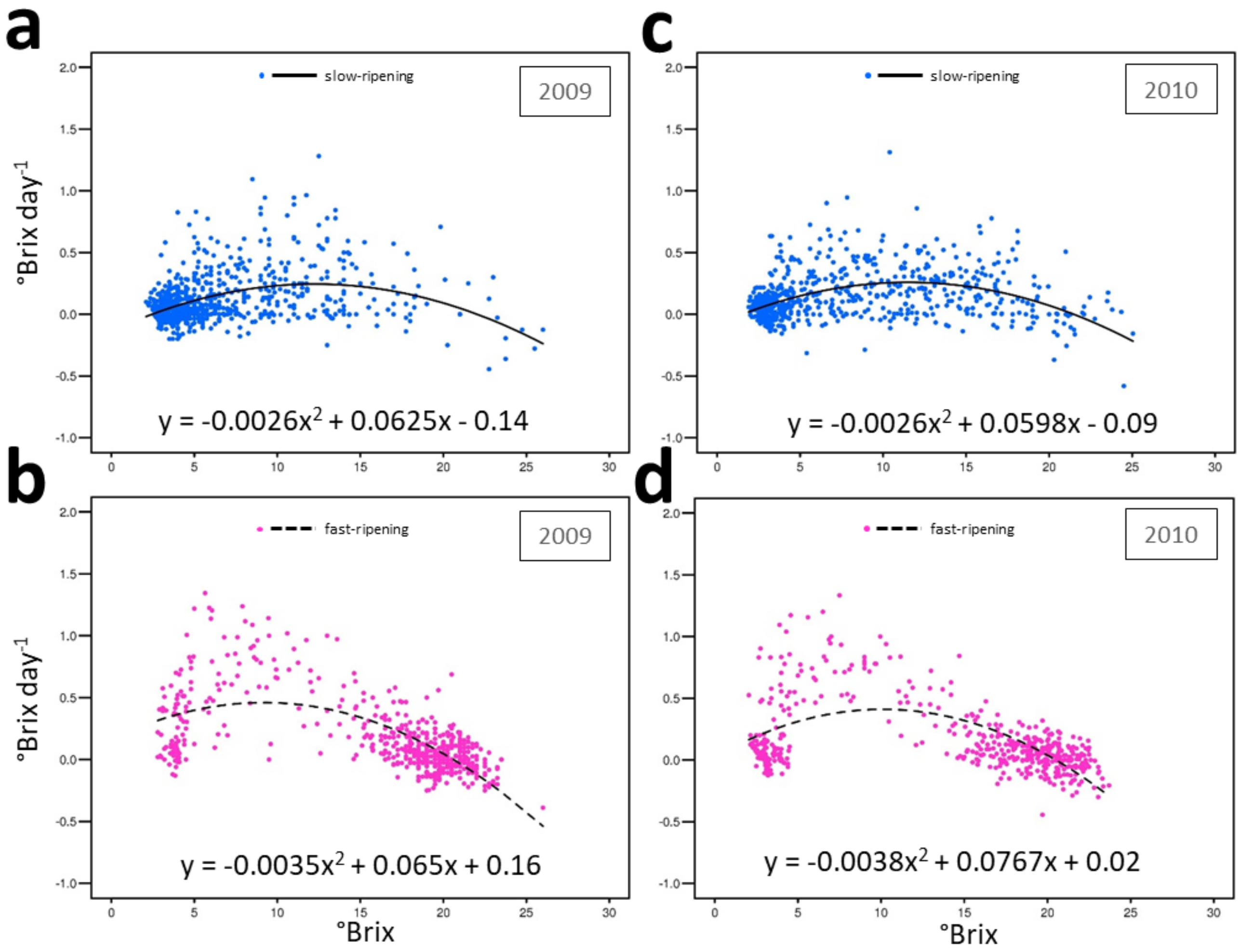


| Season | Class | Brix $day^{-1}{}_{max}$ | @ Brix |
|---|---|---|---|
| 2009 | Slow−ripening | 0.2447344 | 12.25 |
| 2010 | Slow−ripening | 0.2585707 | 11.5 |
| 2009 | Fast−ripening | 0.4597689 | 9.35 |
| 2010 | Fast−ripening | 0.4104667 | 9.95 |

**Supplementary Figure S8 - Pace of soluble solids accumulation as a function of the progress of ripening in slow-ripening (blue, a and c) and fast-ripening (magenta b and d) genotypes in the 2009 (a-b) and 2010 (c-d) seasons.** Second-order polynomial regression is plottes as a black curve with the equation shown in the corresponding panel. $Y_{max}$ values of the regression curves are reported below the panels, along with the corresponding x values.

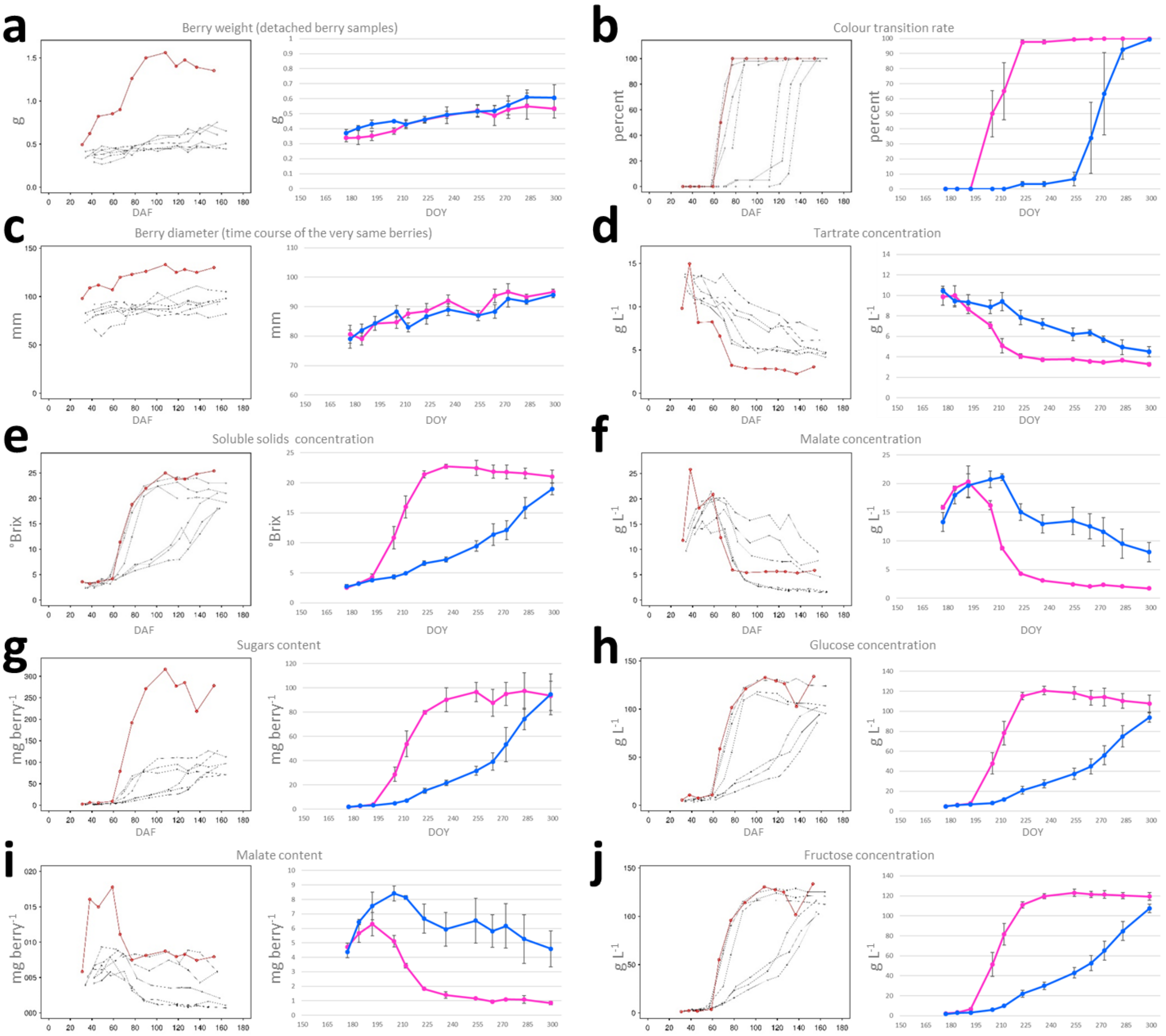


**Supplementary Figure S9 - Curves of berry growth (expressed as fresh berry weight in a, green-to-red colour transition rate in b and berry diameter in c), sugars and organic acids concentration and content per berry in six VvCHA×VrGdM full-siblings and 'Chardonnay' in the 2012 season.** In left-hand side plot of each panel, y-values are reported for every individual and are referred to individual-specific days after flowering (DAF) on the x-axis. Chardonnay is shown with red dots. In right-hand side plot of each panel, dots represent averages among three fast-ripening (magenta) and three slow-ripening (blue) full-siblings. Average values per class (y-axis) are calculated using records taken on the same day of year (DOY) and are referred to DOY of monitoring on the x-axis.

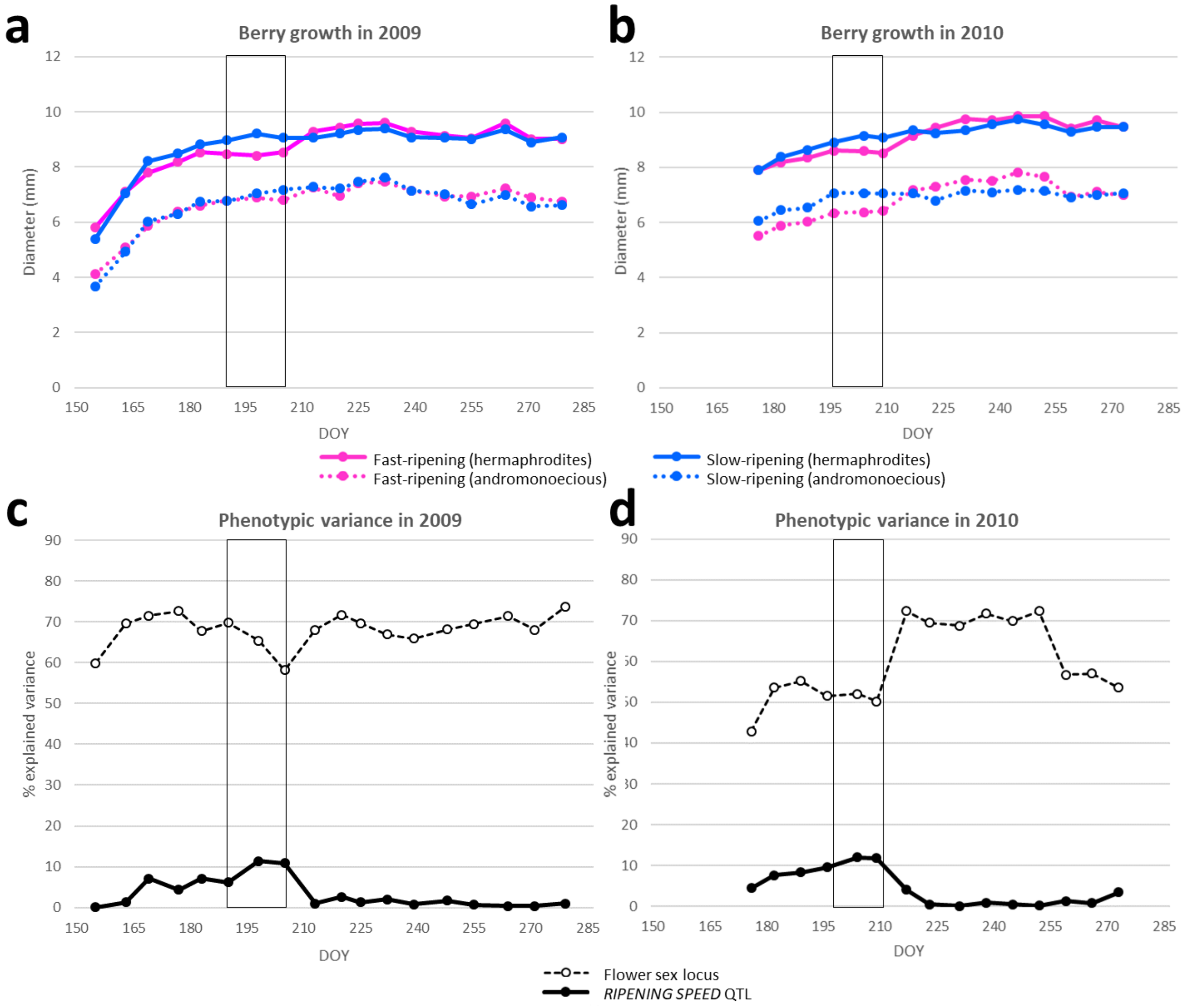


**Supplementary Figure S10 - Average berry diameters (a-b) and percent of phenotypic variance explained by the flower sex locus (c-d, open circles, dashed lines) and by the *RIPENING SPEED* QTL(c-d, solids circles and lines) across two seasons (a-c and b-d) in fast-ripening (magenta) and slow-ripening (blue) genotypes of the VxCHA×VrGdM progeny sorted in a and b into hermaphrodite (solid lines) and andromonoecious (dashed lines) genotypes.** Averages among individuals are referred to the DOY of monitoring. The range of variation in shifts between DOY and DAF among seedlings is shown in **Figure 2**. The period of time encompassing the lag-phase of berry growth and the beginning of the nearly-linear phase of soluble solids accumulation in VvCHA×VrGdM fast-ripening genotypes, and corresponding in VvCHA×VrGdM slow-ripening genotypes to a prolonged phase of berry growth followed without discontinuity by mild increments of soluble sugars concentration is shown by the boxed area.

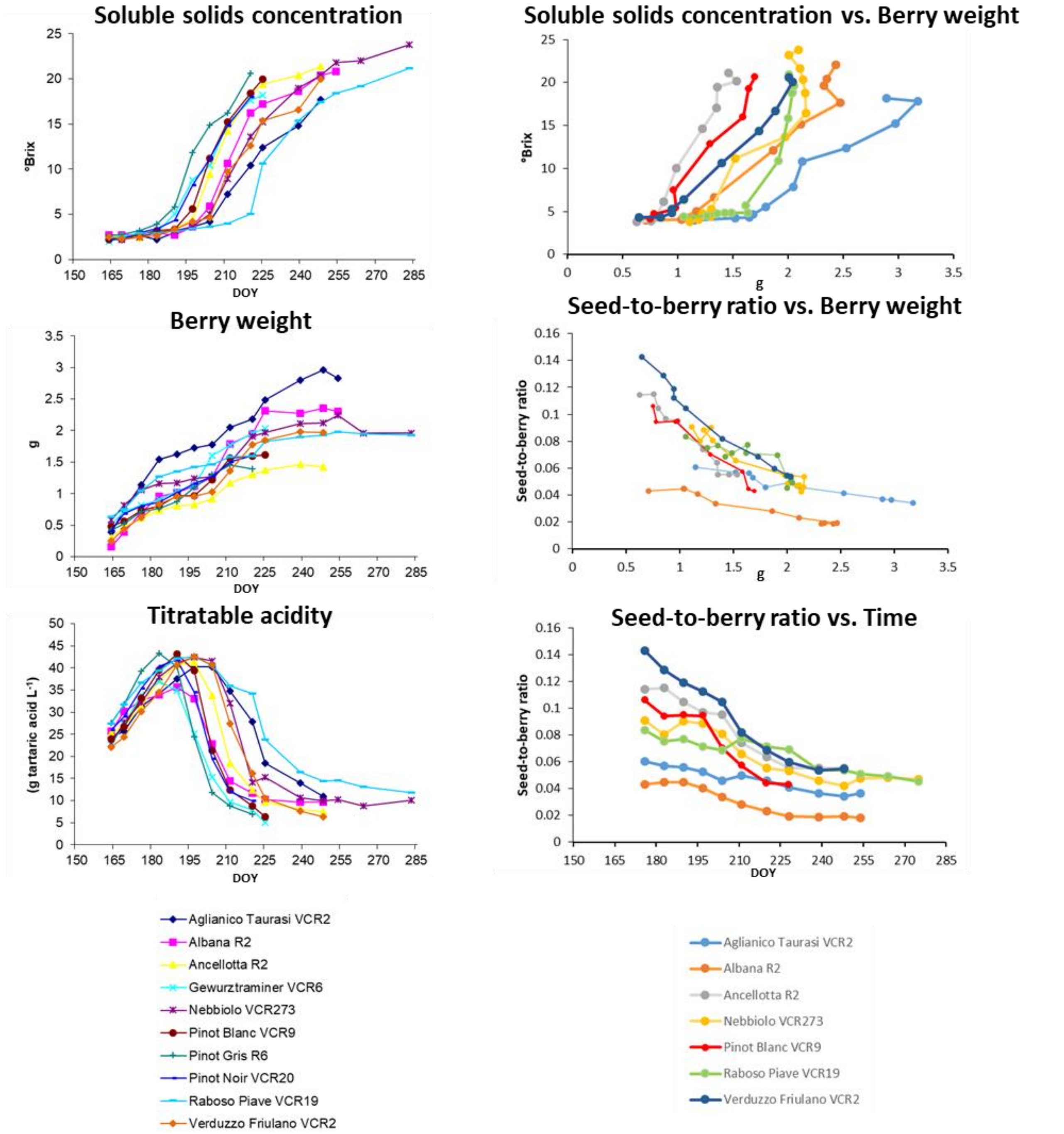


**Supplementary Figure S11 - Ripening curves in 10 *V. vinifera* genotypes and relations of sugars and seeds with berry weight.** Soluble solids concentration is used as a proxy for sugars. Seed-to-berry ratio is used to express pericarp growth relative to seed content. Samples were collected in the 2007 season at the germplasm repository of Vivai Cooperativi Rauscedo (46°03’ N, 12°50’ E). Codes following the varietal names indicate the commercial clone (bud sport) used for this analysis. ‘Pinot Noir’, ‘Pinot Blanc’ and ‘Pinot Gris’ are bud sports of a single seedling originated from sexual reproduction.

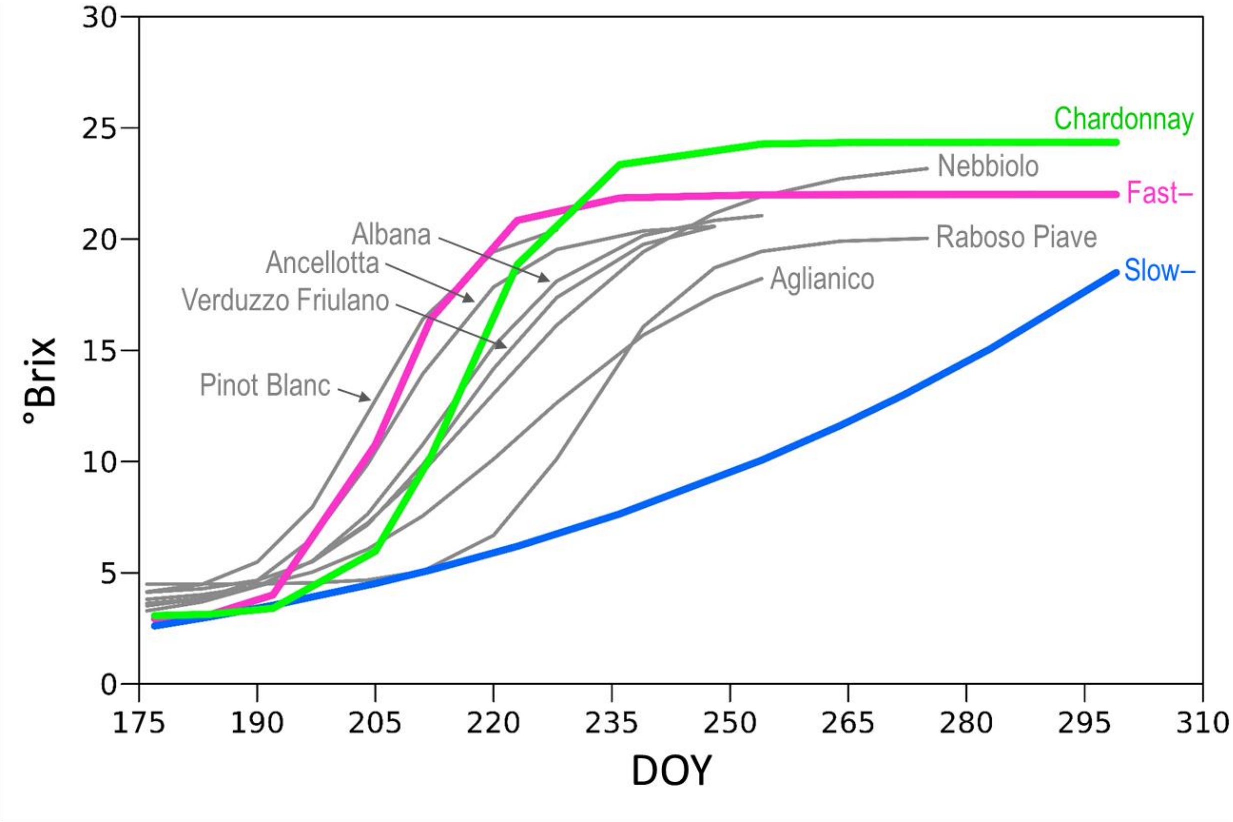


**Supplementary Figure S12 - Modelling of soluble solids accumulation (expressed as juice concentration) using a chronological scale in *V. vinifera* varieties as well as in VvCHA×VrGdM slow-ripening and fast-ripening genotypes shown in Supplementary Figure S11.** Data from the control 'Chardonnay' and VvCHA×VrGdM genotypes were taken under the same growing and monitoring conditions. Data from all *V. vinifera* cultivars and fast-ripening VvCHA×VrGdM genotypes fit a log-logistic model. Data from slow-ripening VvCHA×VrGdM genotypes fit a power model.

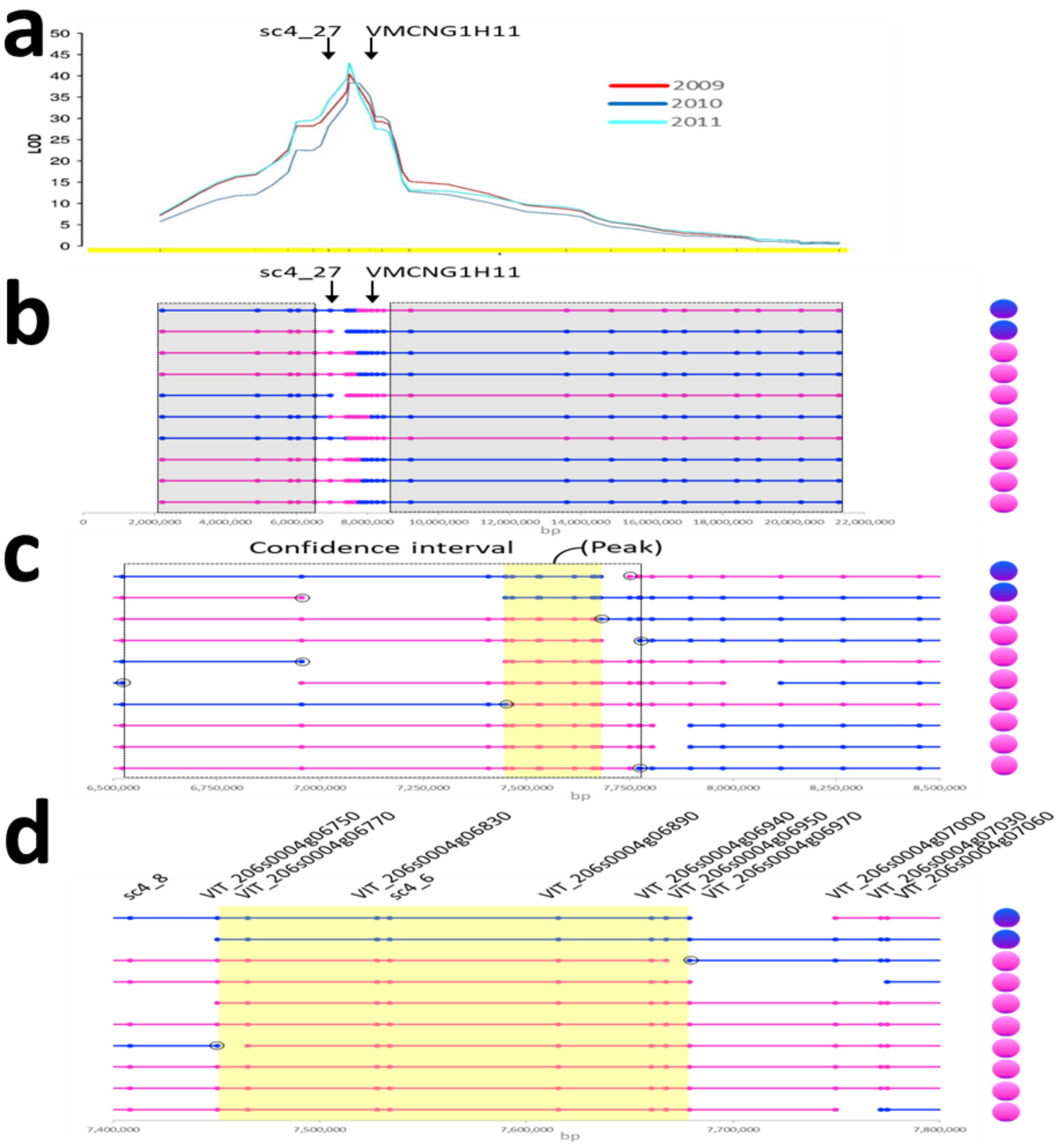


**Supplementary Figure S13 - *RIPENING SPEED* QTL peak and confidence interval. a** Chromosomal plot of the LOD score of soluble solids concentration on the day of the year on which the maximal annual LOD score was recorded annually. The horizontal yellow bar represents the chromosome physical length. Ticks on the horizontal yellow bar indicate the physical position of microsatellite markers. The relation between physical position and genetic distance of microsatellite markers is shown in **Supplementary Figure S35**. **b-d** Diagram of the recombination events between VrGdM-derived chromosomes across the *RIPENING SPEED* region in the most informative VvCHA×VrGdM seedlings. **b** Chromosome-scale representation of recombinant chromosomes. The non-shadowed interval is magnified in panel **c**. **c** Magnified view of the chromosomal region spanning the QTL peak, defined by the closest recombination event on each side and highlighted by the yellow background, and the QTL confidence interval, defined by three additional recombination events on each side and delimited by the dash-lined rectangle. The black circles indicate the informative recombination point to define the QTL confidence interval and peak. **d** Recombination points identified by segregating SNPs within predicted genes across the QTL peak region. Gene IDs and chromosomal coordinates in **b**-**d** refer to the V2.1 gene prediction (Vitulo et al 2014) of the *V. vinifera* ‘PN40024’ reference genome 12Xv0 (Jaillon et al 2007). Stylized berries indicate fast-ripening (magenta) and slow-ripening (blue) phenotypes. Stylized chromosomes indicate fast-ripening (magenta) and slow-ripening (blue) homologs in VrGdM.

Fasoli et al (2018) Timing and order of the molecular events marking the onset of berry ripening in grapevine. Plant Physiology 178:1187-1206

Jaillon et al (2007) The grapevine genome sequence suggests ancestral hexaploidization in major angiosperm phyla. Nature 449:463-467

Vitulo et al (2014) A deep survey of alternative splicing in grape reveals changes in the splicing machinery related to tissue, stress condition and genotype. BMC Plant Biology 14:99

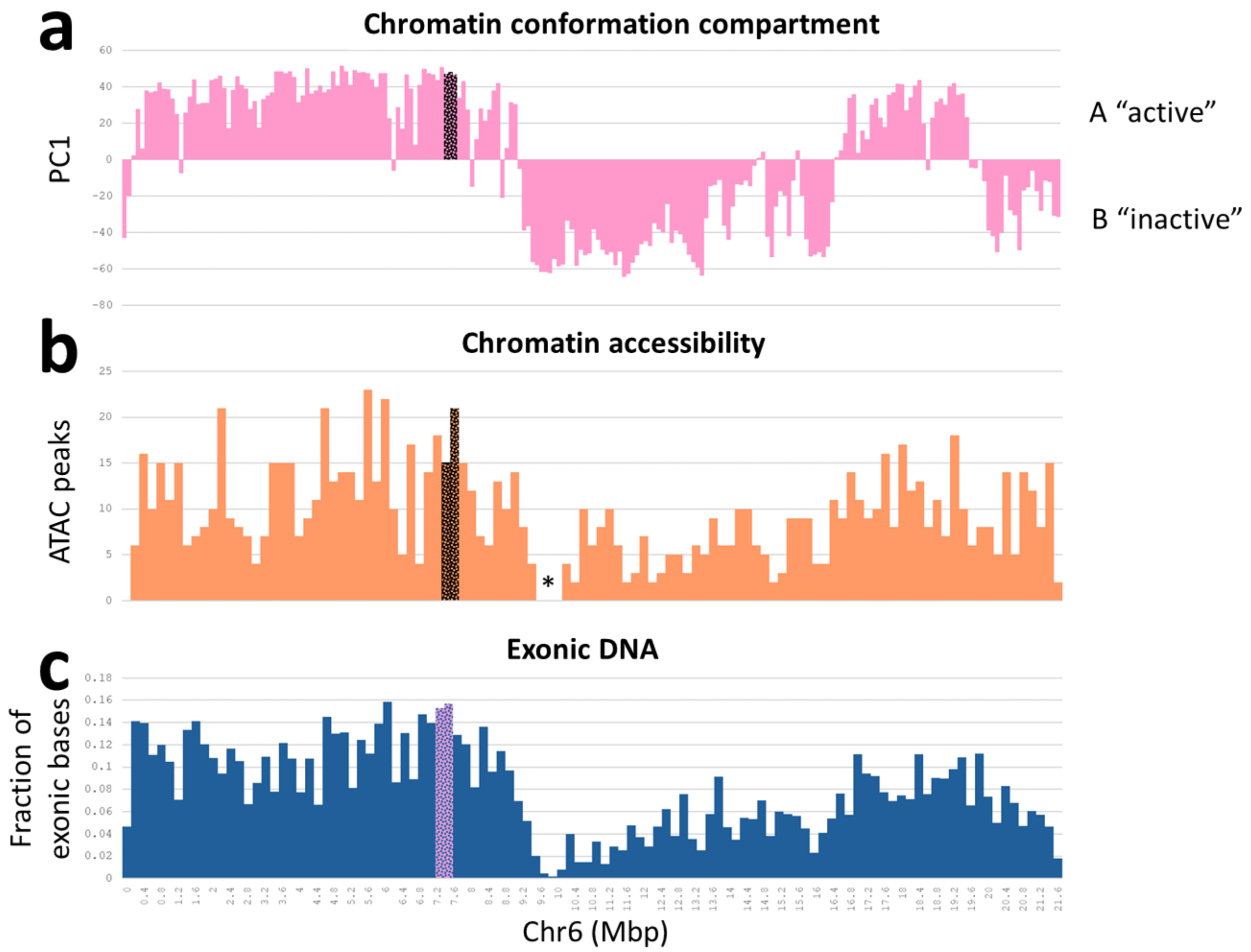


**Supplementary Figure S14 - Chromatin organization and accessibility, and gene density on Chr6, wherein the *RIPENING SPEED* QTL peak is located. a** Principal component analysis defining A/B compartments in non-overlapping genomic windows of 100 Kb using chromatin capture Hi-C data. **b** Counts of Assay for Transposase Accessible Chromatin (ATAC) peaks in non-overlapping genomic windows of 200 Kb using sequencing (ATAC-Seq) data. Data for generating the histograms in **a** and **b** were obtained from Schwope et al (2021) and analysed following the procedure described therein. **c** Gene density expressed as fraction of exonic bases in non-overlapping genomic windows of 200 Kb. Values were calculated using the GFF file of the V2.1 gene prediction from Vitulo et al (2014). The bars spanning the *RIPENING SPEED* QTL peak are highlighted by a contrasting background. Chromosome coordinates refer to the *V. vinifera* ‘PN40024’ reference genome 12Xv0 (Jaillon et al 2007). The asterisk indicate the expected location of the centromeric region.

Jaillon et al (2007) The grapevine genome sequence suggests ancestral hexaploidization in major angiosperm phyla. Nature 449:463-467

Schwope et al (2021) Open chromatin in grapevine marks candidate CREs and with other chromatin features correlates with gene expression. Plant Journal 107:1631-1647

Vitulo et al (2014) A deep survey of alternative splicing in grape reveals changes in the splicing machinery related to tissue, stress condition and genotype. BMC Plant Biology 14:99

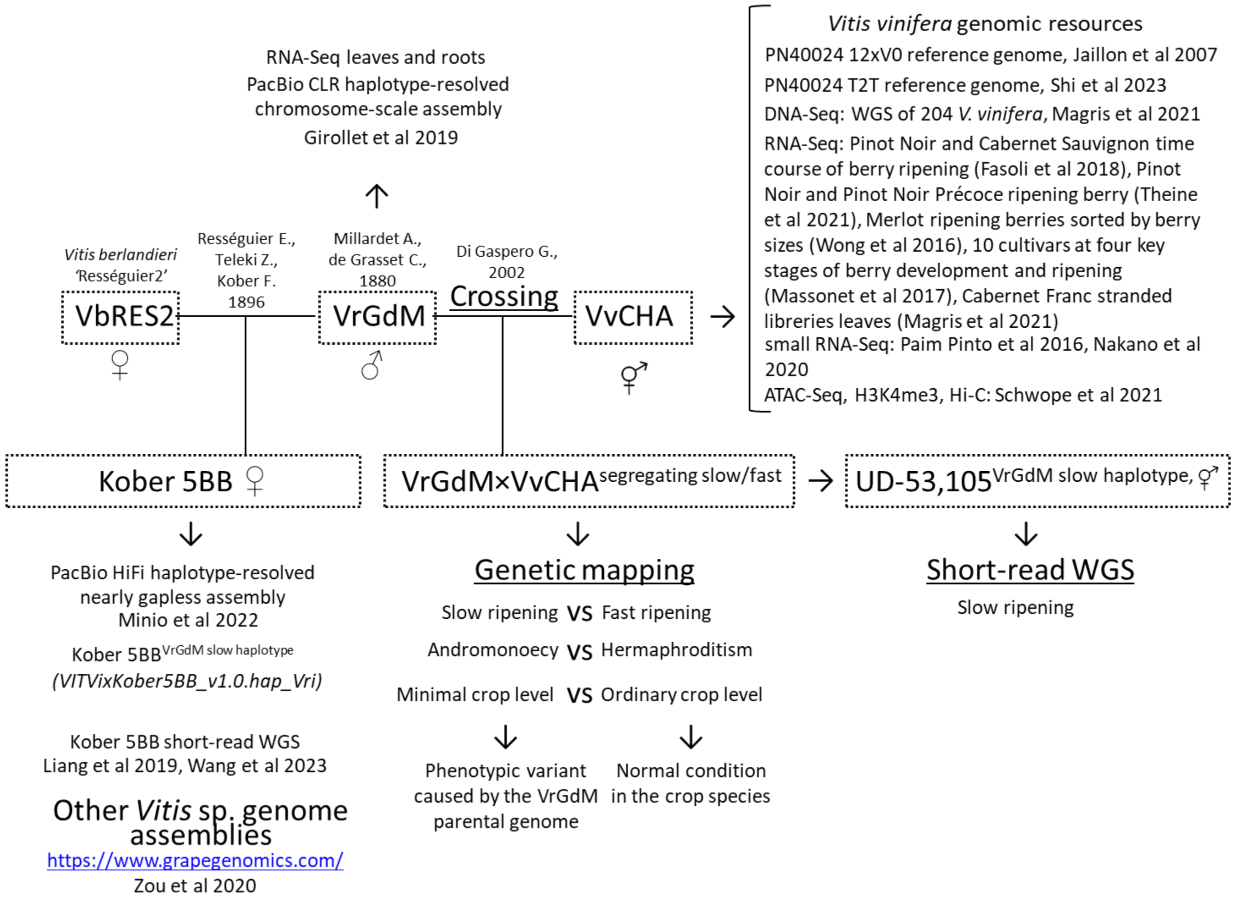


**Supplementary Figure S15 - Genetic and genomic resources for mapping and characterizing the *RIPENING SPEED* QTL.** Genetic and genomic resources generated in this study are underlined.

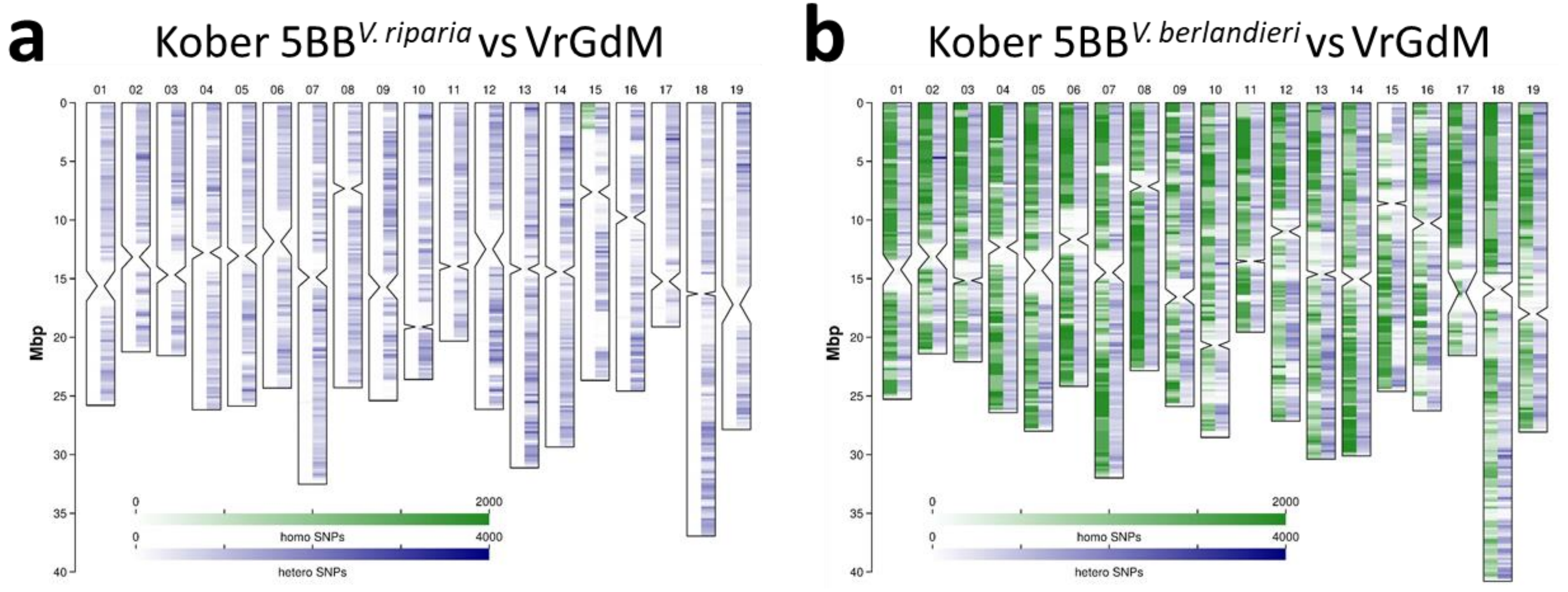


**Supplementary Figure S16 - Chromosomal patterns of SNP density and zigosity between *V. riparia* 'Gloire de Montpellier' and 'Kober 5BB'.** SNP calls are obtained from the alignment of VrGdM short reads with the 'Kober 5BB' genome assembly. Vertical ideograms represent chromosomes. Constrictions indicate the location of centromeric repeats. Chromosomal plots of heterozygous (white-to-blue heatmap shown in the right-hand portion of each chromosome) and homozygous (white-to-green heatmap shown in the left-hand portion of each chromosome) SNP densities in non-overlapping 200-Kb windows in *V. riparia* 'Gloire de Montpellier' compared to haplotype-resolved genome sequence of 'Kober 5BB'. **a** *V. riparia* 'Gloire de Montpellier' SNP densities are referred to the *V. riparia* 'Gloire de Montpellier'-derived haplotype of 'Kober 5BB'. **b** *V. riparia* 'Gloire de Montpellier' SNP densities are referred to the *V. berlandieri* 'Rességuier 2'-derived haplotype of 'Kober 5BB'. The presence of homozygous SNPs in the upper subtelomeric portion of Chr15 in **a** and the symmetric lack of homozygous SNPs over the same region in **b** is due to a switch between *V. riparia*-derived and *V. berlandieri*-derived haplotypes in the genome assembly of 'Kober 5BB'. Large blocks of consecutive white windows in both left-hand and right-hand portions of each chromosome and in both panels represent pericentromeric windows with predominance of multimapping reads that were not used for SNP calling. Values higher than 2000 in the white-to-green heatmap and higher than 4000 in the white-to-blue heatmap were plotted as maximum value of the scale.

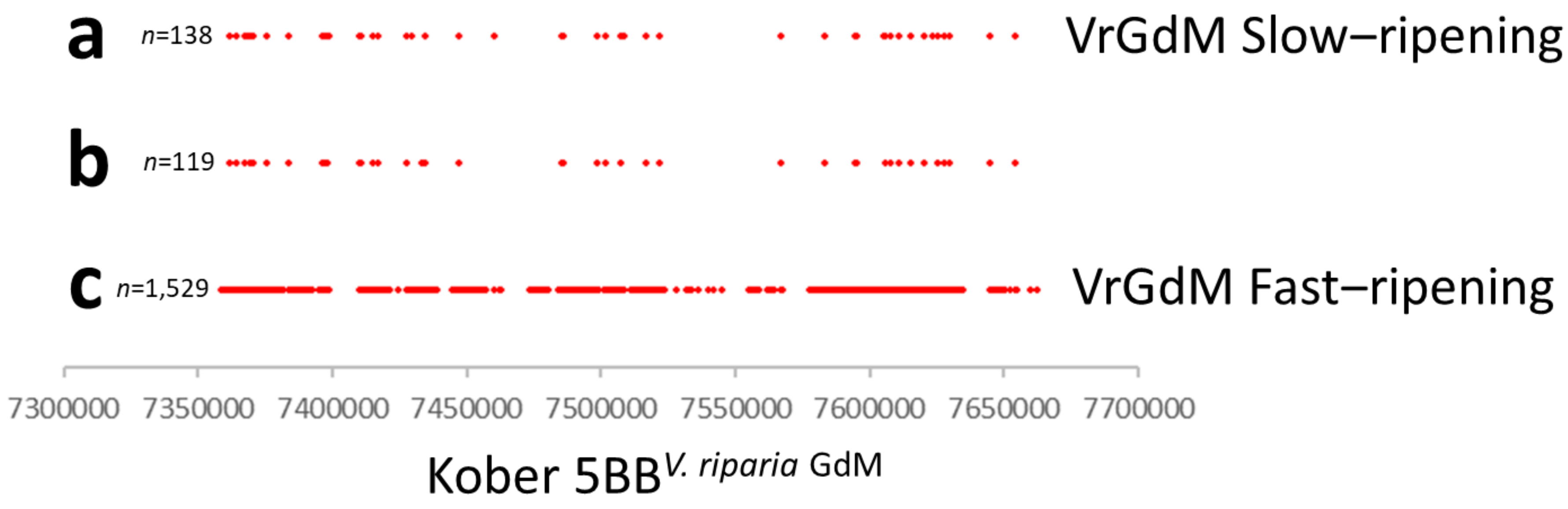


**Supplementary Figure S17 - Variant sites between either the slow-ripening haplotype or the fast-ripening haplotype in *V. riparia* 'Gloire de Montpellier' and the *V. riparia* 'Gloire de Montpellier'-derived haplotype in 'Kober 5BB'.** Red dots represent mismatches, insertions and deletions identified in pairwise sequence comparisons of the assembled sequences that were performed with the wfmash software (Marco-Sola et al 2020). **a** Variant sites between the slow-ripening haplotig in VrGdM (VrGdM 349.RGM.pseudoHap2.1) and the *V.riparia*-derived chromosome pseudomolecule in 'Kober 5BB'. **b** Variant sites between both the slow-ripening and the fast-ripening haplotigs in VrGdM compared to the *V.riparia*-derived chromosome pseudomolecule in 'Kober 5BB', likely representing sequence inaccuracies in either assembly. **c** Variant sites between the fast-ripening haplotig in VrGdM (VrGdM 349.RGM.pseudoHap2.2 ) and the *V.riparia*-derived chromosome pseudomolecule in 'Kober 5BB'. *n* indicates the number of variant sites.

Marco-Sola et al (2020) Fast gap-affine pairwise alignment using the wavefront algorithm. Bioinformatics 37:456-463

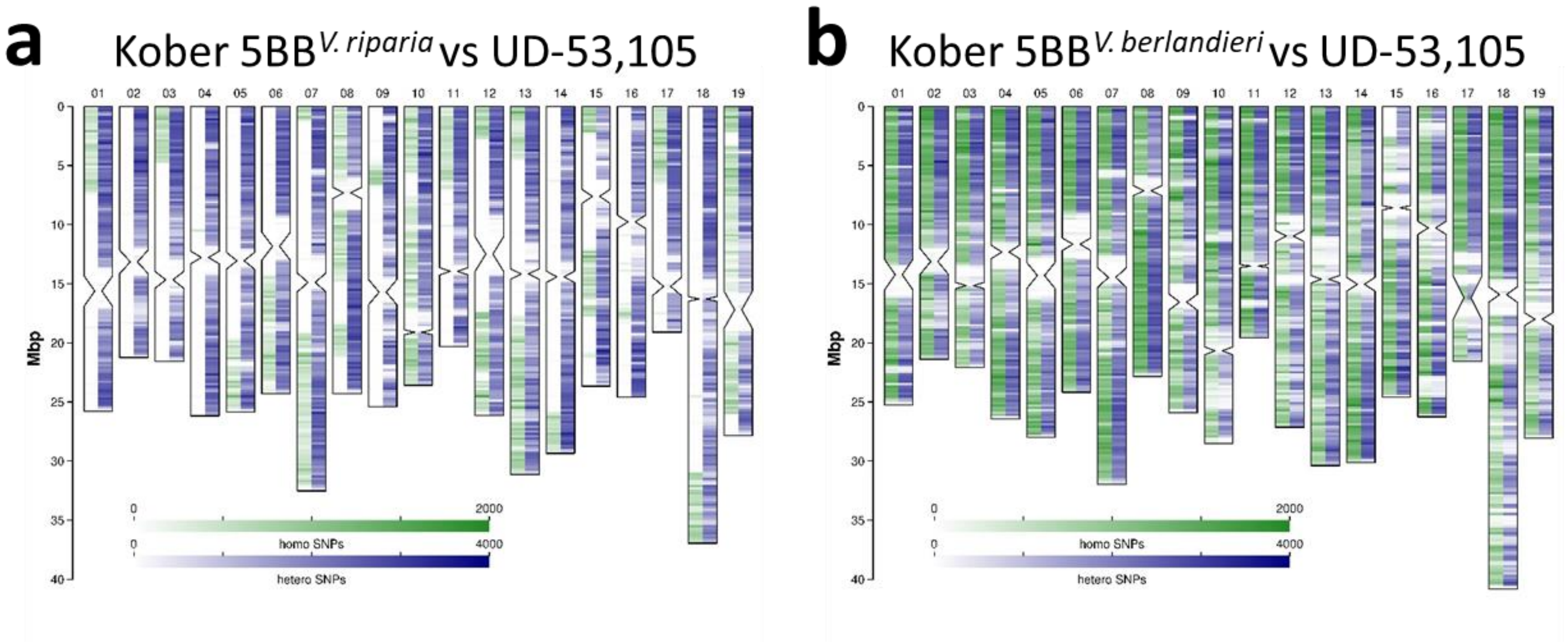


**Supplementary Figure S18 - Chromosomal patterns of SNP density and zigosity between 'Kober 5BB' and UD-53,105.** SNP calls are obtained from the alignment of UD-53,105 short reads with the 'Kober 5BB' genome assembly. Vertical ideograms represent chromosomes. Constrictions indicate the location of centromeric repeats. Chromosomal plots of heterozygous (white-to-blue heatmap shown in the right-hand portion of each chromosome) and homozygous (white-to-green heatmap shown in the left-hand portion of each chromosome) SNP densities in non-overlapping 200-Kb windows in the VvCHA×VrGdM offspring UD-53,105 compared to haplotype-resolved genome sequence of 'Kober 5BB'. UD-53,105 is a slow-ripening offspring. **a** SNP densities in UD-53,105 are referred to the *V. riparia* 'Gloire de Montpellier'-derived haplotype of 'Kober 5BB'. **b** SNP densities in UD-53,105 are referred to the *V. berlandieri* 'Resseguier 2'-derived haplotype of 'Kober 5BB'. The presence of homozygous SNPs in the upper subtelomeric portion of chr15 in **a** and the symmetric lack of homozygous SNPs over the same region in **b** is due to a switch between *V. riparia*-derived and *V. berlandieri*-derived haplotypes in the genome assembly of 'Kober 5BB'. Large blocks of consecutive white windows in both left-hand and right-hand portions of each chromosome and in both panels represent pericentromeric windows with predominance of multimapping reads that were not used for SNP calling. Values higher than 2000 in the white-to-green heatmap and higher than 4000 in the white-to-blue heatmap were plotted as maximum value of the scale.

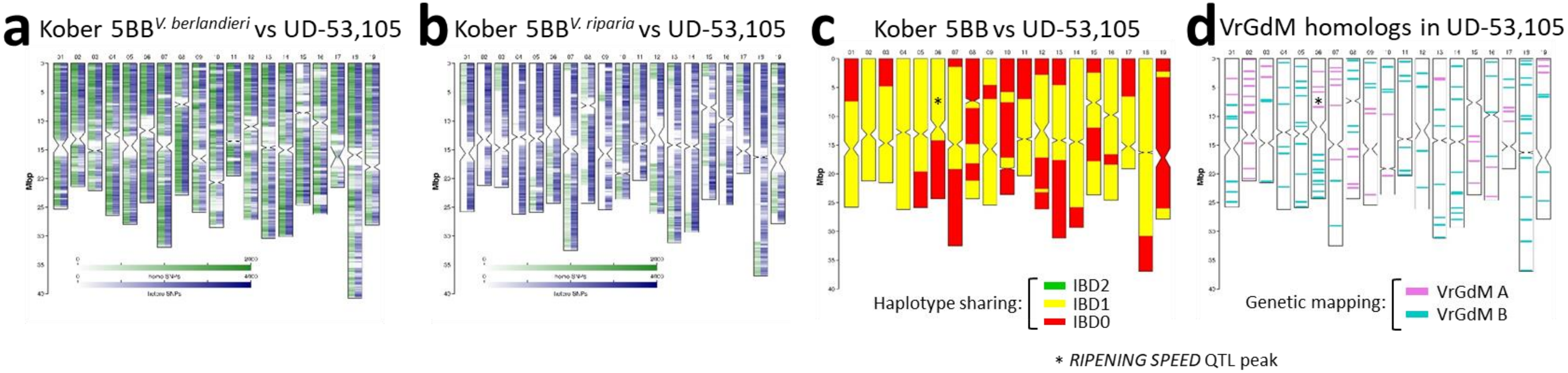


**Supplementary Figure S19 - Chromosomal patterns of SNP density and zigosity in UD-53,105 and haplotype sharing between 'Kober 5BB' and UD-53,105.** SNP density and zigosity in **a** and **b** is obtained from SNP calls of the alignment of short reads with the 'Kober 5BB' genome assembly. Chromosomal plots of heterozygous (white-to-blue heatmap shown in the right-hand portion of each chromosome) and homozygous (white-to-green heatmap shown in the left-hand portion of each chromosome) SNP densities in non-overlapping 200-Kb windows in UD-53,105 compared to haplotype-resolved genome sequence of 'Kober 5BB'. **a** SNP densities in UD-53,105 are referred to the *V. riparia* 'Gloire de Montpellier'-derived haplotype of 'Kober 5BB'. **b** SNP densities in UD-53,105 are referred to the *V. berlandieri* 'Rességuier 2'-derived haplotype of 'Kober 5BB'. Large blocks of consecutive white windows in both left-hand and right-hand portions of each chromosome and in both panels represent pericentromeric windows with predominance of multimapping reads that were not used for SNP calling. Values higher than 2000 in the white-to-green heatmap and higher than 4000 in the white-to-blue heatmap were plotted as maximum value of the scale. **c** Haplotype sharing between **'**Kober 5BB' and UD-53,105 inferred from the plots in **a** and **b**. Colours indicate the segments within each chromosome wherein the pair shares two haplotypes (IBD=2, green), one haplotype (IBD=1, yellow), or no haplotype (IBD=0, red). **d** Recombination of parental VrGdM chromosome in the germ cell that produced the male gamete at the origin of UD-53,105 based on genetic mapping data of segregating microsatellite markers. Vertical ideograms in all panels represent chromosomes. Constrictions indicate the location of centromeric repeats. The asterisks in **c** and **d** indicate the location of the *RIPENING SPEED* QTL peak on Chr6.

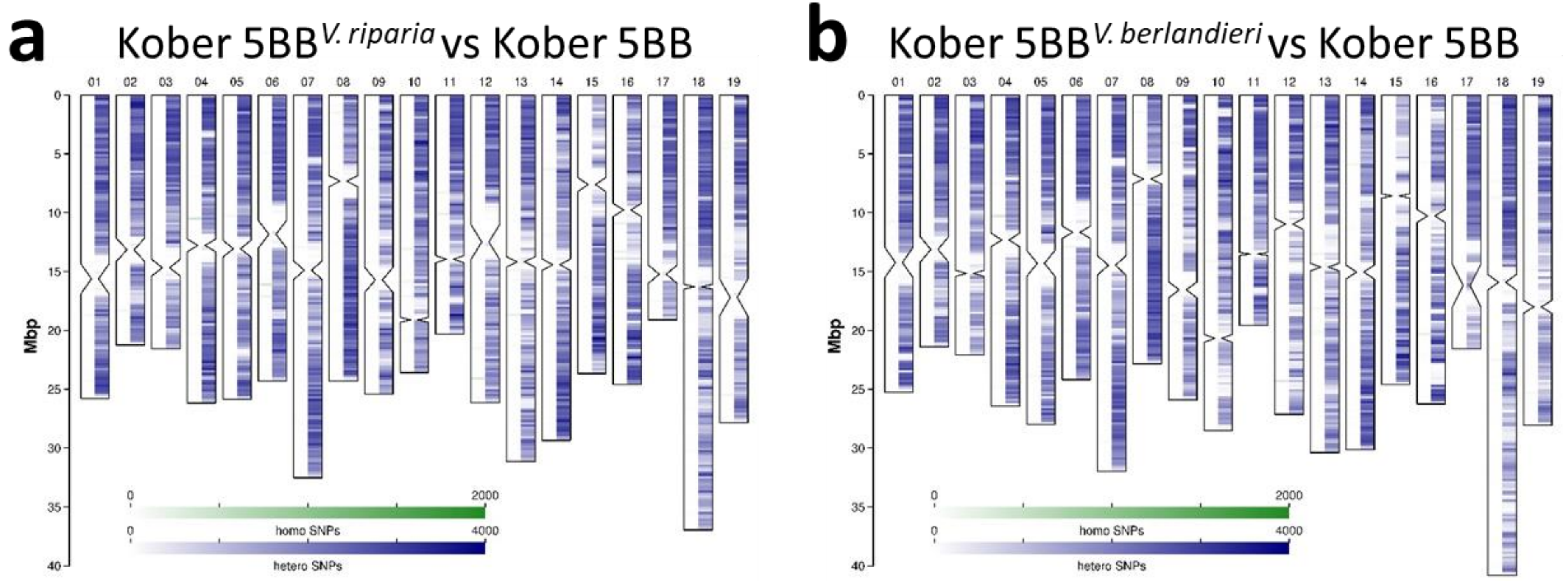


**Supplementary Figure S20 - Chromosomal patterns of SNP density and zigosity in different specimens of 'Kober 5BB'.** SNP calls are obtained from the alignment of three 'Kober 5BB' short read batches [samples TA-215 (SRR5891618) and TA-6141 (SRR5891617) from Liang et al (2019) BioProject PRJNA393611 and 5BB Austria (SRR15962938) from Wang et al (2023) BioProject PRJNA764455) with the 'Kober 5BB' genome assembly. The three batches of Illumina reads were merged to obtain sufficient coverage, following the assessment that they were all generated from the same genotype. Vertical ideograms represent chromosomes. Constrictions indicate the location of centromeric repeats. Chromosomal plots of heterozygous (white-to-blue heatmap shown in the right-hand portion of each chromosome) and homozygous (white-to-green heatmap shown in the left-hand portion of each chromosome) SNP densities in non-overlapping 200-Kb windows in 'Kober5BB' compared to haplotype-resolved genome sequence of 'Kober 5BB'. **a** SNP densities in Kober 5BB are referred to the *V. riparia* 'Gloire de Montpellier'-derived haplotype of 'Kober 5BB'. **b** SNP densities in 'Kober 5BB' are referred to the *V. berlandieri* 'Rességuier 2'-derived haplotype of 'Kober 5BB'. Large blocks of consecutive white windows in both left-hand and right-hand portions of each chromosome and in both panels represent pericentromeric windows with predominance of multimapping reads that were not used for SNP calling. Values higher than 2000 in the white-to-green heatmap and higher than 4000 in the white-to-blue heatmap were plotted as maximum value of the scale.

Liang et al (2019) Whole-genome resequencing of 472 *Vitis* accessions for grapevine diversity and demographic history analyses. Nature Communications 10:1190

Wang et al (2023) Whole-genome re-sequencing, diversity analysis, and stress-resistance analysis of 77 grape rootstock genotypes. Frontiers in Plant Science 14:1102695

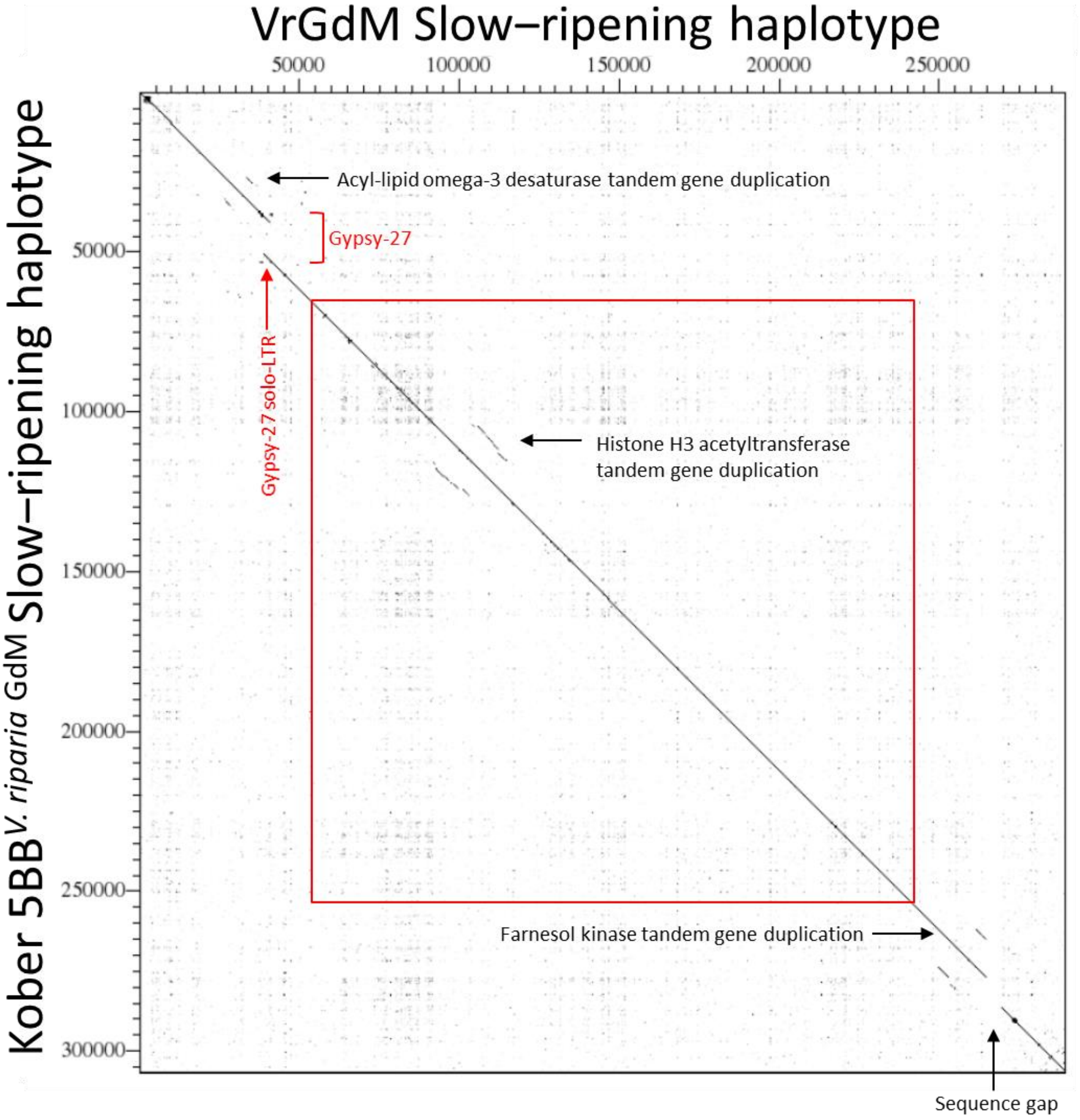


**Supplementary Figure S21 - Dotplot comparison of the slow-ripening haplotype assembled from VrGdM (horizontal, from PacBio reads) and the slow-ripening haplotype assembled from 'Kober 5BB' (vertical, from HiFi PacBio reads).** The red rectangle delimits the *RIPENING SPEED* QTL peak. The absence of the Gypsy-27 element in the assembly obtained from PacBio reads (horizontal) is not supported by their realignment against the assembly obtained from HiFi PacBio reads (vertical). The sequence gap indicated by the black arrow in the assembly obtained from PacBio reads (horizontal) has been filled in the assembly obtained from HiFi PacBio reads (vertical). See Integrative Genome Viewer (Falginella et al 2024).


Falginella et al (2024) Grape ripening speed slowed down using natural variation. *figshare* 10.6084/m9.figshare.25991668

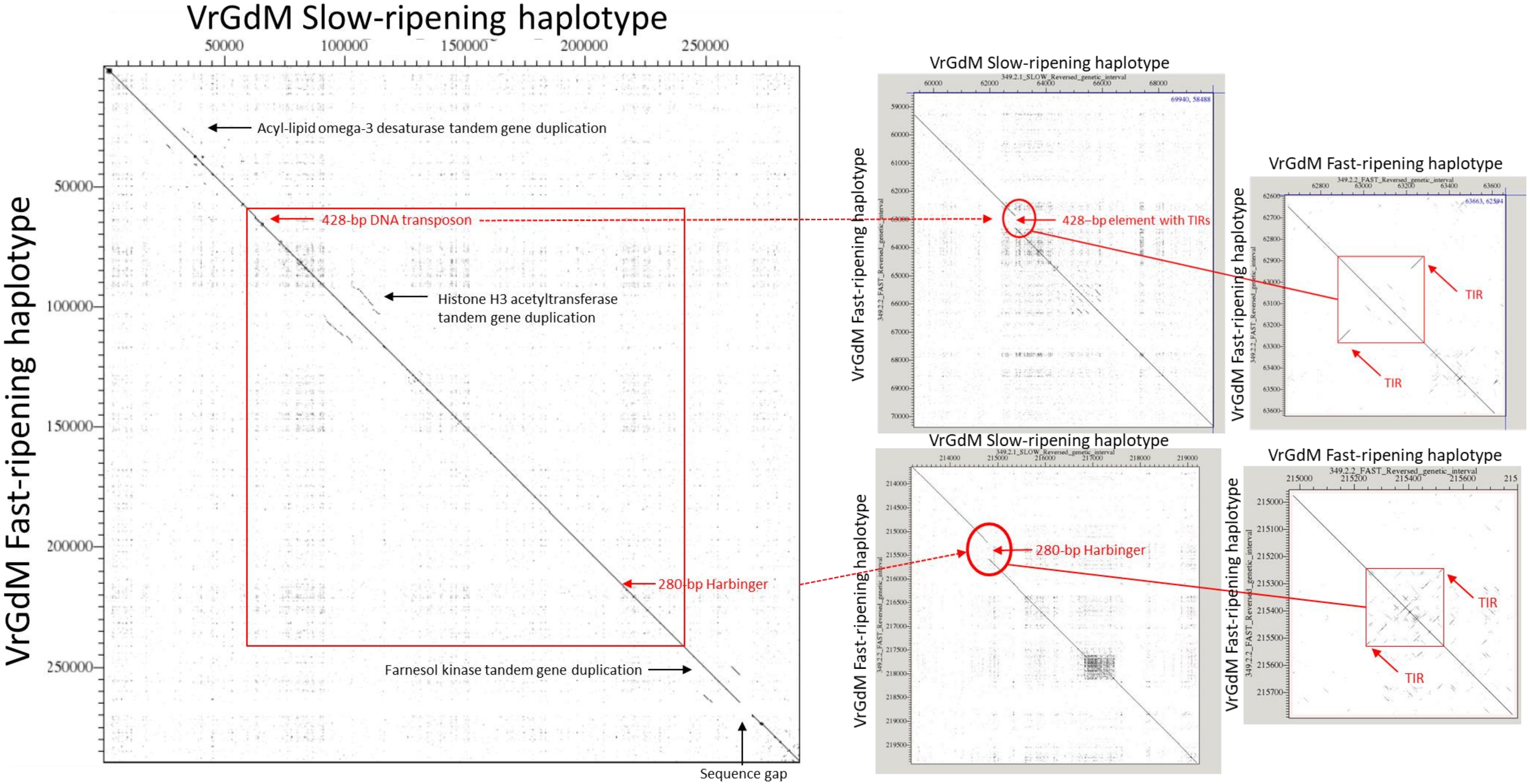


**Supplementary Figure S22 - Dotplot comparison of the slow-ripening haplotype (horizontal) and the fast-ripening haplotype (vertical) assembled from VrGdM.** The red rectangle delimits the *RIPENING SPEED* QTL peak. The magnified views show the insertion sites of the DNA transposons in the fast-ripening haplotype and their tandem inverted repeats (TIRs). Supporting evidence for the heterozygous insertions in VrGdM is provided by the alignment of VrGdM short and long reads with the VrGdM-derived haplotype assembled from 'Kober 5BB'. See Integrative Genome Viewer (Falginella et al 2024). Blastn searches of the flanking sequences on each side of the insertion site indicates that the 428-bp DNA transposon is present in the same genomic position in other *V. riparia* haplotypes, in *V. acerifolia*, in *V. rupestris* and in the haplotype that *V. rupestris* has donated to the interspecific rootstock 110R. It is absent from *V.*

*mustangensis*, *V. monticola*, *V. arizonica*, *V. berlandieri*, *V. aestivalis* and *V. cinerea*, from the haplotypes that *V. berlandieri* has donated to 110R and Kober 5BB, from *Muscadinia rotundifolia* and from *V. vinifera* (Zou et al 2020, grapegenomics.com). The same analysis indicated that the Harbinger insertion is private to VrGdM among the accessions available at grapegenomics.com or sequenced by Zou et al (2020).

Falginella et al (2024) Grape ripening speed slowed down using natural variation. *figshare* 10.6084/m9.figshare.25991668

grapegenomics.com. Available at: https://www.grapegenomics.com/

Zou et al (2020) Haplotyping the *Vitis* collinear core genome with rhAmpSeq improves marker transferability in a diverse genus. Nature Communications 11:413

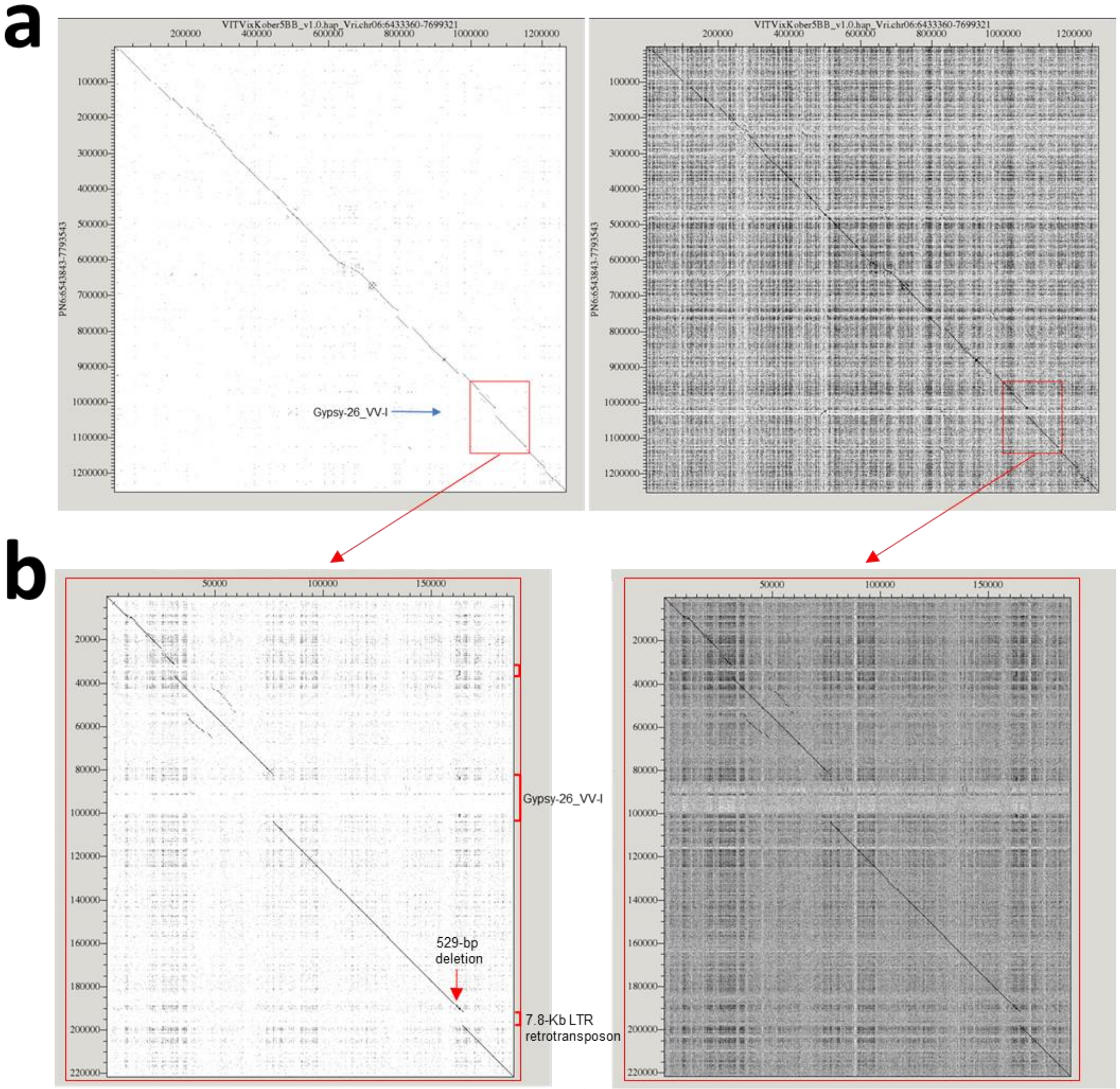


**Supplementary Figure S23 - Dotplot comparison in the *RIPENING SPEED* QTL confidence interval (a) and in the QTL peak (b) of the gapless slow-ripening haplotype assembled from 'Kober 5BB' (horizontal, from PacBio HiFi reads) and a gapless *V. vinifera* haplotype assembled from 'PN40024' (vertical, from PacBio HiFi reads).** The red rectangle delimits the *RIPENING SPEED* QTL peak in **a**, which is magnified in **b**. The two plots in the same panel show two different grayscale ramp settings to emphasize coarse-grained (left plot) or fine-grained (right plot) sequence identity.

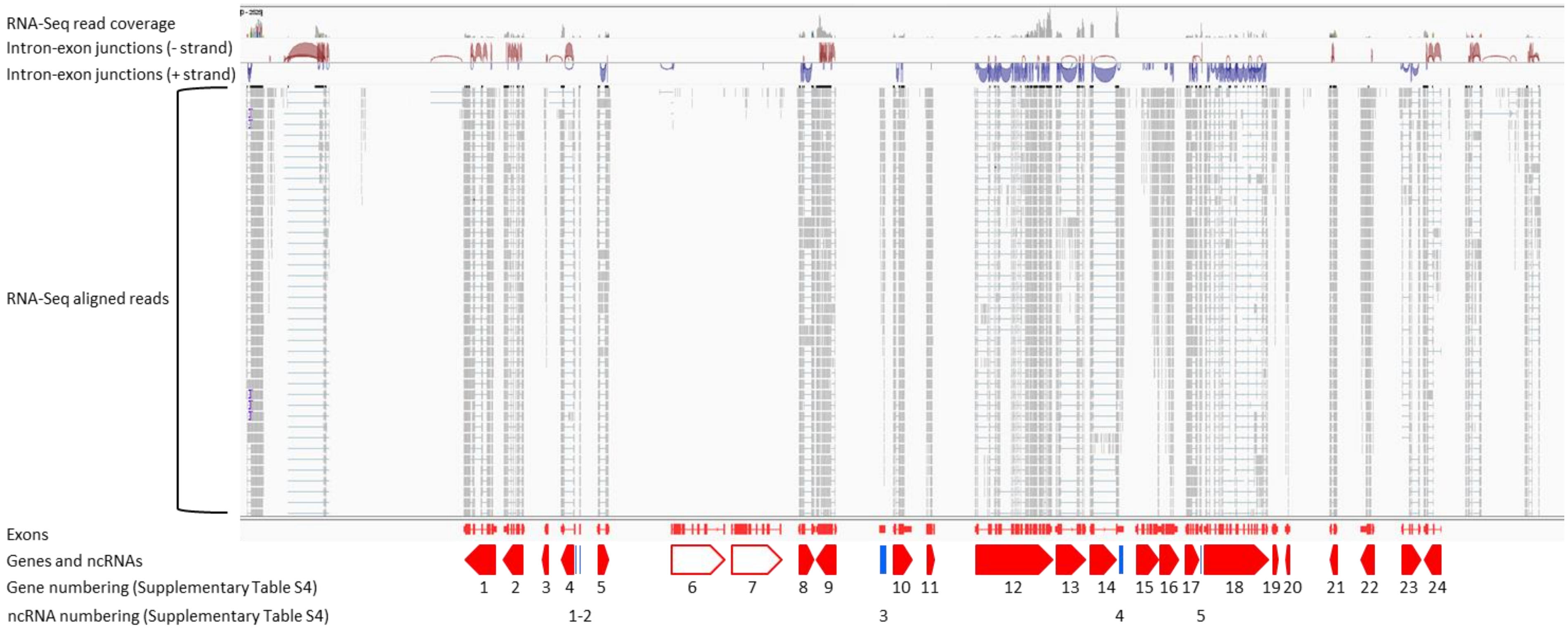


**Supplementary Figure S24 - Gene spacing in the slow-ripening haplotype.** Red polygons represent genes. The arrowed shape of the polygon points towards the 3'end of the gene. Arrowed shapes pointing to the left represent genes that are transcribed from the negative strand. Arrowed shapes pointing to the right represent genes that are transcribed from the positive strand. Solid symbols represent expressed genes. Open symbols represent genes without evidence of transcription from pooled leaf and root RNA-Seq libraries. Blue rectangles represent ncRNA loci with evidence of transcription. Gene and ncRNA symbols are identified by numbers, which are cross-referenced to gene IDs and functional annotation in **Supplementary Table S4**. Tracks of RNA coverage and reads and predicted intron-exon junction are derived from the alignment of RNA-Seq data in leaves and roots of *V. riparia* 'Gloire de Montpellier' (Girollet et al 2019) with the slow-ripening haplotype assembled from 'Kober 5BB' (Minio et al 2022).

Girollet et al (2019) De novo phased assembly of the *Vitis riparia* grape genome. Scientific Data 6:127
Minio et al (2022) HiFi chromosome-scale diploid assemblies of the grape rootstocks 110R, Kober 5BB, and 101-14 Mgt. Scientific Data 9:660

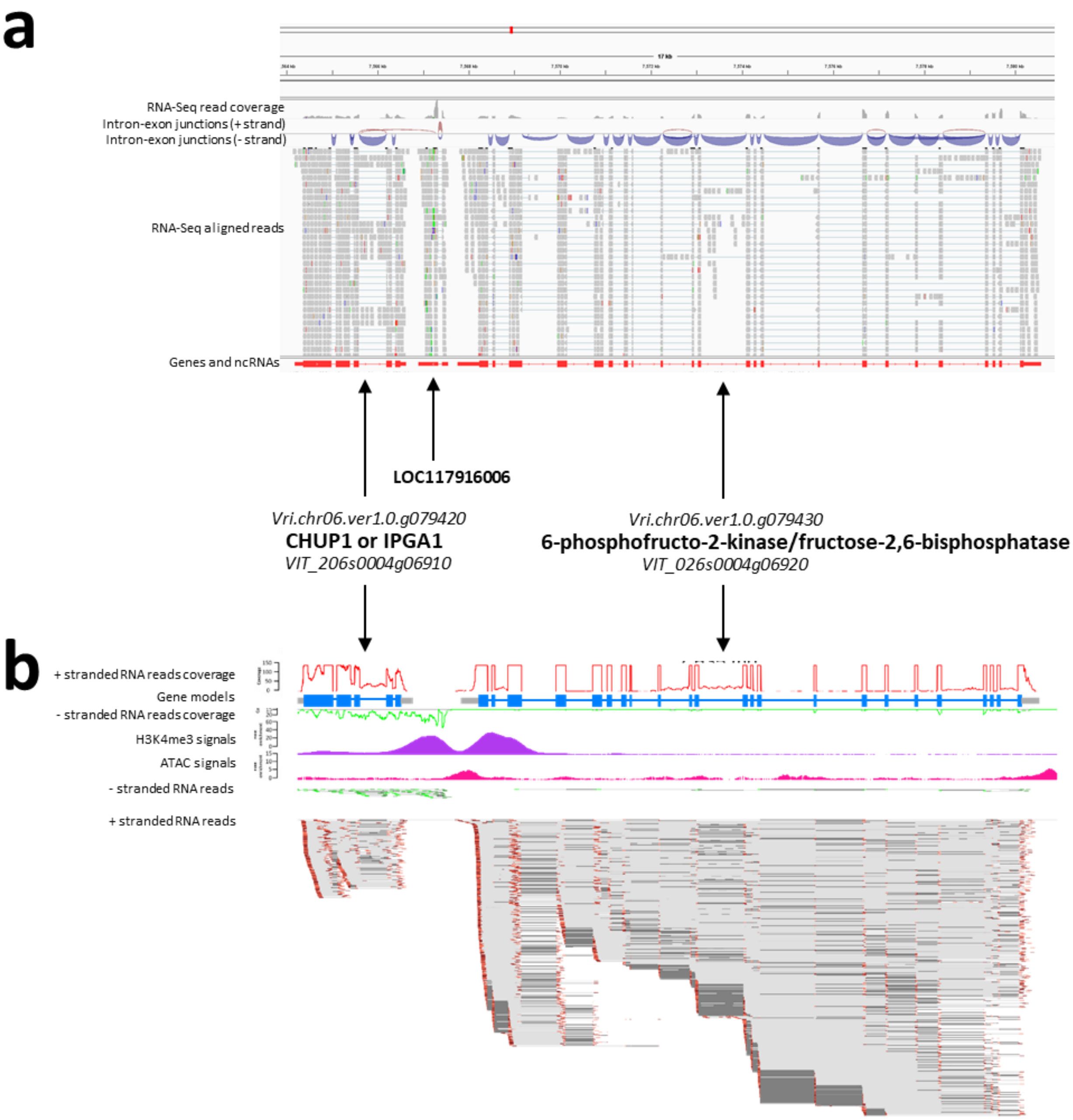


**Supplementary Figure S25 - Prediction and expression profiles of a ncRNA locus (LOC117916006) and the adjacent genes encoding a Chloroplast Outer Envelope Protein (CHUP1) or INCREASED PETAL GROWTH ANISOTROPY 1 (IPGA1) and a 6-phosphofructo-2-kinase/fructose-2,6-bisphosphatase. a** Expression profiles are derived from the alignment of RNA-Seq data in leaves and roots of *V. riparia* 'Gloire de Montpellier' (Girollet et al 2019) with the slow-ripening haplotype assembled from 'Kober 5BB' (Minio et al 2022). **b** Gene view of mRNA coverage in *V. vinifera* 'Cabernet Franc' leaves from stranded RNA-Seq libraries (BioProject number PRJNA373967, Magris et al 2021) aligned with the *V. vinifera* 'PN40024' reference genome 12Xv0 (Jaillon et al 2007). Coverage of mRNA transcribed from the positive and the negative strand is reported with respect to the predicted intron-exon structure. Coverage values greater than 150 were levelled to 150. Blue boxes indicate CDS, grey boxes indicate UTRs, grey lines indicate introns. Read alignments on the positive (red paired-end reads) and the negative strand (green paired-end reads) are reported. For each

RNA fragment, R1 reads are dark coloured, R2 reads are light coloured. Light grey connectors connect paired-end reads. Dark grey connectors connect split reads. H3K4me3 and ATAC signals are derived from data deposited under the BioProject number PRJNA643441 (Schwope et al (2021). The active histone mark H3K4me3 at the LOC117916006 transcription start site and the negative stranded RNA reads indicate that LOC117916006 is transcribed in antisense direction with respect to *VIT_206s0004g06910*. Sense and antisense transcripts from the two transcriptional units largely overlap.


Girollet et al (2019) De novo phased assembly of the *Vitis riparia* grape genome. Scientific Data 6:127

Magris et al (2021) The genomes of 204 *Vitis vinifera* accessions reveal the origin of European wine grapes. Nature Communications 12:7240

Minio et al (2022) HiFi chromosome-scale diploid assemblies of the grape rootstocks 110R, Kober 5BB, and 101-14 Mgt. Scientific Data 9:660

Schwope et al (2021) Open chromatin in grapevine marks candidate CREs and with other chromatin features correlates with gene expression. Plant Journal 107:1631-1647

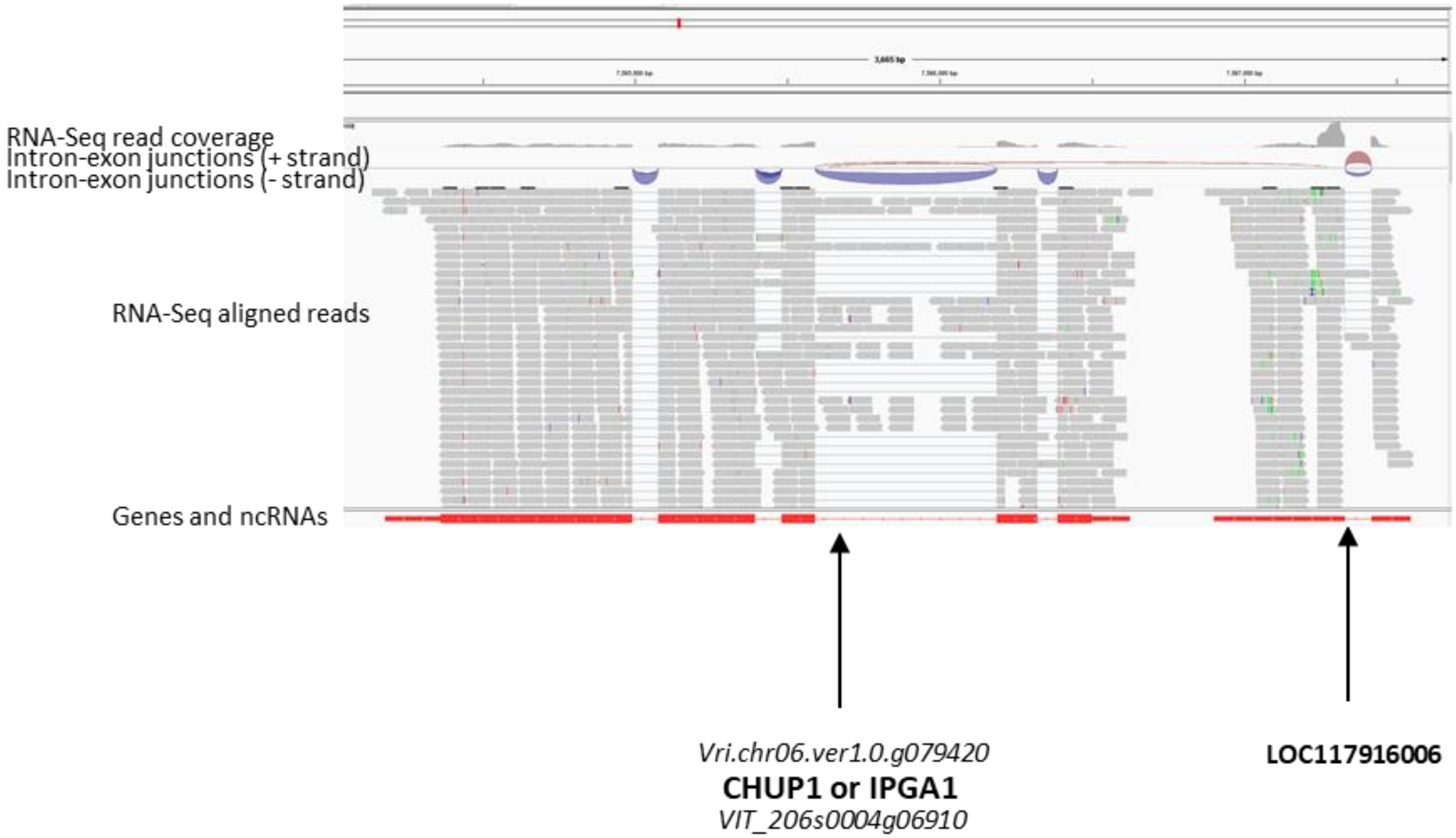


**Supplementary Figure S26 - Magnified view of the prediction and expression profiles of a Chloroplast Outer Envelope Protein (CHUP1) or INCREASED PETAL GROWTH ANISOTROPY 1 (IPGA1) coding gene in *V. vinifera* (gene model *VIT_206s0004g06910*) and in *V. riparia* 'Gloire de Montpellier' (gene model *Vri.chr06.ver1.0.g079420*) and the downstream LOC117916006 ncRNA.** Expression profiles are derived from the alignment of RNA-Seq data in leaves and roots of *V. riparia* ‘Gloire de Montpellier’ (Girollet et al 2019) with the slow-ripening haplotype assembled from ‘Kober 5BB’ (Minio et al 2022).

Girollet et al (2019) De novo phased assembly of the *Vitis riparia* grape genome. Scientific Data 6:127

Minio et al (2022) HiFi chromosome-scale diploid assemblies of the grape rootstocks 110R, Kober 5BB, and 101-14 Mgt. Scientific Data 9:660

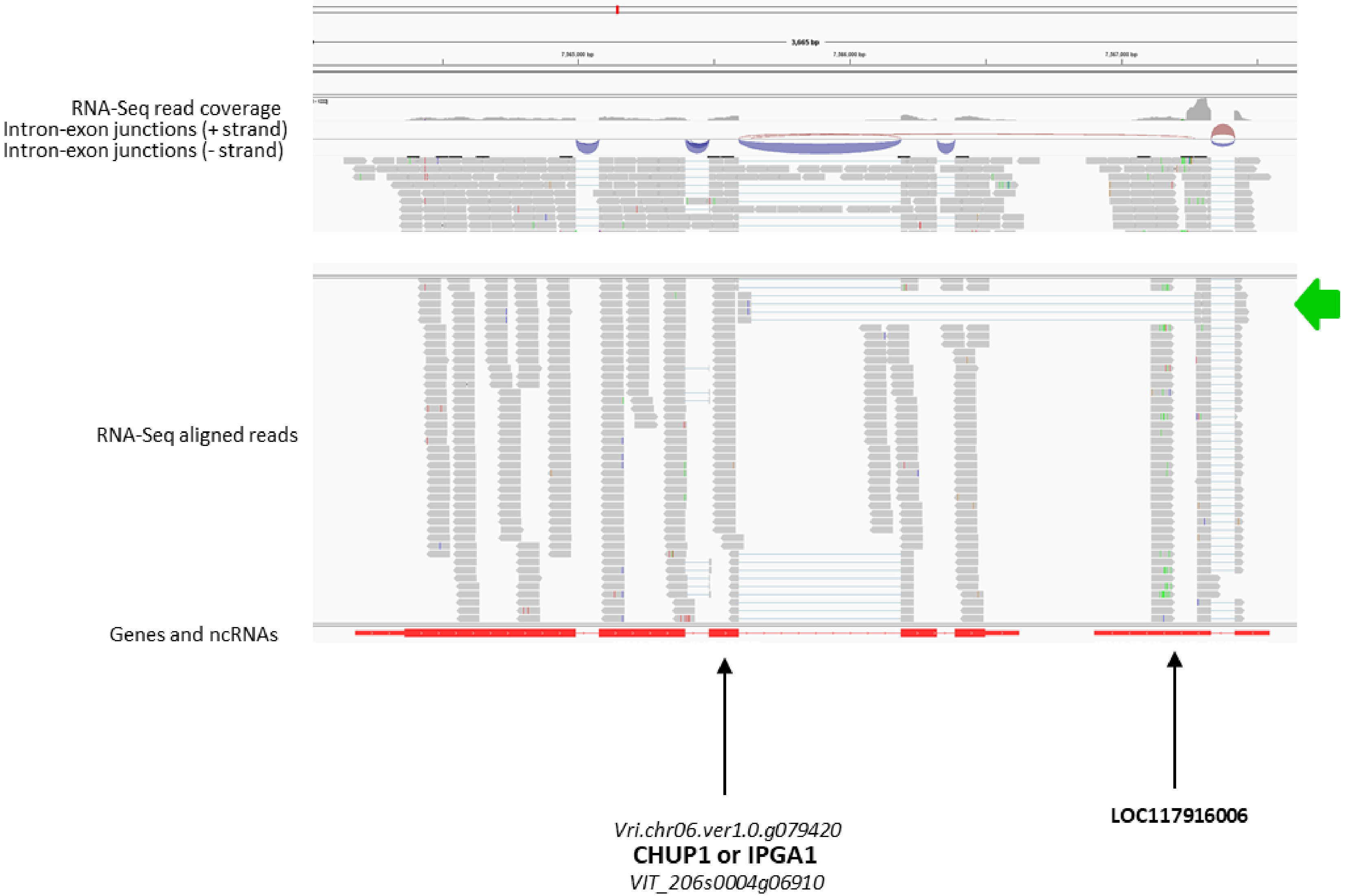


**Supplementary Figure S27 - Prediction and expression profiles of a Chloroplast Outer Envelope Protein (CHUP1) or INCREASED PETAL GROWTH ANISOTROPY 1 (IPGA1) coding gene in *V. riparia* 'Gloire de Montpellier' (gene model *Vri.chr06.ver1.0.g079420*) and the downstream LOC117916006 ncRNA.** Expression profiles are derived from the alignment of RNA-Seq data in leaves and roots of *V.*

*riparia* ‘Gloire de Montpellier’ (Girollet et al 2019) with the slow-ripening haplotype assembled from ‘Kober 5BB’ (Minio et al 2022). The green arrow indicates spliced LOC117916006 transcripts that overlap with *Vri.chr06.ver1.0.g079420* transcripts.

Girollet et al (2019) De novo phased assembly of the *Vitis riparia* grape genome. Scientific Data 6:127
Minio et al (2022) HiFi chromosome-scale diploid assemblies of the grape rootstocks 110R, Kober 5BB, and 101-14 Mgt. Scientific Data 9:660

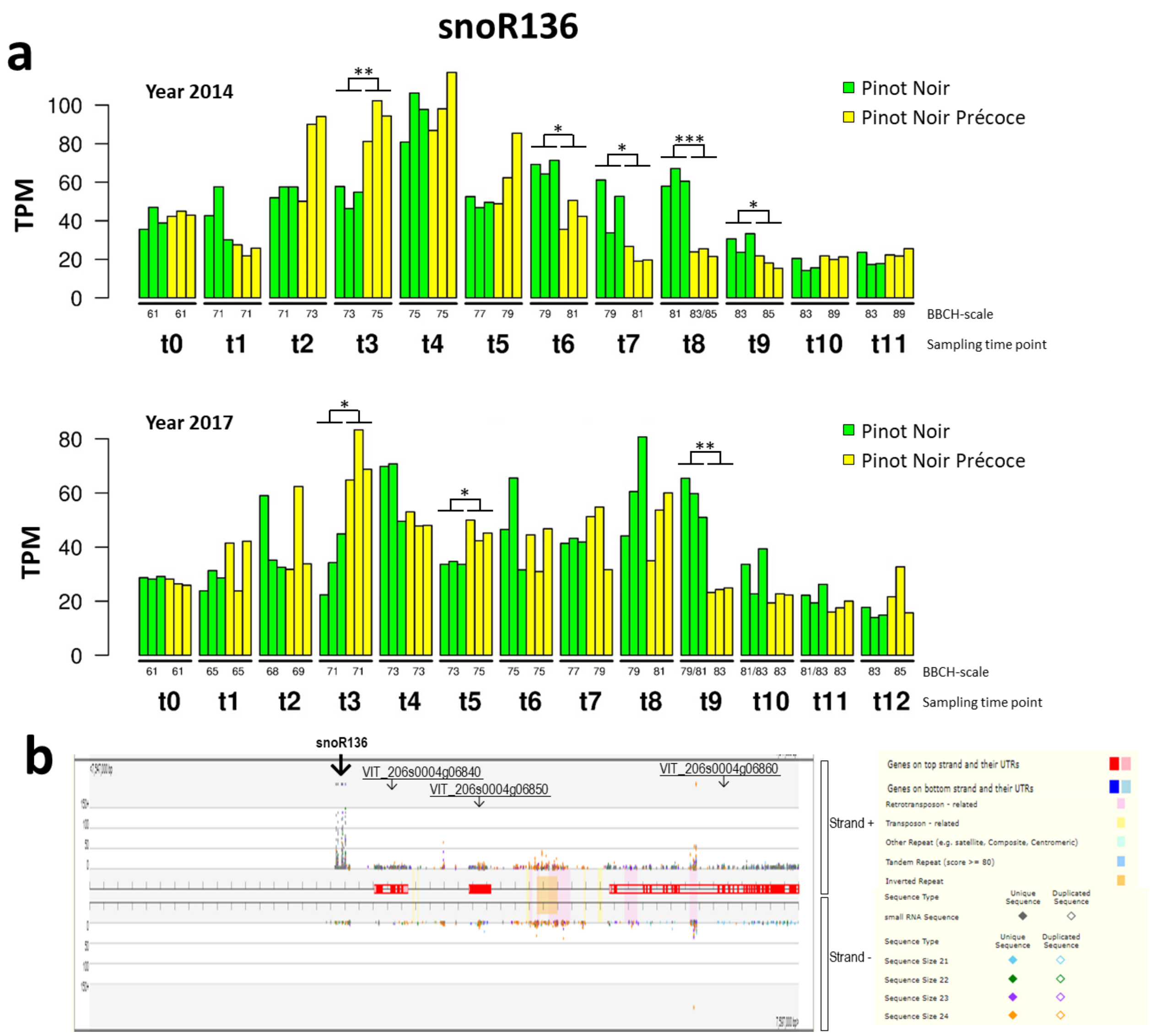


**Supplementary Figure S28 - Expression profile of a small nucleolar RNA transcript (snoR136) in berries of 'Pinot Noir' isogenic lines, one of which ('Pinot Noir Précoce') differs from the wild-type for the ripening pattern, and small RNAs produced from the snoR136 transcripts in 'Cabernet Sauvignon' and 'Sangiovese' berries. a** Sampling time points (t*n*) and developmental/ripening stages (BBCH scale) are expressed as reported by Theine et al (2021). Each bar represents a biological replicate. Significant differences between means of biological replicates were determined using a Student's t-test and $log_{10}$-transformed TPM values between isogenic lines at the same time point are indicated by asterisks (* <0.05, ** <0.01, ***<0.001). **b** Read counts of small RNAs on positive and negative strands. The colour of the dot indicates read size as reported in the legend. Small RNA data corresponded to GEO Accession Numbers GSM2279692-9739 (Paim Pinto et al 2016) and graphic visualization was obtained from the https://mpss.danforthcenter.org/10 (Nakano et al 2020). Small RNA counts from the snoR136 transcripts are shown relative to those produced from nearby coding sequences of the genes *VIT_206s0004g6840*, *VIT_206s0004g6850* and *VIT_206s0004g6860* as well as from intronic and intergenic transposable elements.

Nakano et al (2020) Next-generation sequence databases: RNA and genomic informatics resources for plants. Plant Physiology 182:136-146

Paim Pinto et al (2016) The influence of genotype and environment on small RNA profiles in grapevine berry. Frontiers in Plant Science 7:1459

Theine et al. (2021) Transcriptomic analysis of temporal shifts in berry development between two grapevine cultivars of the Pinot family reveals potential genes controlling ripening time. BMC Plant Biology 21:237

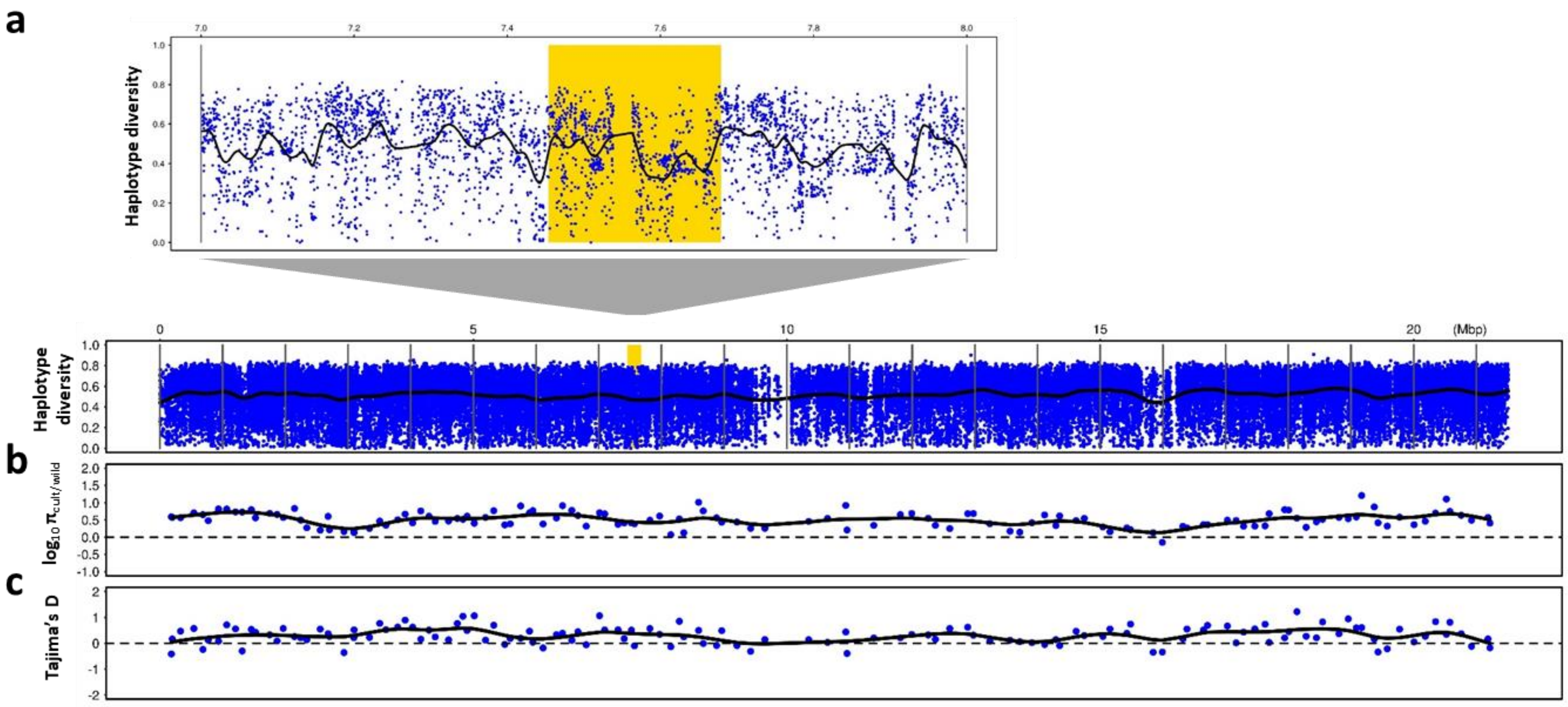


**Supplementary Figure S29 - Genetic diversity and search for selection signals in *V. vinifera* across Chr6. a** Haplotype diversity in cultivars with (above) magnification of the interval comprised between 7-8 Mbp containing the *RIPENING SPEED* QTL peak highlighted in amber background. Haplotype diversity was calculated in blocks of five consecutive variant sites and plotted as the average of 50 consecutive blocks (blue dots). **b** Relative nucleotide diversity in cultivars with respect to the wild progenitor. **c** Tajima's D in cultivars. Blue dots in b and c represent average values in genomic windows containing 100 Kb of non-repetitive DNA. Black lines in all panels represent cubic smoothing splines of values. Original data for generating these plots were obtained from Magris et al (2021) using the methods described therein. Chromosomal coordinates refer to the *V. vinifera* 'PN40024' reference genome 12Xv0 (Jaillon et al 2007).

Jaillon et al (2007) The grapevine genome sequence suggests ancestral hexaploidization in major angiosperm phyla. Nature 449:463-467
Magris et al (2021) The genomes of 204 *Vitis vinifera* accessions reveal the origin of European wine grapes. Nature Communications 12: 7240

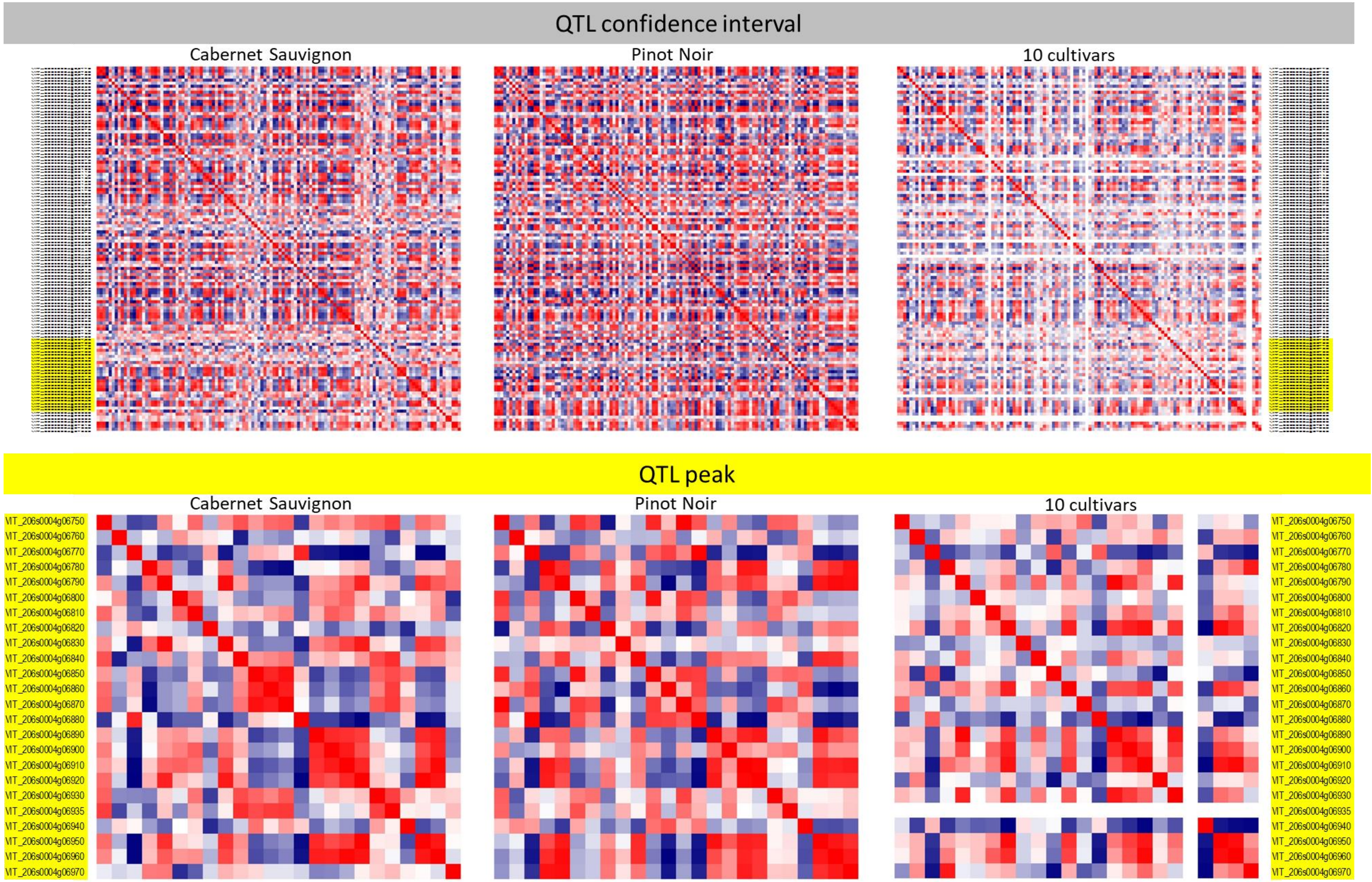


**Supplementary Figure S30 - Correlation matrix of gene expression in developing and ripening berries of ‘Cabernet Sauvignon’ and ‘Pinot Noir’ with short sampling intervals (Fasoli et al 2018) and 10 cultivars sampled at 4 key stages (Massonnet et al 2017) among genes in the *RIPENING SPEED* QTL confidence interval and peak.** Pearson’s correlation of transcript levels of genes ordered by chromosomal location. The red-to-blue heat map indicate co-regulated (red) and anti-regulated (blue) gene pairs. Gene IDs correspond to the V2.1 gene prediction (Vitulo et al 2014) of the *V. vinifera* ‘PN40024’ reference genome 12Xv0 (Jaillon et al 2007). Genes that were not predicted in the V1 gene prediction used by Massonnet et al (2017) are blank in the matrix.

Fasoli et al (2018) Timing and order of the molecular events marking the onset of berry ripening in grapevine. Plant Physiology 178:1187-1206

Jaillon et al (2007) The grapevine genome sequence suggests ancestral hexaploidization in major angiosperm phyla. Nature 449:463-467

Massonnet et al (2017) Ripening transcriptomic program in red and white grapevine varieties correlates with berry skin anthocyanin accumulation. Plant Physiology 174:2376-2396

Vitulo et al (2014) A deep survey of alternative splicing in grape reveals changes in the splicing machinery related to tissue, stress condition and genotype. BMC Plant Biology 14:99

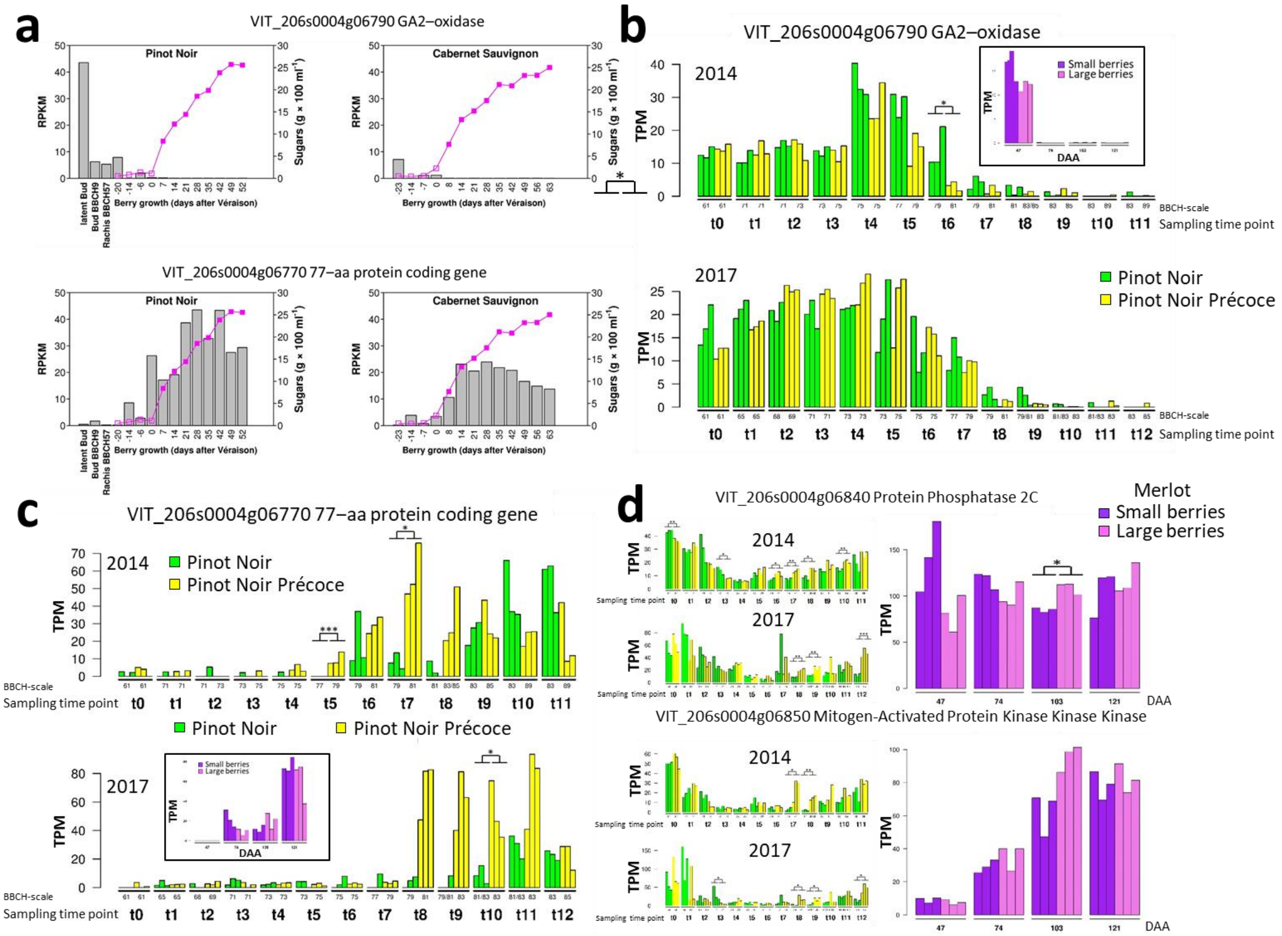


**Supplementary Figure S31 – Contrasting expression patterns in *V. vinifera* developing and ripening berries of two positional candidate genes either up-regulated or down-regulated with ripening and two genes putatively involved in hormonal, developmental or environmental signal transduction. a** Time-course of gene expression at weekly intervals in 'Pinot Noir' and 'Cabernet Sauvignon'. Data is derived from Gene Expression Omnibus accession number GSE98923 and Fasoli et al (2018). The progression of berry growth and ripening in 'Pinot Noir' and 'Cabernet Sauvignon' is illustrated by the curve of sugars concentration (shown on the secondary y-axis) at preveraison (empty squares) and postveraison (full squares) stages. Data at three stages of inflorescence development within latent and bursting buds and just prior to flowering in 'Pinot Noir' is derived from Rossmann et al (2020). **b-d** Gene expression in 'Pinot Noir' isogenic lines differing for ripening patterns. Data is derived from European Nucleotide Archive (ENA) accession numbers PRJEB39261-4 and Theine et al (2021). Sampling time points (tn) and developmental/ripening stages (BBCH scale) are expressed as reported by Theine et al (2021), sampling was repeated over two years (2014 and 2017). Each bar represents a biological replicate. Significant differences between means of biological replicates were determined using a Student's t-test and $\log_{10}$-transformed TPM values between time points are indicated by asterisks (* <0.05, ** <0.01, ***<0.001). Gene expression in the epicarp of 'Merlot' ripening berries of different sizes at 4 developmental stages (DAA 47, 74, 103, 121). Data is derived from NCBI Sequence Read Archive BioProject PRJNA316157 and Wong et al (2016). Each bar represents a biological replicate. Significant differences between means of biological replicates were determined using a Student's t-test and $\log_{10}$-transformed TPM values between time points are indicated by asterisks (* <0.05, ** <0.01, ***<0.001).

Rossmann et al (2020) Mutations in the miR396 binding site of the growth-regulating factor gene VvGRF4 modulate inflorescence architecture in grapevine. Plant Journal 101:1234-1248

Fasoli et al (2018) Timing and order of the molecular events marking the onset of berry ripening in grapevine. Plant Physiology 178:1187-1206

Theine et al. (2021) Transcriptomic analysis of temporal shifts in berry development between two grapevine cultivars of the Pinot family reveals potential genes controlling ripening time. BMC Plant Biology 21:237

Wong et al (2016) Combined physiological, transcriptome, and cis-regulatory element analyses indicate that key aspects of ripening, metabolism, and transcriptional program in grapes (*Vitis vinifera* L.) are differentially modulated accordingly to fruit size. BMC Genomics 17:416

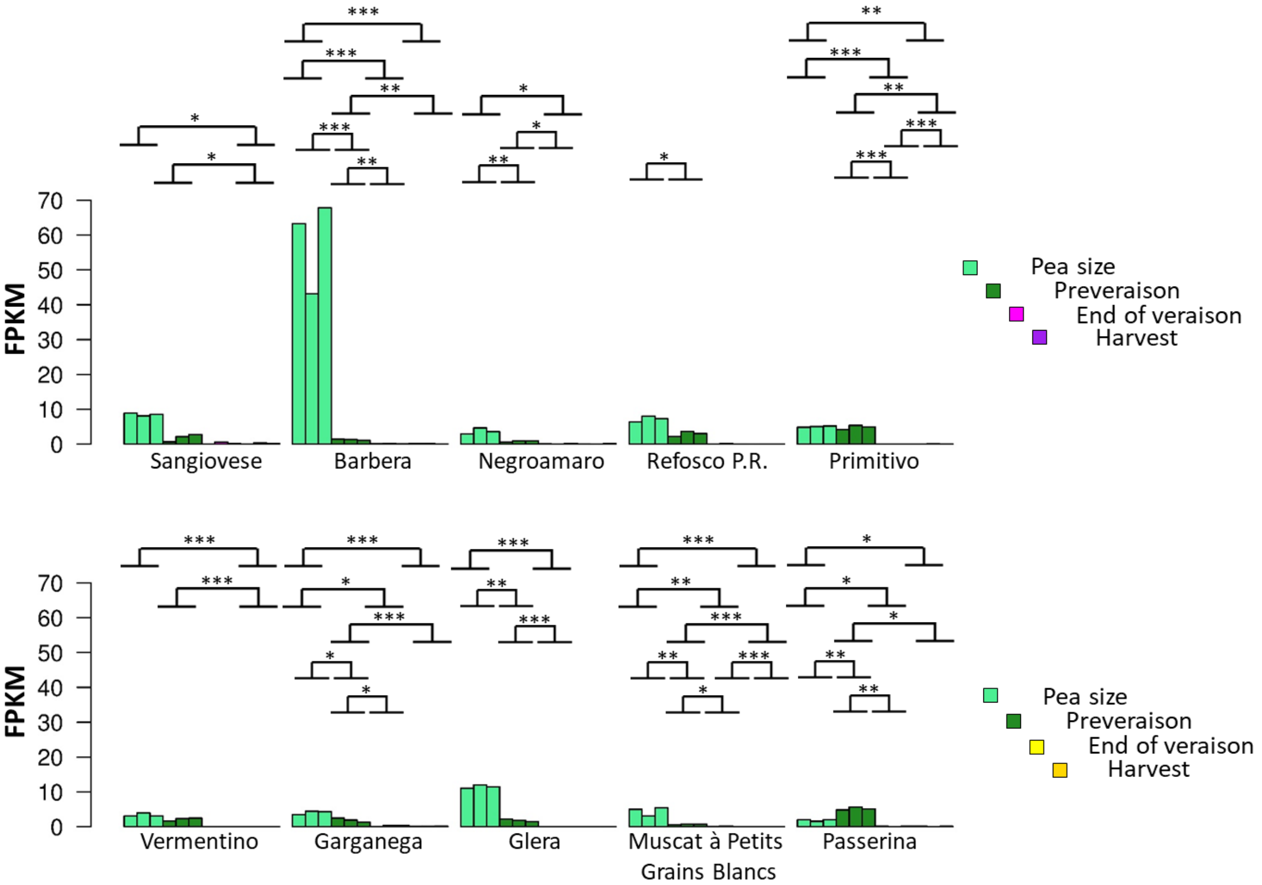


**Supplementary Figure S32 - Expression of *VIT_206s0004g06790* encoding a GA2-oxidase in berries at four developmental stages (three replicates per stage) in five red-skinned (top) and five white-skinned varieties (bottom).** Y-axes report FPKM. The initial two stages correspond to the first phase of green berry growth. The following two stages correspond to the second phase of berry growth that accompanies ripening. All y-axes are scaled a maximum value of 70 FPKM. Soluble solids concentrations of each sample are reported in Massonnet et al (2017). Significant differences between means of biological replicates were determined using a Student's t-test and $log_{10}$-transformed FPKM values between time points are indicated by asterisks (* <0.05, ** <0.01, ***<0.001).

Massonnet et al (2017) Ripening transcriptomic program in red and white grapevine varieties correlates with berry skin anthocyanin accumulation. Plant Physiology 174:2376-2396

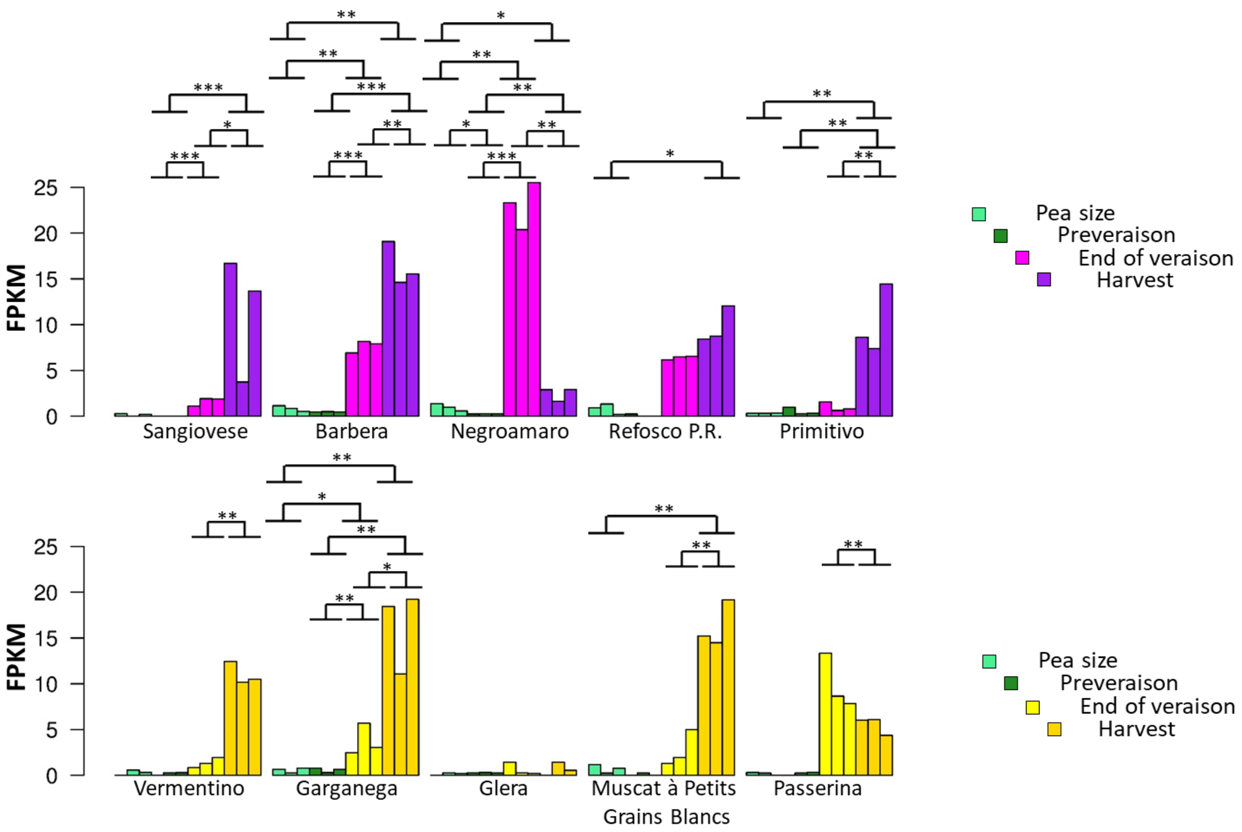


**Supplementary Figure S33 - *VIT_206s0004g06770* expression in berries at four developmental stages (three replicates per stage) in five red-skinned (top) and five white-skinned varieties (bottom).** Y-axes report FPKM. The initial two stages correspond to the first phase of green berry growth. The following two stages correspond to the second phase of berry growth that accompanies ripening. All y-axes are scaled a maximum value of 25 FPKM. Soluble solids concentrations of each sample are reported in Massonnet et al (2017). Significant differences between means of biological replicates were determined using a Student's t-test and $log_{10}$-transformed FPKM values between time points are indicated by asterisks (* <0.05, ** <0.01, ***<0.001).

Massonnet et al (2017) Ripening transcriptomic program in red and white grapevine varieties correlates with berry skin anthocyanin accumulation. Plant Physiology 174:2376-2396

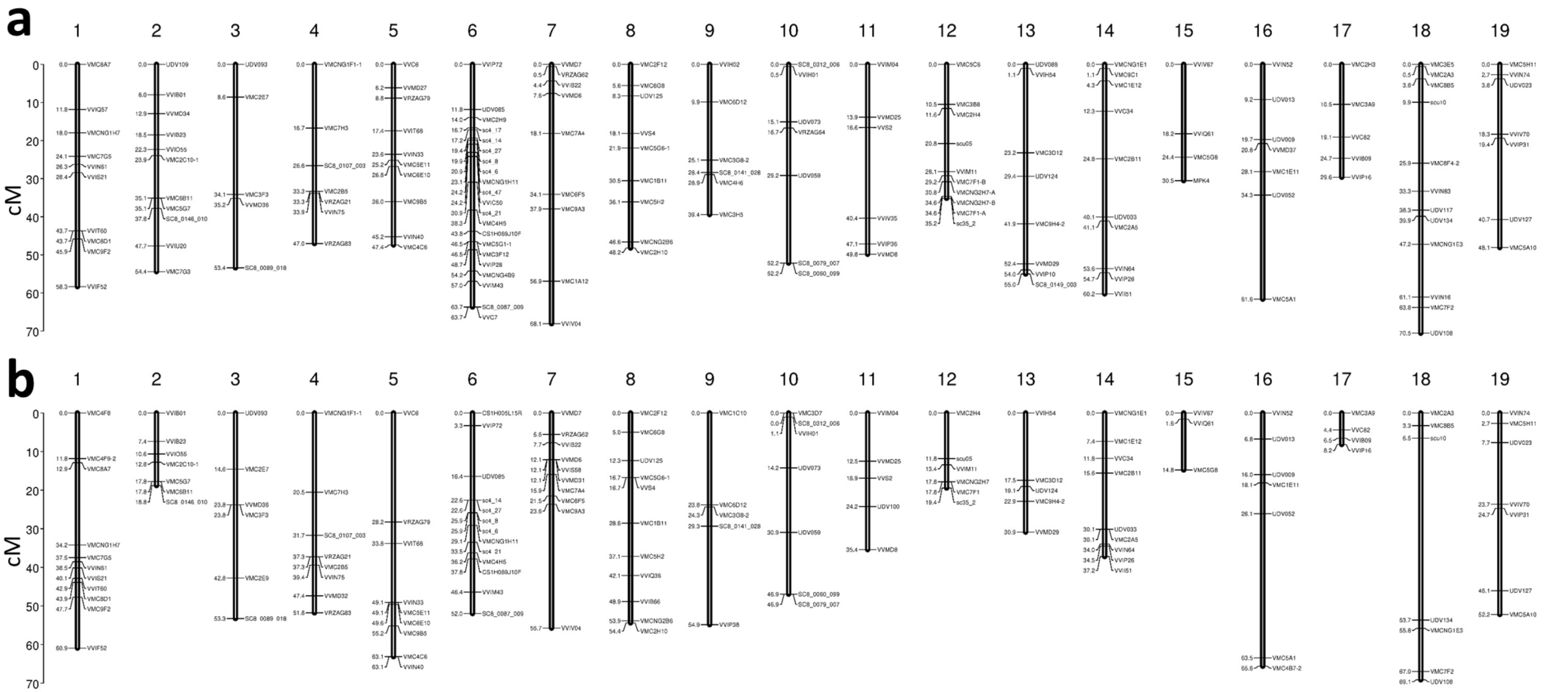


**Supplementary Figure S34 - Genetic maps in *V. riparia* 'Gloire de Montpellier' (a) and *V. vinifera* Chardonnay (b).** Genetic distances are expressed in cM. Linkage groups are ordered and oriented according to the numbering and orientation of the corresponding chromosomes in the *V. vinifera* 'PN40024' reference genome 12Xv0 (Jaillon et al 2007).

Jaillon et al (2007) The grapevine genome sequence suggests ancestral hexaploidization in major angiosperm phyla. Nature 449:463-467

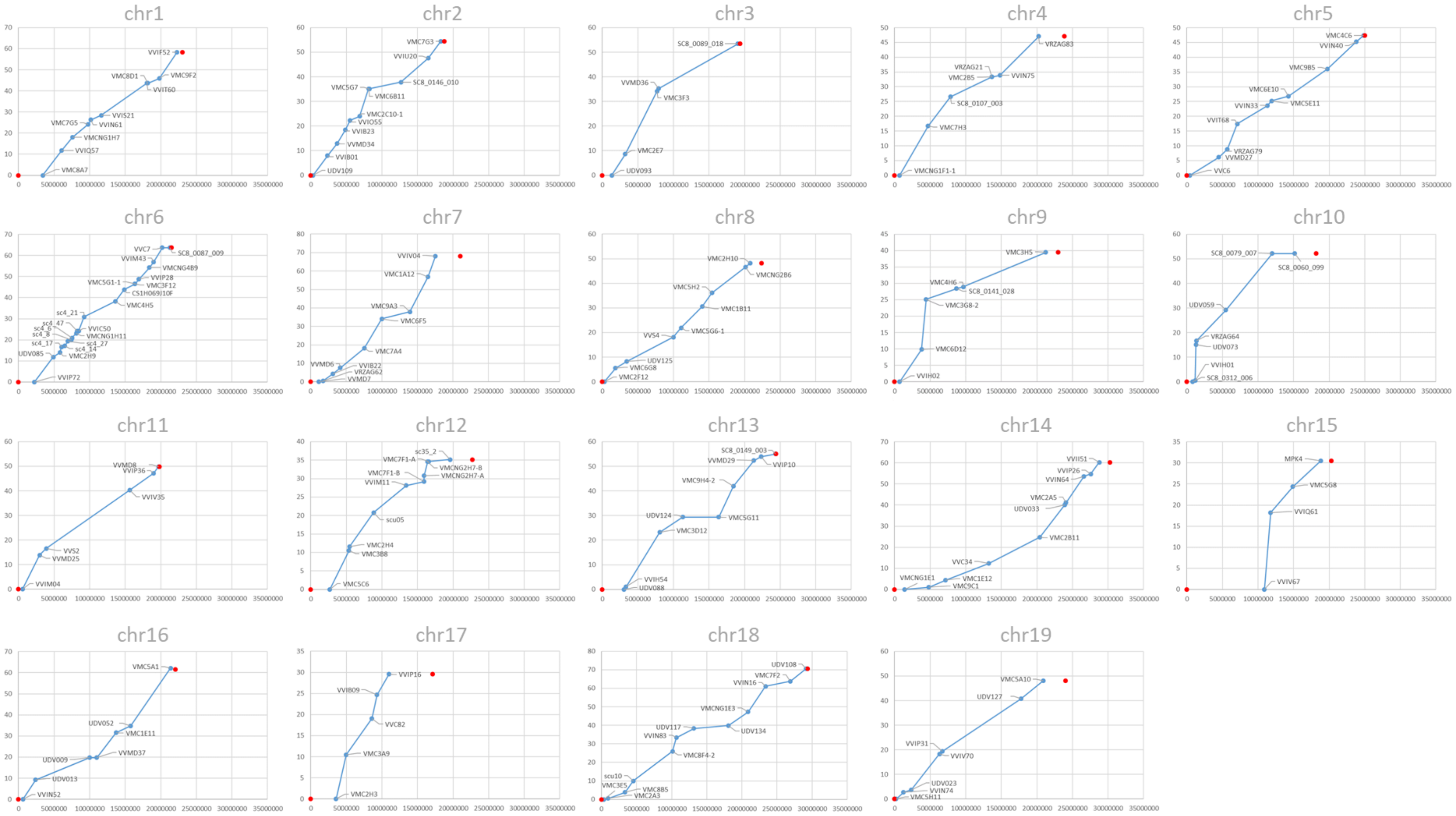


**Supplementary Figure S35 - Physical coverage of the *V. riparia* ‘Gloire de Montpellier’ genetic map.** Relation of physical distance and genetic recombination. X-axis indicates the chromosomal coordinates of the markers in the *V. vinifera* ‘PN40024’ reference genome 12Xv0 (Jaillon et al 2007). Y-axis indicates cumulative genetic distance in Kosambi cM. Red dots indicate the initial and final coordinates of the chromosome.

Jaillon et al (2007) The grapevine genome sequence suggests ancestral hexaploidization in major angiosperm phyla. Nature 449:463-467

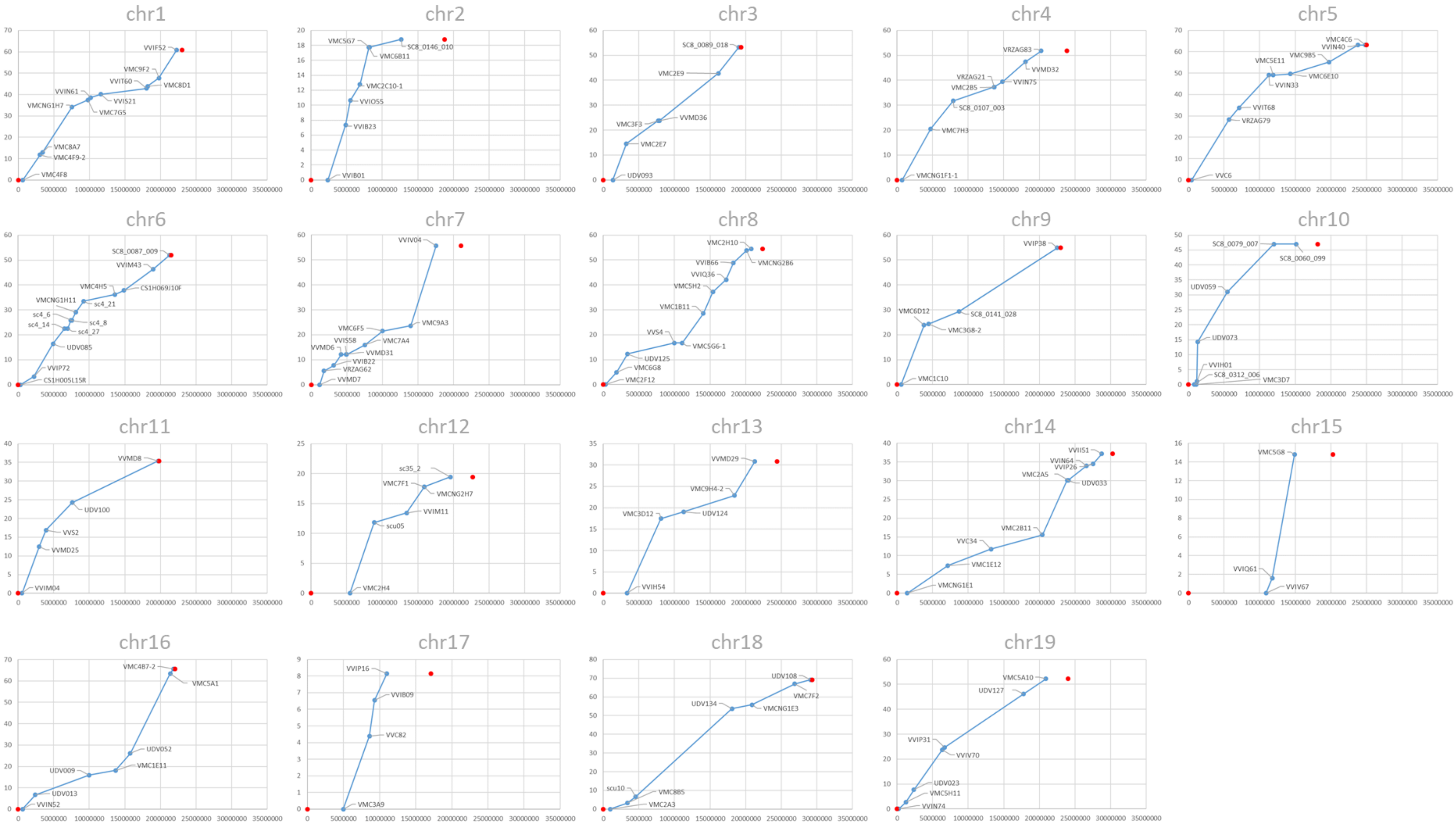


**Supplementary Figure S36 - Physical coverage of the *V. vinifera* 'Chardonnay' genetic map.** Relation of physical distance and genetic recombination. X-axis indicates the chromosomal coordinates of the markers in the *V. vinifera* 'PN40024' reference genome 12Xv0 (Jaillon et al 2007). Y-axis indicates cumulative genetic distance in Kosambi cM. Red dots indicate the initial and final coordinates of the chromosome.

Jaillon et al (2007) The grapevine genome sequence suggests ancestral hexaploidization in major angiosperm phyla. Nature 449:463-467
